# CarpeDeam: A *De Novo* Metagenome Assembler for Heavily Damaged Ancient Datasets

**DOI:** 10.1101/2024.08.09.607291

**Authors:** Louis Kraft, Johannes Söding, Martin Steinegger, Annika Jochheim, Peter Wad Sackett, Antonio Fernandez-Guerra, Gabriel Renaud

## Abstract

*De novo* assembly of ancient metagenomic datasets is a challenging task. Ultra-short fragment size and characteristic postmortem damage patterns of sequenced ancient DNA molecules leave current tools ill-equipped for ideal assembly. We present CarpeDeam, a novel damage-aware *de novo* assembler designed specifically for ancient metagenomic samples. Utilizing maximum-likelihood frameworks that integrate sample-specific damage patterns, CarpeDeam demonstrates improved recovery of longer continuous sequences and protein sequences in many simulated and empirical datasets compared to existing assemblers. As a pioneering ancient metagenome assembler, CarpeDeam opens the door for new opportunities in functional and taxonomic analyses of ancient microbial communities.

## 2 Background

DNA recovered from ancient organisms, termed ancient DNA (aDNA), has transformed evolutionary science. Besides the exploration of ancestral human populations, aDNA enables the reconstruction of past environments covering eukaryotic species diversity and microbial community dynamics [1, 2, 3, 4]. However, the computational analysis of aDNA is challenging [5, 6]. Two primary processes lead to the breakdown and chemical alteration of aDNA. Hydrolytic mechanisms cause the degradation of DNA molecules and deamination converts cytosines into uracil molecules. These are misinterpreted as putative base substitutions during DNA sequencing [7]. This phenomenon, known as “damage”, occurs predominantly at the ends of aDNA fragments, and the rate of occurrence of damage at each fragment position can be quantified [8, 9, 10, 11]. As a result, aDNA fragments show elevated frequencies of C→T substitutions at the 5^′^ end and G→A substitutions at the 3^′^ end, respectively [12]. Along with the short fragment size, damage profiles are commonly used for aDNA authentication [13, 14, 15, 16]. However, the ambiguity introduced by deaminated bases hampers downstream analysis such as read mapping or genome assembly [17, 18, 19].

Metagenomes represent the collective genetic material from various species within a sample, derived from environments such as the human gut or soil, including bacteria, archaea, viruses, and eukaryotes [20, 21]. The field has expanded with the advent of massive parallel sequencing, leading to a deeper understanding of microbial diversity and functional profiling on Earth [22, 23, 24]. However, analysing metagenomic datasets relies on dedicated tools as the data is inherently complex due to high species diversity and varying abundance profiles, and the choice of tools depends on the specific research goals [25]. There are two main approaches that follow different objectives when analysing metagenomic data: taxonomic classification and functional annotation [26].

Taxonomic classification categorizes genomic sequences to determine microbial composition, thus offering an overview of the taxonomic diversity [27]. Numerous tools have been developed, utilizing both read-based and assembly-based approaches. Read-based methods employ various strategies such as marker gene analysis, database alignment, or *k*-mer-based techniques [28, 27, 29, 30]. Assembly-based methods, which work with either assembled contigs or long reads, potentially increase classification accuracy. These approaches rely on tailored data structures or alignment techniques to efficiently process large numbers of sequences [31, 32, 33, 34]. Notably, taxonomic classification has been widely applied to ancient metagenomes [35, 36, 37, 38, 39, 3], with specialized tools developed to address the unique challenges posed by degraded, short aDNA fragments [40, 41].

While functional annotation of sequences can be achieved by mapping reads to protein databases, assembly-based methods often provide more accurate and comprehensive results, particularly in metagenomic studies with many unknown or distantly related species [42]. *De novo* assembly is commonly used to uncover a wide range of elements, from protein-coding genes to entire genomes. This approach is especially advantageous for ancient samples, where modern references may fail to capture distantly related species. Moreover, short fragment lengths and characteristic aDNA damage patterns further complicate alignment-based methods [43].

Several studies have demonstrated the applicability of *de novo* assembly for searching aDNA samples for antibiotic-resistance genes or unknown metabolites, or for reconstructing whole genomes [44, 17, 45, 46, 47, 48, 49]. For instance, a study by Wan *et al.* [50] recently demonstrated the potential of mining proteomes from extinct species, termed the “extinctome”, to identify novel antimicrobial genes. Considering that approximately 90% of prokaryotic genomes are protein-coding [51, 52, 53], *de novo* assembly of extinct species is the gateway for revealing unknown proteomes. While Klapper *et al.* [17] highlighted the fact that deeply sequenced datasets can be assembled using conventional assembly algorithms, the assembly process becomes more challenging when dealing with samples that exhibit high damage rates and low coverage.

To demonstrate the effects of varying fragment length and damage patterns, we simulated fragments of a simple metagenomic environment with different fragment sizes and deamination rates. As shown in Fig. **1(A)**, fragment length and damage drastically influence the outcome of ancient metagenome assembly. The damage patterns used for the simulations (Fig. **1(B)**) represent two levels of damage intensities as they can be found in studies profiling ancient microorganisms. Even at moderate damage levels with the first position of the 5^′^ end exhibiting a damage rate of ∼35%, we observed a significant drop in assembly performance. Several studies have documented rates of nucleotide misincorporation at the 5^′^ end, ranging from 15% to 60% [17, 36, 54, 55, 56, 3, 57]. Our findings highlight the need for optimized assembly algorithms to address the unique characteristics of ancient metagenomic datasets.

**Figure 1:**
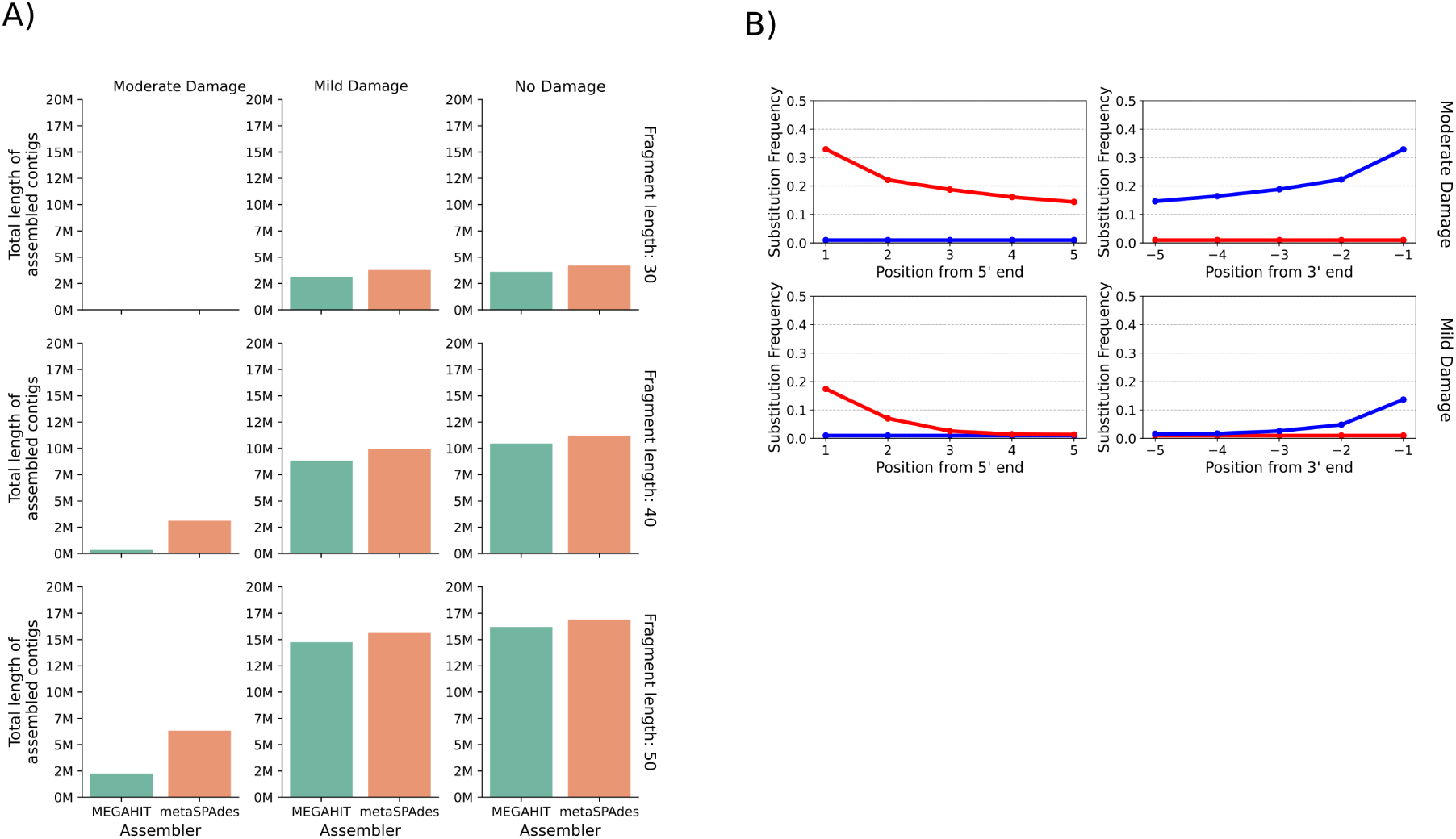
**A)** Impact of aDNA damage and fragment length on metagenomic assembly. The plots show the sum of all contigs larger than 500bp after assembling a simulated toy dataset with either MEGAHIT [58] or metaSPAdes [59]. Each bar plot refers to a different combination of fragment length and damage rate of the simulated data. While empirical data is inherently more complex in terms of variation in fragment lengths and damage patterns, our simplified dataset demonstrates how common assemblers are limited by aDNA damage. **B)** Damage patterns used for the simulation of aDNA fragments. We used two levels of deamination rates for the simulations: moderate damage and mild damage compared to the rates found in ancient microbial studies [17, 36, 54, 55, 56, 3, 57]. The blue traces represent C-to-T substitution rates, while the red traces indicate G-to-A substitution rates.

There are two major classes of *de novo* assembly algorithms: Overlap Layout Consensus (OLC) and de Bruijn graphs. OLC algorithms rely on the computation of full read overlaps, while de Bruijn graph algorithms construct contigs using sub-sequences of length *k* (*k*-mers) [60]. Although OLC algorithms are precise, they are computationally impractical for assembling large datasets produced by high-throughput sequencing technologies [61, 62]. Consequently, de Bruijn graph algorithms have become the prevalent algorithmic basis for most assemblers developed in the past decade. De Bruijn graph assemblers connect *k*-mers that overlap by at least *k*-1 nucleotides in a graph structure, thereby efficiently assembling short reads into contigs [63, 64, 58]. However, de Bruijn graph assemblers face a precision-sensitivity trade-off when assembling complex metagenomic data. Longer *k*-mers provide higher specificity for individual species but suffer from reduced sensitivity in populations with high intra-species diversity. Conversely, shorter *k*-mers offer increased sensitivity but lack the specificity required to effectively distinguish between closely related species [65, 62]. Moreover, as *k*-mer length increases, sequencing errors are more likely to interfere with matching *k*-mers correctly, making read correction methods necessary to address this issue. Ancient metagenomic datasets, characterized by damaged and short fragments, amplify this issue by naturally increasing diversity and complexity and thereby stretching the limits of de Bruijn graph assemblers [6]. New assembly strategies are needed to overcome these issues.

The *de novo* metagenomic assembler PenguiN [62] uses a greedy-iterative overlap-based assembly method. Inspired by the protein level assembler PLASS [65], it clusters reads based on shared *k*-mers. Extension candidates are then selected from these clusters using a Bayesian model that leverages alignment length and sequence identity between the query and extension candidate in both protein and nucleotide space. The extension process continues iteratively, first clustering, then extending. Clustering reduces the computational complexity of finding overlaps from quadratic in the number of reads to quasi-linear. This overlap-centric approach avoids the precision-sensitivity trade-off seen for de Bruijn graph assemblers, resulting in improved recovery from highly diverse metagenomic datasets [65, 62].

Here we introduce CarpeDeam, a *de novo* assembler specifically designed for ancient metagenomic data, based upon the greedy-iterative workflow of PenguiN [62]. It addresses the aforementioned challenge of high damage levels. Each iteration comprises three phases: First, sequences are clustered based on shared *k*-mers analogues to PenguiN. CarpeDeam additionally refines clusters by using a filter that employs a purine-pyrimidine encoding [66]. This encoding, which we refer to as RYmer space, ensures that cluster assignments of sequences are robust to aDNA damage events. In the second phase, sequences with base substitutions that are likely to be due to damage in the aDNA fragments are corrected. The final phase involves the elongation of contigs through an extension rule that takes into account aDNA-specific substitution patterns.

We introduce CarpeDeam, a novel assembler specifically designed for ancient metagenomic datasets. Our analysis highlights the challenges assemblers face when dealing with aDNA datasets, as performance varies greatly depending on fragment length distributions and damage patterns. CarpeDeam demonstrates superior performance in simulated datasets and in many empirical datasets, recovering unique genomic segments that are missed by other assemblers. CarpeDeam offers two modes: a default “safe” mode for reducing errors and an “unsafe” mode for increased sensitivity. These promising results mark an important step forward in damage-specific assembly; however, further advancements will be needed to fully address the complexities of ancient metagenomic datasets.

## 3 Results

### Workflow of CarpeDeam

CarpeDeam is an assembler based on the metagenome assembler PenguiN. It employs a similar greedy, iterative overlap strategy, yet it incorporates several critical adjustments tailored for assembling ancient metagenomic data. Whereas PenguiN utilizes six-frame translated reads to find overlaps in amino acid space, we omit this step as ancient DNA fragments are ultrashort, resulting in even shorter amino acid sequences. In the text, we use the term aDNA fragments to describe the trimmed and merged sequencing reads (see discussion in Lien *et al.* [67]).

Overall CarpeDeam relies on three iteratively repeated steps: During **PHASE 1** (see Fig. **2**), sequences are clustered based on shared *k*-mers via the MMseqs2 linclust algorithm [68]. In this process, each sequence within a cluster is required to align to the center sequence with a minimum sequence identity. The center sequence is defined as the longest sequence within the cluster and becomes the focus for subsequent correction or extension processes, depending on the stage. The remaining sequences in the cluster, termed member sequences, overlap with the center sequence and contribute to its damage correction. In the extension phases, they serve as extension candidates. While PenguiN applies a relatively high default sequence identity threshold (99%), the presence of deaminations in aDNA requires a reduction in this threshold as the sequence identity is naturally lower even for sequences of the same provenance. CarpeDeam filters clusters using a reduced sequence identity threshold of 90% while introducing the concept of RYmer sequence identity, which converts sequences to a reduced nucleotide alphabet of purines (adenine and guanine) and pyrimidines (cytosine and thymine) to account for deaminated bases. First, the clusters are generated with all member sequences sharing at least one *k*-mer with the center sequence and the sequence similarity of the full overlap is computed following PenguiN’s workflow. Additionally, we compute the sequence similarity in RYmer space, where an A or a G is encoded as R (one letter encoding for purine) and a C or T as Y (one letter encoding for pyrimidine), of the full overlap. For a detailed explanation of this concept, please refer to Section S7 of the Supplementary Material. Overall, the RYmer space sequence identity allows for mismatches due to deamination events.

**Figure 2:**
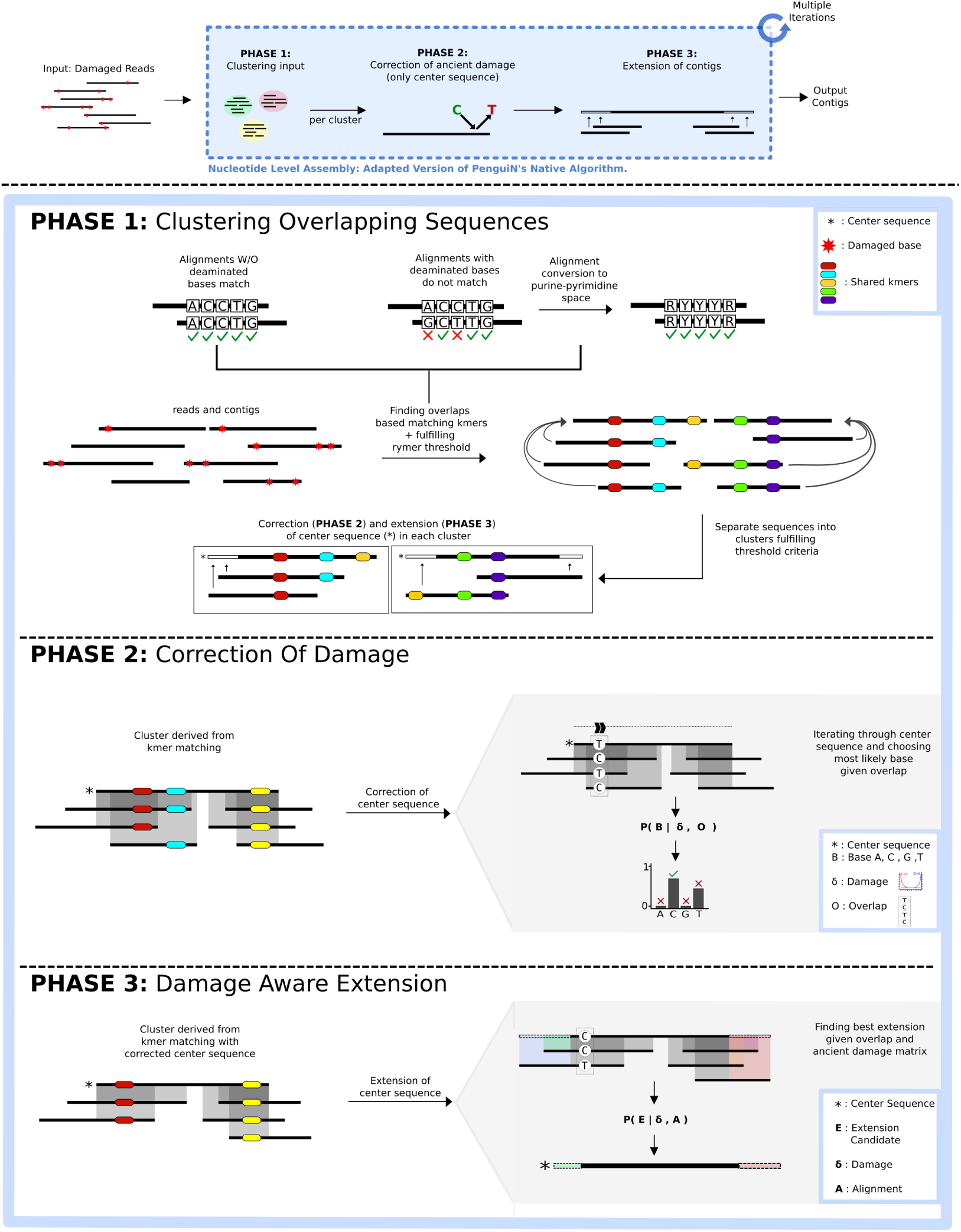
CarpeDeam’s main workflow: The input are aDNA sequences (FASTQ format) which have been trimmed and, for paired-end data, merged. During an iterative process, the fragments are corrected and extended to long contigs. In **PHASE 1** the fragments are grouped into clusters sharing at least one *k*-mer as well as an overlap sequence identity of 99% in RYmer space. **PHASE 2** corrects deaminated bases. In particular, the center sequence of each cluster (which is always the longest sequence) is assigned the most likely base per position given the evidence of overlapping sequences in the cluster and the user-provided damage patterns. In **PHASE 3**, the center sequence of each cluster is extended by the candidate sequence from the cluster that is most likely to be the correct extension. In fact, **PHASE 3** is divided into two steps. First, only aDNA fragments (non-extended sequences) are taken into account for extension, as the provided damage patterns are only valid for non-extended sequences. In the second step of **PHASE 3** exclusively contigs (sequences that already have been extended at least once) are used for the extension, applying a modified Bayesian extension model from the native PenguiN assembler.

In **PHASE 2** (see Fig. **2**), CarpeDeam corrects deaminated bases. Any base in the center sequence of a cluster that is covered by at least one other sequence from the cluster can be corrected. Employing the user-provided damage patterns, the method utilizes a maximum-likelihood estimation to infer the most probable base for each position. The likelihood model is explained in more detail in Methods Section 6.

During **PHASE 3** (refer to Fig. **2**), sequences are extended. PenguiN employs an extension rule based on a Bayesian model, which selects the most suitable extension for each cluster. In contrast, we split the extension process into two distinct phases. The initial phase exclusively targets non-extended sequences (i.e. aDNA fragments). The key aspect is that the damage pattern provided by the user is only valid for these sequences, enabling the application of our likelihood model. Consequently, the initial phase focuses on extending the center sequence using yet non-extended sequences. Within each iteration, the extension continues as long as our likelihood model supports a strong belief in the accuracy of the extension (see Methods Section 6). The subsequent phase involves merging contigs: sequences that have already been extended and corrected. For this, CarpeDeam applies PenguiN’s Bayesian model with adjustments to improve its applicability to ancient datasets (see Methods Section 6).

We assessed the performance of CarpeDeam by generating 18 simulated datasets derived from three distinct ancient environments: gut microbiome [44], dental calculus [35], and bone material [69]. For each environment, six datasets were created with varying levels of coverage and damage. Furthermore, we assembled and evaluated two empirical datasets, performing taxonomic classification and protein similarity search on the contigs as evaluation criteria (see Methods Section 6 for further details).

According to our findings, MEGAHIT [58] and metaSPAdes [59] are the metagenomic assemblers of choice and have been applied in published studies for *de novo* assembly of ancient metagenomic datasets [44, 17, 45, 46, 47, 48, 49]. To evaluate CarpeDeam’s efficacy, we conducted an extensive benchmark comparison against the state-of-the-art assemblers MEGAHIT and metaSPAdes as well as the novel assembler PenguiN [62].

Overall, during our analysis we found that CarpeDeam can produce an elevated number of misassemblies. To offer users more flexibility in managing this trade-off, we introduced two operational modes for CarpeDeam: safe and unsafe. The safe mode (set as default) additionally implements a consensus-calling approach during the extension phase to reduce misassemblies. Users can opt for the unsafe mode, which disables only the consensus calling mechanism, potentially increasing sensitivity at the cost of a higher rate of chimeric contigs. This dual-mode approach allows users to balance between assembly sensitivity and accuracy based on their specific research needs and dataset characteristics. A detailed description of the consensus approach can be found in Section 6.

### Simulated Data

#### Evaluation Strategy of Assembly Quality with metaQUAST and Prokka

We simulated 81 datasets of metagenomic paired-end Illumina short reads of three different environments by varying three parameters: average coverage depth (3X, 5X, and 10X), fragment length distributions (medium, short, and ultra-short), and DNA damage patterns (moderate, high, and ultra-high). The simulation procedure is described in Section 6. The taxonomic profiles of our simulated datasets were derived from three publications [69, 35, 44], representing varying levels of complexity in species diversity. For instance, the most complex dataset mirrors a gut microbiome consisting of 116 identified microbial species derived from Wibowo *et al.* [44], with individual species abundances ranging from a maximum of 10.77% to a minimum of 0.011%. At 10X average coverage, this results in the least abundant species having a coverage of only 0.15X. While all species had the same simulated rate of damage and fragmentation patterns, we also created non-uniform simulations where different species have different rates of damage and fragmentation patterns (see Supplementary Material, Section S4).

We assembled the simulated datasets with CarpeDeam, PenguiN [62], MEGAHIT [58] and metaSPAdes [59].

All assemblers were run with their default parameters, except for the minimum contig length, which was set to a minimum of 500bp. For CarpeDeam we ran both the *safe* and the *unsafe* mode. We specified the flag--only-assembler for metaSPAdes because we observed the program getting stuck in the step that aims to correct sequencing errors. Skipping this step was advised in the issues section of the metaSPAdes GitHub repository (issue 306).

The reference genomes underlying our simulations were used to evaluate the assemblies of all four assemblers. We used the original references to compute the following alignment-based metrics: NA50, LA50, largest alignment, duplication ratio, misassemblies and mismatches per 100kb as generated by metaQUAST [70]. We set the sequence identity threshold at 90% for the metaQUAST [70] alignments as suggested in the metaQUAST documentation. Additionally, we mapped the aDNA fragments back to the contigs using Bowtie2 [71] with--very-sensitive-local and reported the fraction of mapped fragments against the contigs via SAMtools [72].

### Comparison of Assemblers Based on Standard Metrics Reported by metaQUAST

For our benchmark, we set the minimum contig length to 500 bp. Figure **3** presents four key metrics—largest alignment, number of misassemblies per contig, covered genome fraction, and NA50—for nine selected datasets: moderate damage and short fragment length at coverage depths of 3X, 5X, and 10X. Results for all 81 datasets are provided in Section S2 of the Supplementary Material.

**Figure 3:**
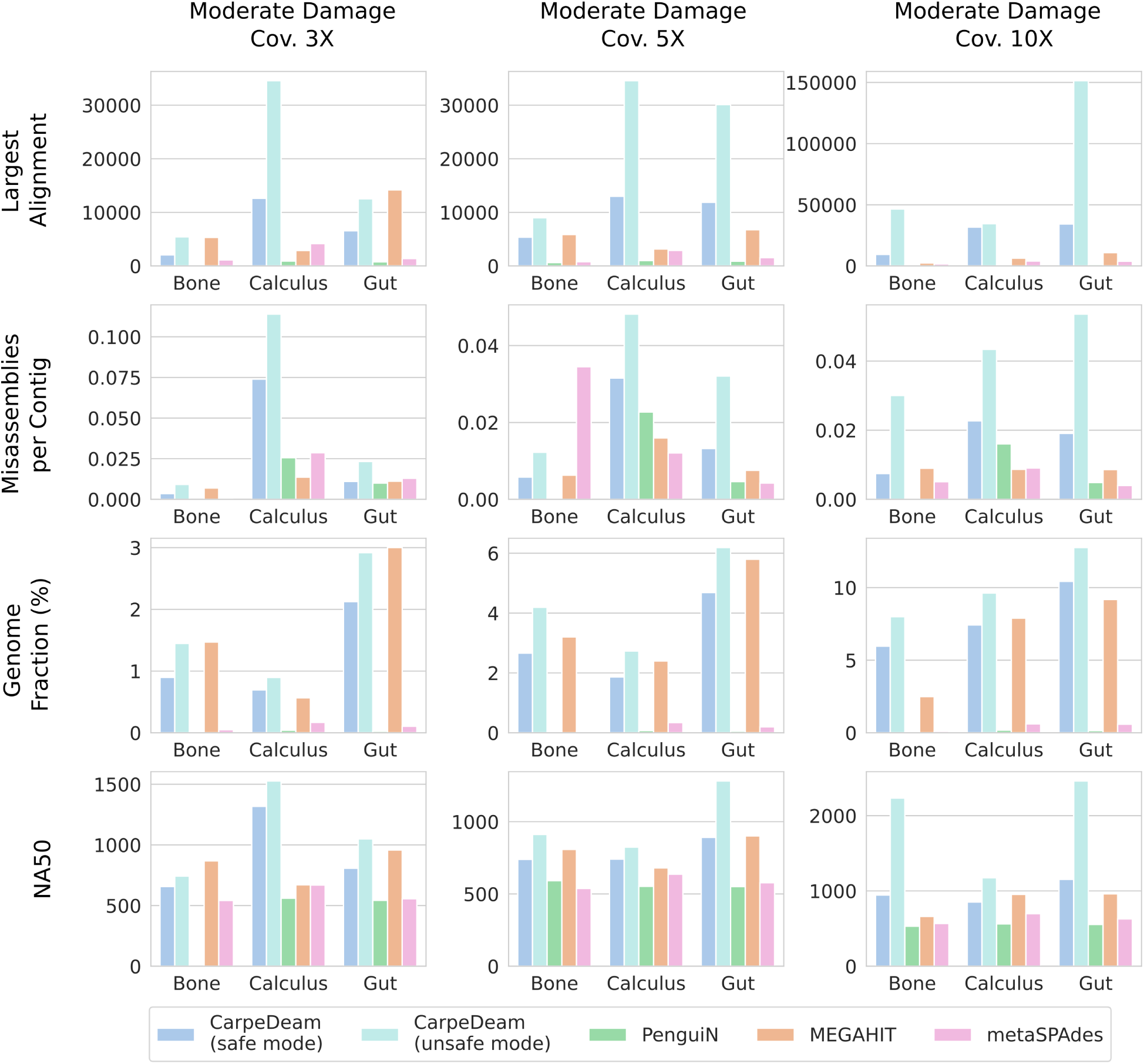
Performance evaluation of assemblers CarpeDeam (safe and unsafe modes), MEGAHIT, metaSPAdes, and PenguiN across nine simulated datasets. Results are presented for datasets with category moderate damage and short fragment length distribution, simulated for three environments (gut, dental calculus, and bone) and three coverage levels (3X, 5X, and 10X). The metrics shown are largest alignment (row 1), misassemblies per contig (row 2), genome fraction (row 3), and NA50 (row 4). Each bar represents the performance of an assembler for a specific metric, coverage, and environment.

CarpeDeam, in both its safe and unsafe modes, and MEGAHIT demonstrated strong performance across the presented assembly metrics. In contrast, PenguiN and metaSPAdes were less effective at assembling comparable fractions of the metagenomes.

A notable difference was observed in the “largest alignment” metric (Fig. **3**, row 1), where CarpeDeam’s unsafe mode consistently produced substantially longer contigs compared to other assemblers. This advantage was particularly evident in the calculus and gut datasets. For most datasets even CarpeDeam’s safe mode yielded larger maximum alignments than MEGAHIT, while not achieving ultra-long alignments as the unsafe mode did. Only for the 3X gut dataset, MEGAHIT had a larger alignment than both CarpeDeam modes. PenguiN and metaSPAdes both had significantly shorter maximum alignments.

As expected, the increased sensitivity of CarpeDeam’s unsafe mode came at the cost of a higher misassembly rate, as shown by the “misassemblies per contig” metric (Fig. **3**, row 2). For the 3X gut dataset CarpeDeam’s unsafe mode generated more than twice as many misassemblies per contig than MEGAHIT, for the 5X gut dataset four times as many per contig than MEGAHIT and for the 10X gut datasets six times as many per contig than MEGAHIT.

Although CarpeDeam’s safe mode exhibited an elevated misassembly rate compared to MEGAHIT, PenguiN, and metaSPAdes, the difference was less pronounced. The most notable difference was observed in the 10X gut datasets, where the safe mode created about 2.5 times as many misassemblies per contig as MEGAHIT.

The fraction of recovered genomic content is shown in row 3 of Fig. **3** and CarpeDeam (both modes) score the highest in this category. Although the genome fraction recovered by MEGAHIT varied across datasets, it generally fell between the values obtained by CarpeDeam’s safe and unsafe modes. For the gut dataset the values of the recovered genomic fractions were for CarpeDeam’s safe mode, CarpeDeam’s unsafe mode and for MEGAHIT (in that order): 2.1%, 2.9% and 3.0% for the 3X dataset, 4.7%, 6.2% and 5.8% for the 5X dataset and 10.4%, 12.8% and 9.2% fot the 10X dataset. Both metaSPAdes and PenguiN recovered a substantially lower genome fraction.

Finally, the NA50 metric is presented in Fig. **3**, row 4. Generally, the N50 metric represents the length of the shortest contig such that half of the total assembled length is contained in contigs of this length or longer. While N50 is a commonly used metric to get a first impression of assembly quality, it does not distinguish between contigs that align with a reference genome and those that do not. The NA50 metric on the other hand considers only those contigs that have aligned to a reference genome. This adjustment helps to mitigate the influence of long chimeric or misassembled contigs that might artificially inflate the N50 value.

The top three performers in terms of NA50 values are CarpeDeam (in both safe and unsafe modes) and MEGAHIT. It is worth noting that CarpeDeam’s unsafe mode sometimes exhibits significantly higher NA50 values for certain datasets.

Table 1 presents additional metrics reported by metaQUAST [70] for evaluating the gut datasets with moderate damage and short fragment length distribution, representing the most complex of the three samples. Additional results for the dental calculus and bone datasets, which are progressively less complex, as well as other parameter combinations of fragment length distributions and damage patterns, are provided in the Supplementary File 1.

**Table 1:**
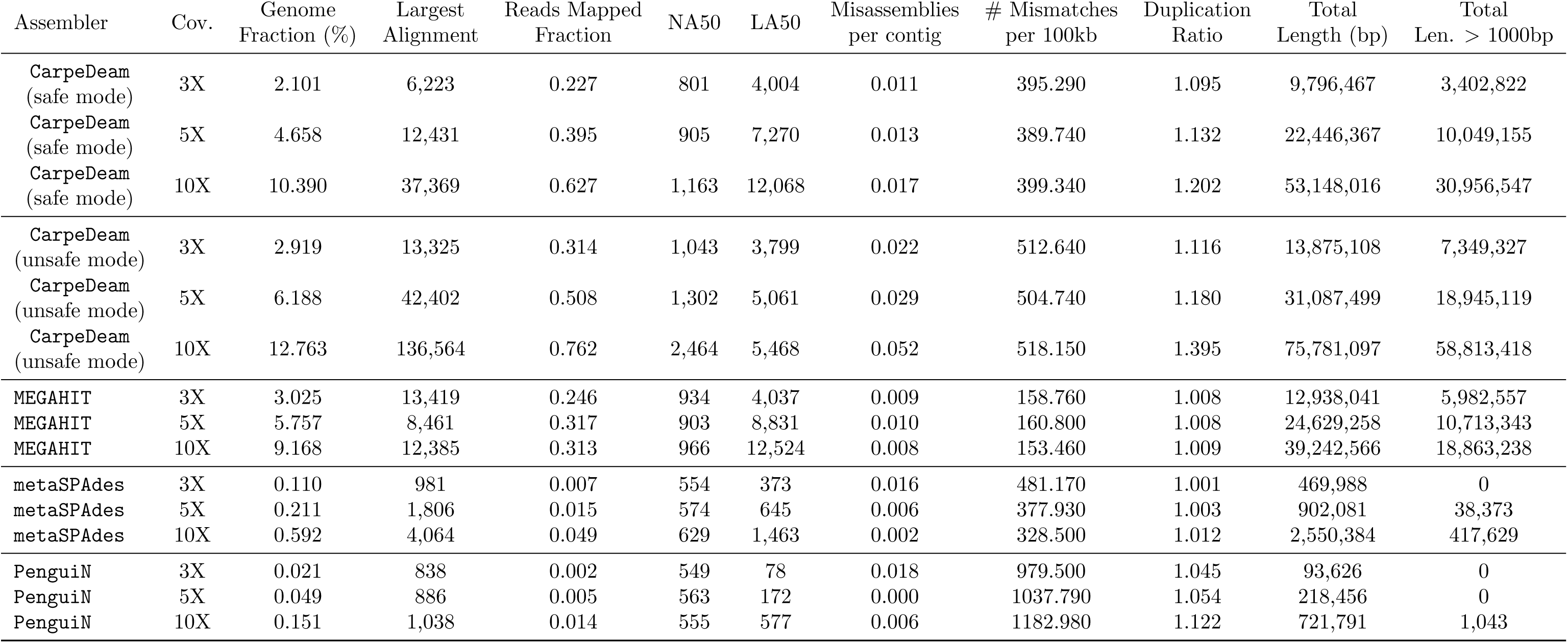
Assembly evaluation metrics for the simulated gut metagenomic environment with moderate damage and short fragment length distribution.

The table shown also includes the fraction of aDNA fragments mapped against the assembly. Across all three datasets, CarpeDeam’s unsafe mode consistently demonstrated a higher number of aDNA fragments mapping to its contigs than other assemblers. Notably, MEGAHIT performed very well for the 3X coverage gut dataset, with the highest values for both the “Genome fraction” and “Largest Alignment” metrics.

The metric “mismatches per 100kb” is also presented in Table 1. MEGAHIT consistently exhibited the lowest number of mismatches per 100kb across all coverage levels. Interestingly, PenguiN displayed the highest mismatch values, exceeding 1000 mismatches per 100kb for all three coverages (3X, 5X, and 10X). This observation aligns with the fact that PenguiN does not perform base correction, as it was designed for modern viral metagenomes. In contrast, the de Bruijn graph assemblers MEGAHIT and metaSPAdes employ bubble merging algorithms to resolve bases in cases where multiple possibilities exist.

Despite CarpeDeam recovering significantly more than PenguiN, the higher mismatch rate per 100kb in CarpeDeam is indicative of its damage correction process, which is based on PenguiN. CarpeDeam’s safe mode shows a lower mismatch rate per 100kb than its unsafe mode, yet is higher than MEGAHIT. For instance, for the 5x gut dataset in Table 1, the mismatch rates are 449 mm.p 100kb (CarpeDeam safe), 613 mm.p 100kb (CarpeDeam unsafe), 159 mm.p 100kb (MEGAHIT), 1147 mm.p 100kb (PenguiN), 379 mm.p.100kb (metaSPAdes). As indicated in Table 1, CarpeDeam exhibits a higher duplication rate. This impacts the mismatch rate per 100 kbp because the number of mismatches computed by metaQUAST is proportional to the duplication ratio. Consequently, mismatch rates can be elevated when there are numerous overlapping alignments, as each assembly alignment is analysed independently. Additionally, the higher misassembly rate contributes to the mismatch rate. For instance, a misassembled contig aligning to the reference can accumulate mismatches at the ends of the aligned region due to chimeric nature of the contig. Since metaQUAST tolerates a certain degree of sequence divergence, these mismatches are included in the calculation, further elevating the mismatch rate.

The tables reveal a general trend where CarpeDeam demonstrates higher sensitivity, producing a larger number of contigs. This is indicated by its elevated duplication ratio and higher mapped fraction, while achieving a genome fraction similar to MEGAHIT. Our observations align with the findings presented in the PenguiN [62] manuscript: PenguiN’s approach recovers more strain-resolved viral genomes, while generally exhibiting a higher duplication rate. In contrast, MEGAHIT appears to prioritize precision, generating fewer contigs with a lower rate of misassemblies, resulting in capturing less of the genomic diversity present in the sample.

We show NA50 and LA50 in Table 1 as they can be viewed as more informative as their “non-aligned” counterparts N50 and L50. A higher NA50 is considered better as it describes the minimum alignment length to be considered, covering half of the alignment length of all contigs. For LA50, however, a lower value is considered better, as it describes the number of aligned blocks that need to be considered to cover half of the alignment length of all contigs. It must be noted that these metrics need to be evaluated in the context of the covered genome fraction. For instance, while PenguiN and metaSPAdes both have small LA50 values, the tools recovered a much smaller genome fraction than CarpeDeam and MEGAHIT.

### Impact of Fragment Length Distributions and Damage Levels on Assembly Performance

Our simulation of three different samples, with varying coverages, fragment lengths, and damage profiles, resulted in a total of 81 datasets. The results of these assemblies are presented in the Supplementary Material, Section S2, as extended heatmaps for various metrics reported by metaQUAST [70].

There were several key observations. Fragment length distributions had a notable impact on the assembly performance. The distributions are shown in the Supplementary Material, Fig. S1 (A). Assemblies derived from the medium fragment length distribution (median 58 bp) generally outperformed those from the short (median 47 bp) and ultra-short (median 42 bp) distribution (Supplementary Material, Fig. S2 - S13). Interestingly, datasets with ultra-short fragments often recovered a slightly higher genome fraction than those with short fragments, despite the latter distribution having a longer median fragment length. This trend is particularly evident for assemblers such as CarpeDeam, PenguiN, and metaSPAdes, while it is less pronounced for MEGAHIT. The distributions differ in their maximum fragment lengths, with the ultra-short distribution reaching up to 140 bp and the short distribution reaching only 120 bp. These rare longer fragments likely have a disproportionate influence on assembly performance.

Damage profiles also played a critical role in assembly outcomes. Counterintuitively, the parameter combination of ultra-high damage and medium fragment lengths yielded the highest genome fractions in many cases. This contrasts with the moderate damage datasets, which consistently performed worse. A closer examination of the damage profiles revealed that the slope of the substitution rates across positions is a significant factor. While moderate damage had the lowest substitution rate at position 1 of the simulated fragments, it maintained relatively high rates at position 5, suggesting that a steeper decline in substitution rates positively influences assembly performance.

### Evaluation of Non-Misassembled Contigs Across Assemblers

To further evaluate the quality of assemblies beyond the genome fraction metric, we assessed the ability of each assembler to reconstruct long, non-misassembled contigs as classified by metaQUAST. While genome fraction provides valuable information, it does not account for the length of individual contigs mapping back to the reference genome (by default metaQUAST considers all alignments over 65bp). Consequently, a high number of short contigs may inflate the genome fraction with-out necessarily representing superior assembly quality. To address this limitation, we analysed the number of non-misassembled contigs exceeding 2000 bp for each assembler across the datasets with short fragment length and moderate damage.

We demonstrate CarpeDeam’s capacity to generate longer accurate contigs across datasets, shown in Supplementary Material, Fig. S18. CarpeDeam (unsafe mode) consistently produced more long contigs for all datasets except one (3X coverage bone dataset, where MEGAHIT outperformed both CarpeDeam modes). CarpeDeam’s safe mode and MEGAHIT exhibited alternating performance, with CarpeDeam producing more longer non-misassembled contigs in four cases and MEGAHIT in four cases. Notably, PenguiN and metaSPAdes generated significantly fewer long contigs compared to other assemblers. Although CarpeDeam produced a higher number of misassemblies, which would require careful filtering in downstream analyses, the increased quantity of non-misassembled contigs over 2000 bp suggests a notable improvement in assembly contiguity.

### Comparison of Non-Coding Genomic Content Across Assemblers

To complement our metaQUAST alignment analysis, we performed an additional examination of the assembly fractions (Fig. **4**). We used skani [73], a tool for calculating average nucleotide identity (ANI) and aligned fraction (AF), employing an approximate mapping method without base-level alignment. Contigs shorter than 1000bp were excluded to focus on long genomic segments.

**Figure 4:**
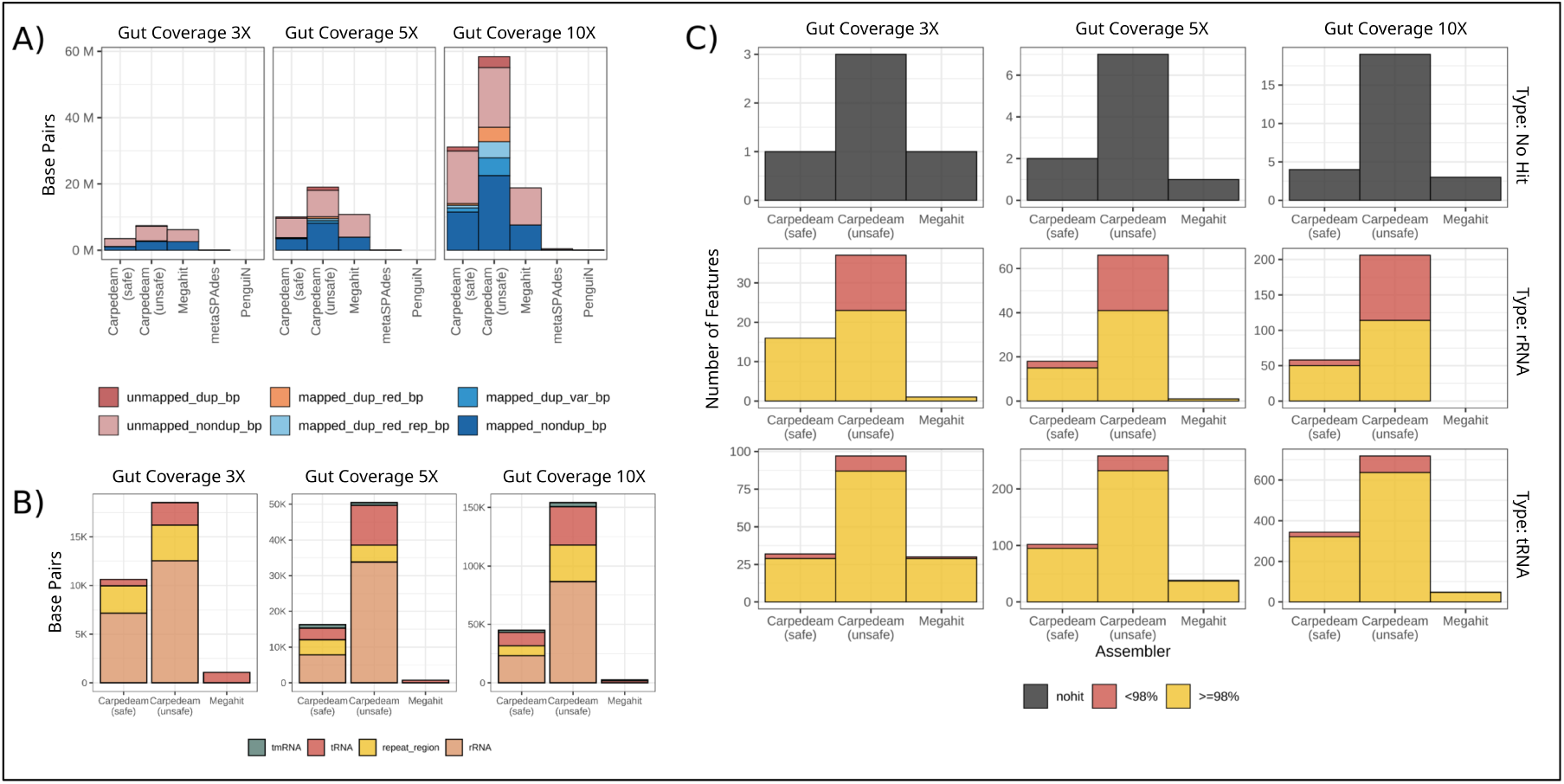
Analysis of mapped fractions of base pairs, genomic features, and RNA recovery across different assemblers and coverage levels for the gut dataset. (A) Distribution of mapping categories (mapped non-duplicated, mapped duplicated variant, mapped duplicated redundant representative, mapped duplicated redundant, unmapped non-duplicated, and unmapped duplicated base pairs) for assemblies of the gut dataset at 10X, 5X, and 3X coverage levels. (B) Types of genomic features recovered in contigs based on Prokka annotations, for different assemblers (gut dataset). (C) Recovery of rRNAs and tRNAs with sequence identities above and below 98% for MEGAHIT and CarpeDeam (safe and unsafe modes) across different coverage levels (gut dataset; moderate damage, short fragment length distribution).

We categorized the mapping results into groups to provide a detailed evaluation of the assembly quality and its ability to handle redundancy and variation. The category *unmapped nondup bp* includes non-redundant nucleotides in the assemblies that do not map to the reference with an average nucleotide identity (ANI) of over 99%. These sequences represent unique content that was not successfully aligned to the reference. In contrast, *unmapped dup bp* comprises nucleotides in the assemblies that also do not map to the reference but include redundant sequences, reflecting over-represented regions introduced by the assembler.

For mapped base pairs, *mapped dup var bp* consists of duplicates that are highly similar to reference sequences (ANI *>* 99%) but show minor variations, such as mutations present in some species but absent in others. This category highlights the assembly’s ability to capture genetic diversity. Meanwhile, *mapped dup red bp* represents redundant mapped base pairs that are likely a result of assembling more than one contig for the same genomic region, indicating redundancy in the assembly process.

The category *mapped dup red rep bp* quantifies the subset of redundant mapped base pairs required to create a representative set. By comparing this to *mapped dup red bp*, one can infer metrics similar to the “duplication ratio” reported by metaQUAST, highlighting how redundant base pairs captured by *mapped dup red bp* are distributed. This subset, defined by *mapped dup red rep bp*, represents the unique base pairs within the redundant regions.

Finally, *mapped nondup bp* captures the non-redundant mapped base pairs, excluding the duplicates. This category indicates how much of the reference is covered by base pairs that map with high ANI and are unique, making them especially valuable for downstream analyses.

Overall, we conducted this mapping strategy to provide a more detailed evaluation of the nucleotide fractions in the assembled contigs that can be mapped to the reference. Unlike metaQUAST’s Genome Fraction metric, which applies a minimum alignment length and a fixed sequence similarity threshold, skani offers a more flexible approach to assessing mappings.

Our six categories capture detailed differences in the assemblies, including their ability to manage redundancy, capture genetic variation (e.g., within highly similar genetic regions), and evaluate completeness in terms of reference coverage. These results help users evaluate CarpeDeams’ suitability for their specific research needs. For instance, while higher redundancy may complicate downstream analyses like binning, it poses less of a concern when focusing on specific gene regions, where sensitivity is of greater interest.

Fig. **4**, panel A, presents the results for the gut dataset across 10X, 5X, and 3X coverage levels. Our analysis revealed that CarpeDeam (unsafe mode) exhibited the highest *mapped non dup bp* fraction, particularly in the 10X coverage dataset. This advantage decreased at lower coverages (5X and 3X), with the difference between CarpeDeam and MEGAHIT diminishing. Substantial fractions of *unmapped non dup bp* sequences were observed across assemblies, most notably in CarpeDeam (safe and unsafe) and MEGAHIT. PenguiN and metaSPAdes showed significantly lower fractions across all mapping categories compared to other assemblers.

We further investigated the types of genomic features other than coding regions recovered by different assemblers (Fig. **4**, panel B). For this analysis, we used Prokka’s annotation output. Our results showed that CarpeDeam demonstrated a strong ability to recover rRNA, tRNA and repeat region segments while MEGAHIT assembled significantly less base pairs that could be classified by Prokka. It should be noted that only the results for MEGAHIT and CarpeDeam (safe and unsafe) are presented here, as other assemblers recovered substantially less genomic content overall, making their inclusion in the plot impractical for comparative purposes. The results refer to the gut dataset with moderate damage and the short fragment length distribution.

Fig. **4**, panel C, illustrates the recovery of rRNAs and tRNAs with sequence identities above and below 98%. For tRNAs in the 3X gut dataset, CarpeDeam (safe mode) and MEGAHIT reconstructed comparable numbers of high-identity (*ge*98%) sequences, while CarpeDeam (unsafe) outperformed both, reconstructing more than three times as many. This difference became more pronounced in the 5X and 10X datasets, where both CarpeDeam modes assembled significantly more high-identity tRNAs than MEGAHIT. For rRNA reconstruction we could observe a distinct pattern: MEGAHIT recovered very few, whereas both CarpeDeam modes recovered several rRNAs. For instance, for the 10X gut dataset, CarpeDeam’s unsafe mode could recover more than 200 rRNAs of which more than 100 had a sequence identity of 98%. Notably, CarpeDeam (safe mode) recovered fewer rRNAs overall, but those recovered predominantly exhibited ≥98% sequence identity. Conversely, CarpeDeam (unsafe mode) recovered a larger quantity of rRNA, albeit with a considerable fraction (*<*50%) showing *<*98% sequence identity. Interestingly, some RNA sequences annotated by Prokka in the assemblies failed to match the reference annotation. While CarpeDeam (safe) and MEGAHIT exhibited similarly low rates of these “no hit” instances, CarpeDeam (unsafe) demonstrated the highest occurrence of unmatched annotations.

### Evaluation of Recovered Protein Content in Assembled Contigs

Effectively reconstructing protein-coding sequences is essential for creating detailed microbial gene catalogs. These catalogs organize genes found in microbial communities and serve as references for standardized analysis across different samples and studies, making the accurate reconstruction of protein-coding sequences a crucial aspect of our evaluation [74]. Therefore, we assessed the performance of the assemblers in reconstructing proteins by predicting proteins from the reference genomes of our simulated metagenomes with short fragment length distribution and moderate damage using Prokka [75] and searching for highly similar proteins with MMseqs2 map [76].

Fig. **5** shows the number of predicted open reading frames (ORFs i.e. DNA translated into amino acid space) that exhibit significant similarity to predicted proteins from the reference genomes.

**Figure 5:**
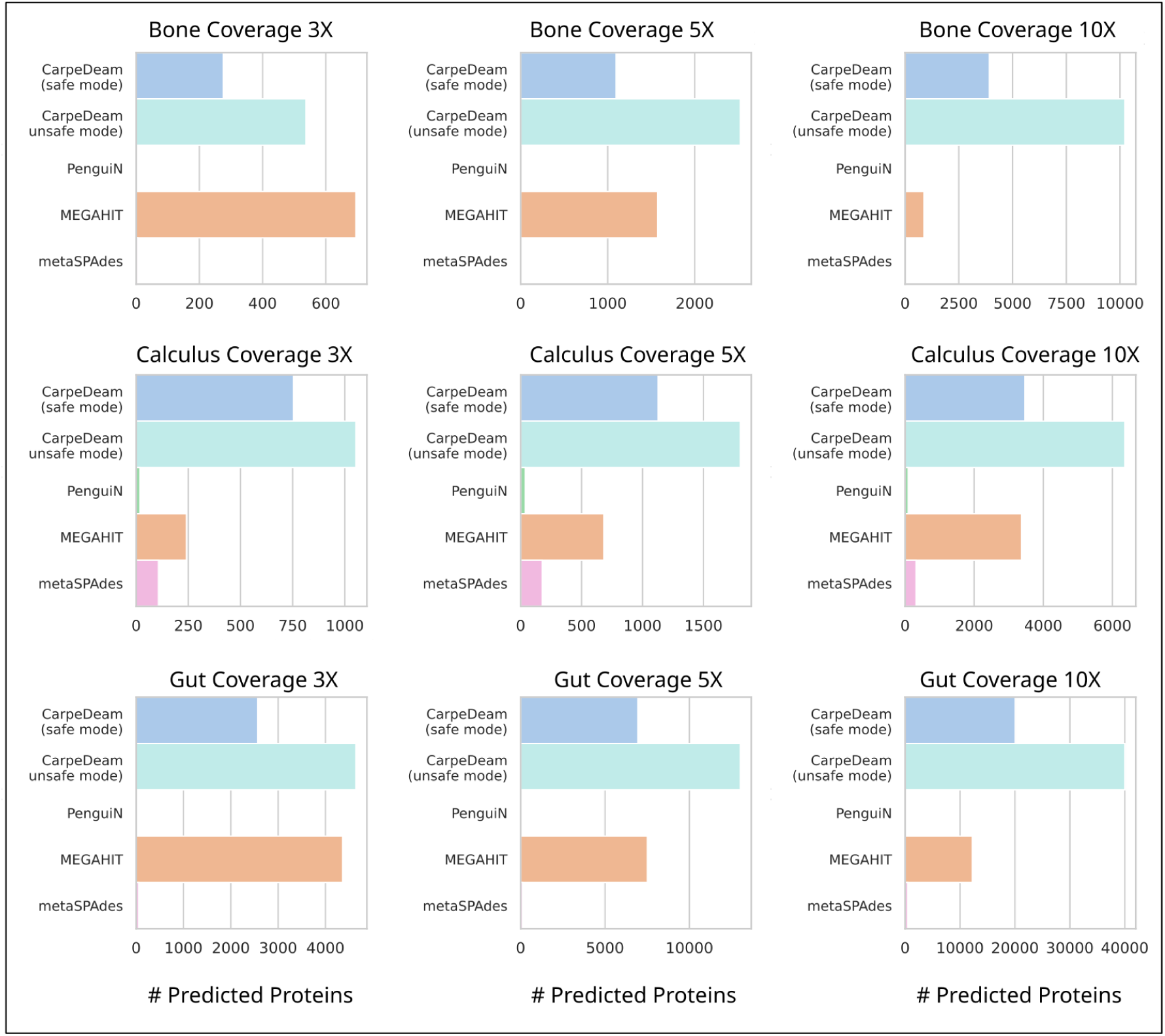
Evaluation of predicted protein sequences. Results are presented for CarpeDeam (safe and unsafe modes), MEGAHIT, metaSPAdes, and PenguiN across datasets with moderate damage and short fragment lengths in three simulated environments (Bone, Dental Calculus, and Gut) with varying coverage levels (3X, 5X, and 10X). The figure shows the number of predicted ORFs with significant similarity to reference proteins, filtered for alignments covering ≥90% of the reference protein and ≥90% sequence similarity.

We reported the number of unique matches in the reference. This approach allowed us to measure the assemblers’ ability to reconstruct biologically meaningful protein-coding sequences.

CarpeDeam’s unsafe mode consistently outperformed other assemblers in assembling more proteins (Fig. **5**, panel **(A)**). However, for the 3X bone dataset, MEGAHIT assembled more proteins than CarpeDeam’s unsafe mode. Apart from the 3X bone dataset, MEGAHIT assembled significantly more proteins than CarpeDeam’s safe mode for the bone 5X and gut 3X datasets. In contrast, CarpeDeam’s safe mode assembled significantly more proteins for all 10X datasets, as well as the 3X and 5X calculus datasets. For the gut 5X dataset, MEGAHIT and CarpeDeam’s safe mode assembled a very similar number of proteins. The other assemblers assembled significantly fewer proteins across all datasets.

As Table 1 of the metaQUAST results indicates that CarpeDeam tends to assemble a higher rate of duplicated contigs compared to other assemblers. Therefore, we conducted additional analysis to assess the impact of duplications on downstream processes. Of particular interest was the potential inflation of unique protein sequence counts in our protein analysis due to duplicated sequences. To address this concern, we performed three additional analyses that account for duplication rates.

First, we clustered predicted proteins with the linclust [68] algorithm at 100% sequence identity and 80% coverage, then searched the cluster representatives against the predicted proteins from the reference using the MMseqs2 map [76] module.

Second, to account for potential duplicates in the reference itself, we clustered the predicted proteins from the reference at 100% sequence identity and 80% coverage. We then searched these clustered reference proteins against the predicted proteins from the contigs using MMseqs2 map.

Finally, we employed miniprot [77] to search the predicted proteins from the reference directly against the DNA contigs. This method bypasses potential undercalling by our protein predictor and focuses on protein-to-DNA alignments directly, providing a complementary perspective to the second analysis. We applied a filter to only obtain hits with at least 95% sequence identity and 95% coverage of the reference proteins. In all cases we reported the number of unique proteins from the reference.

Overall, these additional analyses still favor CarpeDeam, although MEGAHIT recovers similar numbers of proteins especially in the 3X datasets. The detailed results of these analyses are visualized in the Supplementary Material, Section S3.

### Assembly of Empirical Datasets

Next, we extended our analysis to include 20 empirical metagenomic datasets from five distinct sample sites. These datasets were obtained from a study by Fellows Yates *et al.* [36] and represent ancient metagenomic samples from oral microbiomes of Neanderthals and Homo sapiens. The datasets exhibit varying levels of damage, which we used as input for the damage pattern parameter of CarpeDeam. Detailed information about these samples is summarized in Table 2. For the damage pattern input parameter of CarpeDeam, we used the average damage rate across all taxonomies reported.

**Table 2:** aDNA fragments statistics and sample metadata for the empirical ancient oral microbiome datasets from Fellows Yates *et al.* [36]. LSA = Later Stone Age, UP = Upper aleolithic, Meso = Mesolithic, MP = Middle Paleolithic. Library Prep indicates whether the sequencing data was paired-end. The table includes the total number of merged reads, minimum and maximum aDNA read lengths, N50, and GC content.

| Dataset | Microbiome Origin | Period | # Reads (merged) | Length | N50 | GC(%) |
| --- | --- | --- | --- | --- | --- | --- |
| TAF008.B0101 | Modern Human | LSA | 8,929,849 | 35-141 | 53 | 56.17 |
| TAF016.B0101 | Modern Human | LSA | 1,631,218 | 35-141 | 97 | 63.35 |
| TAF017.A0101 | Modern Human | LSA | 1,879,275 | 35-141 | 82 | 56.46 |
| TAF017.C0101 | Modern Human | LSA | 1,981,178 | 35-141 | 47 | 51.25 |
| TAF017.C0101.171215 | Modern Human | LSA | 1,970,380 | 35-141 | 47 | 51.12 |
| TAF018.A0101 | Modern Human | LSA | 5,361,000 | 35-141 | 84 | 63.17 |
| TAF018.B0101 | Modern Human | LSA | 5,327,186 | 35-141 | 57 | 52.6 |
| EMN001.A0101 | Modern Human | UP | 9,911,123 | 35-141 | 69 | 55.41 |
| ECO002.B0101 | Modern Human | Meso | 9,173,902 | 35-141 | 48 | 49.44 |
| ECO002.C0101 | Modern Human | Meso | 8,841,304 | 35-141 | 48 | 49.71 |
| ECO004.B0101 | Modern Human | Meso | 7,748,024 | 35-141 | 52 | 50.08 |
| ECO004.C0101 | Modern Human | Meso | 6,965,132 | 35-141 | 50 | 52.55 |
| ECO006.B0101 | Modern Human | Meso | 7,533,408 | 35-141 | 64 | 61.37 |
| ECO010.B0101 | Modern Human | Meso | 8,903,259 | 35-141 | 48 | 49.89 |
| OAK001.A0101 | Modern Human | LSA | 6,252,614 | 35-141 | 48 | 57.32 |
| OAK003.A0101 | Modern Human | LSA | 13,951,366 | 35-141 | 61 | 53.48 |
| OAK004.A0101 | Modern Human | LSA | 10,976,392 | 35-141 | 53 | 54.21 |
| OAK005.A0101 | Modern Human | LSA | 13,749,370 | 35-141 | 59 | 56.99 |
| GDN001.A0101 | Neanderthal | MP | 13,440,839 | 35-141 | 65 | 58.58 |
| GDN001.B0101 | Neanderthal | MP | 11,972,554 | 35-141 | 52 | 57.12 |

### Identification of Homologous Proteins in Empirical Datasets

Given the absence of ancient reference genomes for the empirical datasets, our evaluation focused on the proteome space. In the context of ancient prokaryotic samples, the ideal scenario would be to identify ancient proteins that exhibit strong similarity to existing sequences while remaining previously undiscovered. To maximize the detection of similar proteins from our assembled contigs, we employed a sensitive screening approach. We used the--easy-search module of MMseqs2 [76] with--search-type 4, which extracts all possible open reading frames (ORFs) from the contigs and searches the resulting amino acid sequences against a provided database. For this analysis, we queried the UniRef100 database [78], which encompasses over 400 million reference protein sequences.

The search results were subsequently filtered using stringent criteria: sequence identity (≥35%), E-value (≤E-12), and alignment length (≥100 residues). Figure **6**, panel **(A)**, presents the number of unique hits in the UniRef100 database, considering only the best hit for each query based on the lowest E-value. This approach allows to assess the potential unknown protein content in our aDNA assemblies while maintaining only highly significant matches.

**Figure 6:**
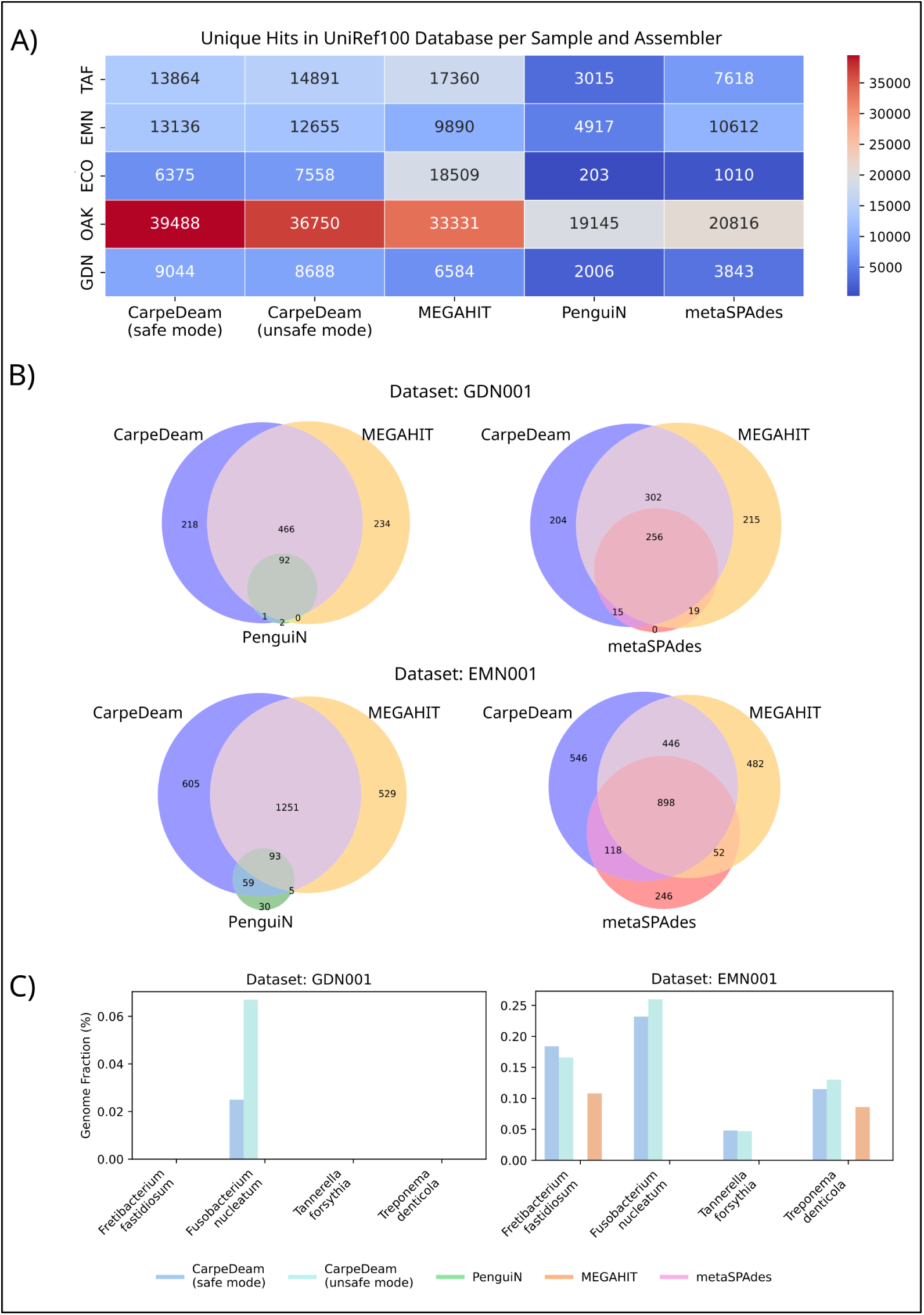
**(A)** Heatmap of unique UniRef100 protein hits for ORFs predicted from contigs assembled by CarpeDeam, MEGAHIT, PenguiN, and metaSPAdes across empirical samples (Grouped by sample site). Hits were were filtered by E-value ≤ *e*^−12^, ≥ 35% identity, alignment length ≥ 100 residues. **(B)** Venn diagrams of species-level taxonomic assignments from translated contigs queried against the Genome Taxonomy Database for the Datasets GDN001.A0101 and EMN001.A0101.**(C)** Recovered genome fraction for highly damaged taxa, as reported by metaQUAST.

Figure 6, panel **(A)**, shows the number of unique protein hits per sample and assembler in a heatmap. The performance of assemblers varies considerably across samples. Overall, MEGAHIT and CarpeDeam generally performed best, with MEGAHIT clearly outperforming all other assemblers in the ECO and TAF samples. In contrast, CarpeDeam achieved the best results in the GDN, EMN and OAK samples.

Interestingly, although metaSPAdes did not perform as well as MEGAHIT in previous analyses, it assembled more unique protein hits in the EMN dataset. When comparing the safe and unsafe modes of CarpeDeam, the safe mode yielded more hits in most datasets, indicating the presence of frameshifts likely caused by misassemblies in the unsafe mode.

### Exploring Taxonomy Assignment through Contig Translation into Protein Space

Another use case for assembled contigs is taxonomic classification. This approach offers several advantages over classifying individual short reads, especially in the context of aDNA analysis. Contig-based classification improves accuracy by leveraging longer sequences, which is particularly beneficial for ultra-short, degraded aDNA fragments. It also helps mitigate misclassifications caused by conserved regions or horizontal gene transfer events [32, 79, 34].

Considering these advantages, we assessed the performance of assemblers in recovering proteins informative for taxonomic assignment using two empirical datasets: EMN001.A0101 and GDN001.A0101. Open reading frames (ORFs) were extracted directly from the assembled contigs using the--easy-taxonomy module of MMseqs2 [34], which translates the contigs in all six reading frames and queries them against the Genome Taxonomy Database (GTDB) [80]. For a detailed description of the workflow, see Section 6.

It is important to note that this approach primarily focuses on detecting bacteria closely related to modern species, specifically those represented in the GTDB, which is derived from RefSeq and GenBank genomes. Consequently, this method may not necessarily identify completely unknown species. The discovery of novel species presents significant challenges, as potential misassemblies would require careful downstream analysis to ensure accurate identification and exclusion of misassembled sequences.

Given these considerations, we adopted a conservative approach with stringent criteria for taxonomic assignment. Matches were filtered to meet the following thresholds: a minimum of 95% sequence identity in amino acid space, at least 70% target protein coverage, and an E-value (≤E-6). This conservative strategy aims to minimize false positives while maintaining high confidence in the reported taxonomic assignments.

We quantified the number of distinct species identified by each assembler based on the proteome alignment (Fig. 6, Panel **(B)**). This metric provides a comparative measure of the assemblers’ performance in recovering taxonomically informative sequences from our empirical aDNA samples.

Figure 6, Panel **(B)**, row 1 presents the taxonomic assignment results for the GDN001 dataset. A substantial number of species were identified by all assemblers, with MEGAHIT and CarpeDeam (unsafe mode) sharing the largest common set. Notably, both MEGAHIT and CarpeDeam identified a considerable number of unique taxa, with MEGAHIT slightly outperforming CarpeDeam (234 vs. 218 unique taxa, respectively; first Venn diagram).

Row 2 of Panel **(A)** displays results for the EMN001.A0101 dataset, revealing a similar pattern to GDN001.A0101. However, consistent with its lower damage rate, the assemblies of the EMN001.A0101 dataset yielded identifications for a larger number of taxa overall. CarpeDeam demonstrated a marginal advantage in unique taxa identification compared to MEGAHIT (605 vs. 529, respectively; third Venn diagram).

A key observation from this analysis is that no single assembler consistently outperforms the others. While the different algorithms naturally share a significant portion of their results, each assembler reveals unique findings.

To further evaluate the assemblers’ performance on highly damaged taxa, we focused on four species reported by Fellows Yates *et al.* [36] to exhibit damage patterns of up to 40% that were identified in both samples: *Fretibacterium fastidiosum*, *Fusobacterium nucleatum*, *Tannerella forsythia*, and *Treponema denticola*. We used the reference genomes of these taxa as input for metaQUAST [70] and reported the recovered genome fraction using a contig length cutoff of 500 bp.

The analysis of the GDN001.A0101 dataset revealed that CarpeDeam was the only assembler capable of recovering any fraction of a reference genome. While no assembler successfully recovered sequences from *F. fastidiosum*, *T. denticola* or *T. forsythia*, CarpeDeam in unsafe mode recovered approximately 0.07% of the *F. nucleatum* genome, whereas safe mode recovered around 0.025%.

In the EMN001.A0101 dataset, most assemblers recovered only small fractions of the reference genomes. Notably, MEGAHIT failed to assemble any contigs longer than 500 bp that mapped to either *F. nucleatum* or *T. forsythia*. Among the assemblers, CarpeDeam performed best, recovering the largest fractions for all four target species. Interestingly, the safe and unsafe modes of CarpeDeam showed similar performance, yielding comparable fractions of the reference genomes.

Table 3 shows the relative abundances of four species presented in Fig. 6, panel **(C)**, derived using MetaPhlan2 [81]. The abundances are generally very low (*<*1%) and differ between the datasets. In GDN001.A0101, *T. forsythia* is absent, while other species have abundances below 0.12%. Notably, only *F. nucleatum* had a small genome fraction assembled by CarpeDeam. In contrast, EMN001.A0101 shows higher abundances for *F. nucleatum* and *F. fastidiosum* (≈0.25%), correlating slightly better with the genome fractions recovered.

**Table 3:**
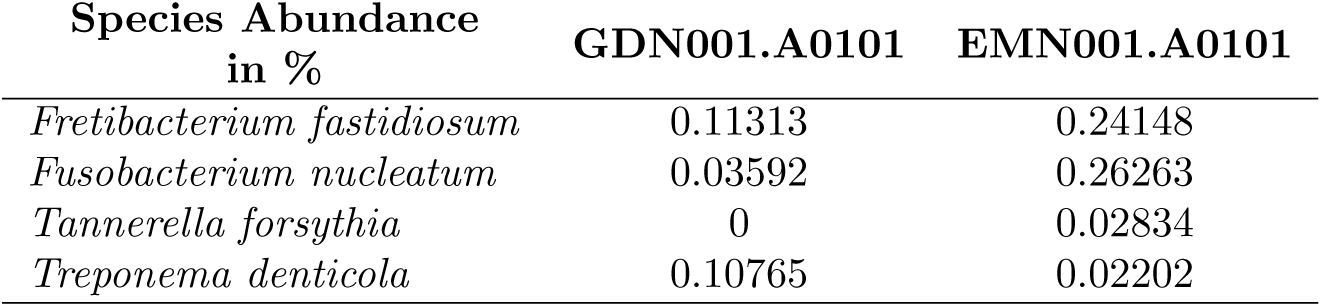
Relative abundances of bacterial species in the two samples as reported by Metaphlan2, found in the data repository of the study by Fellows Yates *et al.* [36].

| Species Abundance<br>in % | GDN001.A0101 | EMN001.A0101 |
| --- | --- | --- |
| *Fretibacterium fastidiosum* | 0.11313 | 0.24148 |
| *Fusobacterium nucleatum* | 0.03592 | 0.26263 |
| *Tannerella forsythia* | 0 | 0.02834 |
| *Treponema denticola* | 0.10765 | 0.02202 |

Overall, no clear proportional relationship is observed between relative abundance and genome recovery, likely due to the low abundances and approximative nature of the tools. Since the contigs were filtered to have a minimum length of 500 bp, this could also contribute to the lack of correlation.

### Detection of rRNA Genes and Potential Biosynthetic Gene Clusters

16S rRNA gene detection is widely used for phylogenetic analysis in metagenomic studies due to the conserved regions of the gene [82]. However, in aDNA studies, detection of the full-length (approximately 1500 bp [82]) 16S rRNA gene is challenging because of the short fragment lengths and characteristic damage patterns inherent to aDNA. Previous studies have targeted several of the nine hypervariable V1–V9 regions [83, 84, 85, 86, 87, 88], which are considerably shorter, thereby requiring the reconstruction of only a few hundred base pairs rather than the full ∼1500 bp gene. Nevertheless, Ziesemer *et al.* [89] demonstrated that targeting specific hypervariable segments in aDNA studies remains challenging when using amplicon sequencing; in such cases, *de novo* assembly can significantly improve the recovery of either full genes or hypervariable regions.

We investigated the detection of 16S rRNA genes by assessing the number of unique hits detected across sequence identity thresholds ranging from 90% to 100%. To ensure robust capture of hypervariable regions, we required that the annotated contigs span at least 80% of a 16S rRNA gene. In Fig. 7, Panel **(A)**, we present the number of unique hits across different sequence identity thresholds rather than relying on a single cutoff.

**Figure 7:**
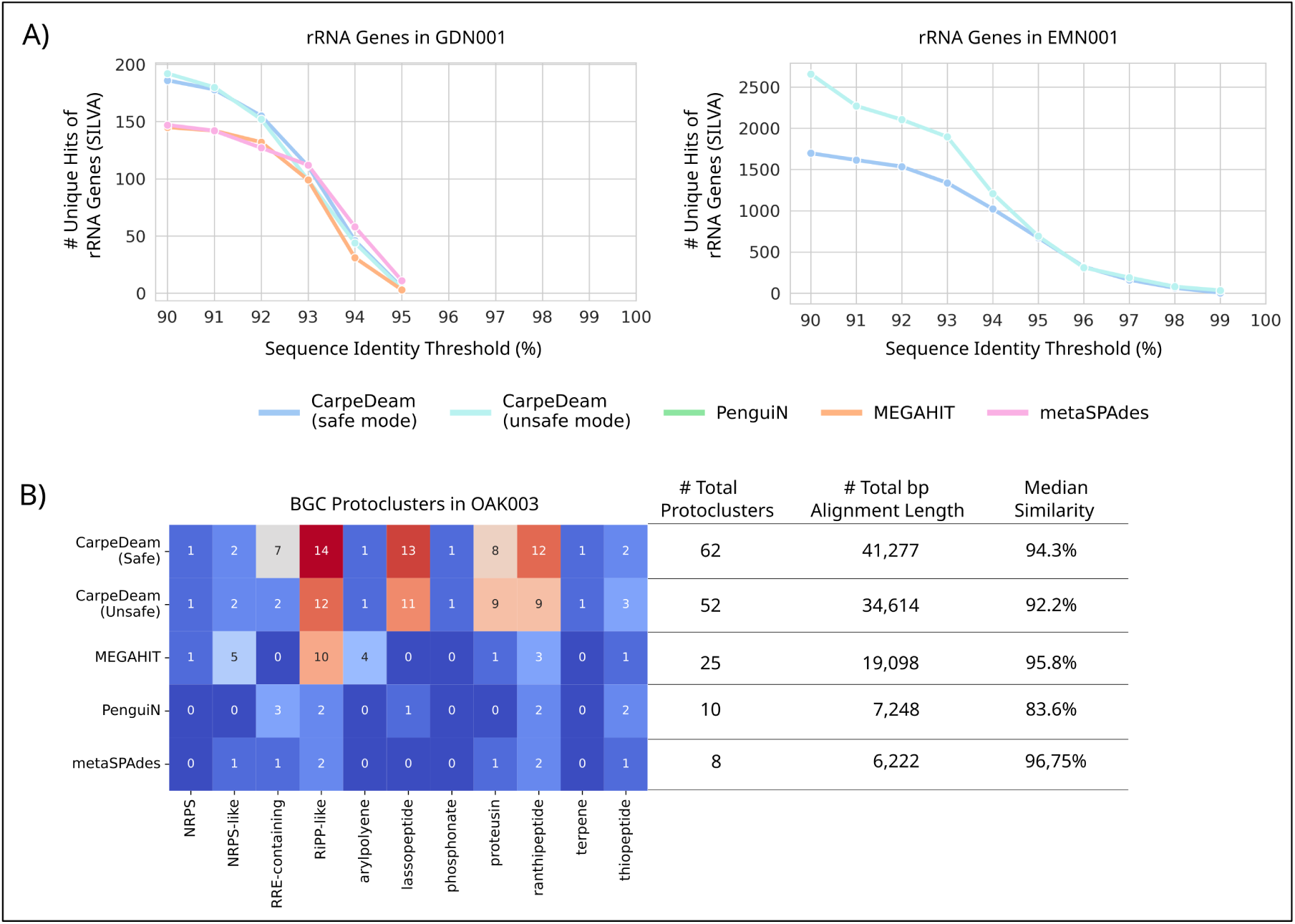
**(A)** Detection of 16s rRNA genes. Shown are the numbers of unique hits in the SILVA database from assembled contigs that were annotated with Prokka and filtered by sequence identity thresholds (90% to 100%) and a minimum coverage of 80%. **(B)** Identification of BGC protoclusters in the OAK003 sample using antiSMASH.

For the GDN001.A0101 dataset, all assemblers performed similarly; however, CarpeDeam in both modes recovered more 16S rRNA hits (*<*200) compared to other assemblers (*<*150) at the 90% similarity threshold. The number of hits converged with increasing sequence identity, and no assembler reconstructed 16S rRNA genes with sequence similarity above 95%. At 95%, metaSPAdes recovered the most hits (11), followed by CarpeDeam and MEGAHIT.

In contrast, the EMN001.A0101 dataset showed a significantly different result. Only CarpeDeam assembled 16S rRNA genes, with the unsafe mode recovering over 2500 hits and the safe mode over 1500 hits at the 90% similarity threshold. At 95% similarity, both modes of CarpeDeam converged, still recovering over 600 unique hits.

For a deeper investigation of the 16S rRNA genes in the EMN001.A0101 dataset, we examined contigs that aligned to the SILVA database covering at least 99% of a full-length 16S rRNA gene with a minimum sequence identity of 95% (a threshold commonly used for genus-level identification [90, 91, 92]). CarpeDeam was the only assembler that produced contigs meeting these criteria. We then assigned taxonomic labels to the recovered 16S rRNA genes and compared these with the genera reported by Fellows Yates *et al.*, where they were using a read alignment approach. To rule out contamination, we also compared the detected genera against a catalogue of potential contaminants provided by Fellows Yates *et al.*. Additionally, we mapped the reads back to the genes to validate their ancient authenticity (PyDamage[93] results in the external data repository). Notably, CarpeDeam recovered three genera on species level (*Actinobacteria*, *Brachymonas*, and *Johnsonella*, shown in bold in Table 4) that were absent in the Fellows Yates *et al.* study and not listed as contaminants. In total, we identified five taxa on species level (sequence similarity ≥99% [94]) and five taxa on genus level (sequence similarity ≥95% [90, 91, 92]). Overall, this analysis underscores the enhanced sensitivity of CarpeDeam and demonstrates its suitability as a complementary method for taxonomic identification alongside read mapping approaches. The result is also remarkable given the low sequencing depth, as the EMN001.A0101 dataset comprised fewer than 10 million merged reads.

**Table 4:**
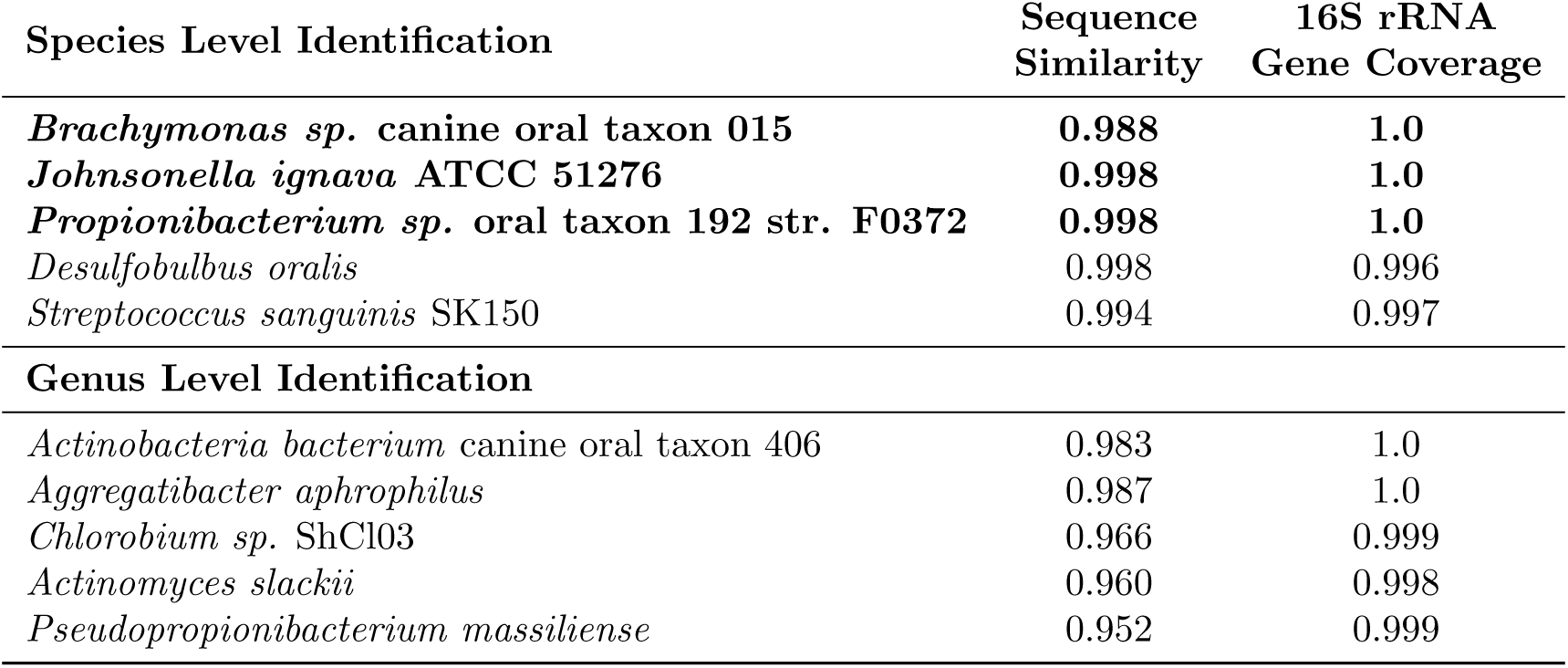
Best hit alignments of assembled contigs to the SILVA 16S rRNA gene database for the EMN001.A0101 dataset. Bold entries denote three species that were not detected by Fellows Yates *et al.* using the MALT approach. The values for sequence similarity and gene coverage correspond to the highest scoring alignment for each genus.

Additionally, we evaluated the assemblers’ ability to reconstruct contigs containing biosynthetic gene clusters (BGCs). Fig. 6 (panel **A**) indicates that the OAK samples are the most protein-rich. Accordingly, we selected the OAK003 sample for further analysis. Using antiSMASH [95], we screened the contigs from each assembler for BGCs and reported the number of protoclusters per BGC type. A protocluster is defined as a genomic region that contains the core biosynthetic genes for a specific secondary metabolite along with its neighbouring genes. Based on curated detection rules, each protocluster corresponds to a single product type [96].

Fig. 7 (panel **B**) shows that antiSMASH identified 11 distinct protocluster types across all assemblers. Both CarpeDeam modes contain the highest number of protoclusters, with 62 identified in safe mode and 52 in unsafe mode. MEGAHIT contains 25 protoclusters, PenguiN contains 10, and metaSPAdes contains 8. The distribution of protocluster types varied among assemblers. For example, MEGAHIT did not detect any *RRE-containing*, *lassopeptide*, or *terpene* protoclusters, which were present in the CarpeDeam assemblies.

Another notable difference was observed in the total number of aligned nucleotides. We searched the protein sequences reported by antiSMASH against the nr database and summed the alignment lengths of the best hits, determined by the lowest E-value. The safe mode of CarpeDeam assembled sequences with more than twice the total aligned length (in base pairs) compared to MEGAHIT. When aligning the BGC protocluster sequences against the nr database, the best hits had a median sequence similarity exceeding 92% for CarpeDeam, MEGAHIT, and metaSPAdes, while PenguiN showed a lower median similarity of over 83%. To ensure ancient authenticity, we mapped the OAK003 reads against the BGC-involved contigs, all of which exhibited characteristic nucleotide substitutions at the fragment ends (see PyDamage[93] results in the external data repository).

## 4 Discussion

### Evaluating Assembly Algorithms: Understanding Strengths, Limitations, and Underlying Assumptions

The reconstruction of full-length genes or entire genomes from ancient metagenomes poses significant challenges for *de novo* assemblers. *De novo* assembly of metagenomic samples is already complex for modern samples and it becomes exceptionally challenging with aDNA due to its degraded nature, characterized by extremely short fragment sizes and deaminated nucleotides.

While modern metagenomic assemblers like MEGAHIT [58] and metaSPAdes [59] are widely used, their de Bruijn Graph implementations face a precision-sensitivity trade-off when dealing with highly complex datasets [65, 62]. As graph complexity increases, these tools must employ simplification strategies, potentially compromising assembly completeness or accuracy, especially in regions of high genomic diversity or low coverage.

PenguiN[62] addresses this issue through a greedy iterative overlap approach, leveraging whole read information for improved performance in variant-rich datasets. Ancient metagenomic datasets, particularly those from poorly preserved samples, present additional challenges for both de Bruijn graph and overlap-based assembly strategies. Base substitutions in aDNA fragments artificially increase the genomic diversity in metagenomic samples. Deep sequencing can facilitate assembly [17], but it may be impractical due to sample limitations or cost constraints [6].

Our tool, CarpeDeam, builds upon PenguiN’s whole-read approach, incorporating a maximum-likelihood framework that utilizes sample-specific damage pattern matrices for contig correction and extension. Given the close relationship between CarpeDeam and PenguiN, with large parts of their algorithms sharing the same codebase, it is essential to clarify their distinctions.

Both assemblers share a foundational concept – leveraging the MMseqs2 linclust algorithm to cluster reads based on shared *k*-mers, yet they differ significantly in their suitability for assembling aDNA datasets. Overall, clustering by shared *k*-mers enables both tools to extend sequences in an overlap-based consensus approach. This method avoids an all-vs-all read comparison, which scales quadratically with the number of input reads and is thus computationally infeasible.

In both assemblers, a cluster is defined by sequences that not only share at least one *k*-mer with the cluster representative (the longest sequence) but also meet a sequence similarity threshold regarding the overlapping portions of two sequences. For PenguiN, this threshold is set to 99% by default, meaning that within the overlap of a member sequence and its cluster representative, at least 99% of the bases must be identical [62]. The assumption that sequences belonging to the same cluster are highly similar makes PenguiN particularly well-suited for assembling viral genomes or microbial 16S rRNAs from modern, non-ancient metagenomic samples, which it was specifically designed for. Furthermore, PenguiN uses a Bayesian framework to infer the most likely extension, given the observations of overlap length and sequence similarity. PenguiN’s framework relies on the premise that mismatches in overlapping reads are rare and primarily arise from sequencing errors. Sequencing errors, especially in modern Illumina datasets, tend to be minimal (*<* 0.1 - 0.6%) [97] and can be further reduced upstream of assembly by quality filtering reads before assembly [98]. Consequently, PenguiN does not include a base correction algorithm; it relies on overlaps between presumed high-quality reads for accurate assembly.

However, this design makes PenguiN unsuitable for aDNA assembly. aDNA fragments inherently contain postmortem base substitutions, which artificially inflate sequence diversity and disrupt assumptions about sequence identity. Without correcting these substitutions, PenguiN faces two critical challenges. First, damaged reads may fail to cluster because the sequence mismatches caused by deaminations reduce overlap identity below the required 99% threshold. Second, even if damaged reads do cluster and overlap, the accumulation of deaminated bases during iterative extensions can propagate errors, further hindering assembly. While it is possible to lower the sequence identity threshold, as we did with CarpeDeam, this alone does not address the issue of accumulating damage.

These challenges are addressed by CarpeDeam, which incorporates damage-aware correction, extension, and *RY*-mer sequence identity filtering to adapt PenguiN’s sensitivity for aDNA samples.

To maintain objectivity, we ran all assemblers in their default modes, as this reflects the typical approach researchers might take when applying these tools. However, we recognize that the performance of assemblers may vary depending on parameter settings, particularly for challenging datasets such as aDNA. In our case, we noticed that metaSPAdes encountered issues during the error correction step, which is designed to address sequencing errors rather than ancient DNA damage. To proceed, we followed the advice provided by the developers in a related GitHub issue (see Section 3, subsection “simulated data”, and ran metaSPAdes in “only assembler” mode as suggested. Given the unique characteristics of ancient DNA, a comprehensive parameter bench-marking study could provide valuable insights into optimizing assembly strategies. However, such an exploration is beyond the scope of the current manuscript.

### Significant Impact of Fragment Length Distribution and Damage Patterns on the Assembly Process

Our extended analysis of simulated aDNA samples, incorporating three fragment length distributions (medium, short, and ultra-short), three levels of DNA damage (moderate, high, and ultra-high), and three depths of coverage (3X, 5X, and 10X), revealed that all these factors significantly influence the *de novo* assembly of aDNA samples.

Modern *de novo* assemblers are generally designed to handle sequencing reads that, when merged upon assembly, are several hundred base pairs in length. In a paired-end sequencing context, read lengths typically range from 2×75 bp to 2×300 bp [99], providing a very different starting condition compared to aDNA samples. aDNA molecules are heavily degraded, resulting in short DNA fragments with read length distributions peaking well below 100 bp [100]. These distributions often follow an inverse exponential relationship between fragment length and abundance [101]. A commonly used metric, the median fragment length, serves as an informative proxy for estimating the most abundant read size, which is expected to significantly influence the assembly process.

Our simulations support this observation to some extent. Larger median fragment lengths and more uniform length distributions, such as those observed in the medium fragment length distribution used in this study (Supplementary Material, Fig. S1 (A)), positively impact assembly quality. However, we also observed that rare long fragments (*>*100 bp) seemingly have a disproportionate influence on assembly outcomes. Despite being a small minority, these long sequences appear to act as exceptionally useful anchors, enabling assemblers to produce more contiguous and less fragmented assemblies. This effect was observed in both de Bruijn graph assemblers, MEGAHIT and metaSPAdes, and was even more pronounced in the overlap-consensus assemblers, CarpeDeam and PenguiN.

Damage levels also had a significant influence on assembly quality. The three damage levels used in this study (moderate, high, and ultra-high) differ in two key aspects. First, the maximum damage at the first position of a fragment varies considerably, with substitution rates ranging from 0.33 in moderate damage to 0.58 in ultra-high damage. Second, the damage levels differ in their slope, representing how quickly substitution rates decline along the fragment. For example, while ultra-high damage exhibits a substitution rate of only 0.05 at position 5, moderate damage maintains a substitution rate of approximately 0.14 at the same position.

Interestingly, all assemblers struggled more with the moderate damage level and its lower decline rate compared to the ultra-high damage level, as demonstrated in the Supplementary Material, Section S2. This counterintuitive result can be explained by the mechanics of the assembly algorithms.

For de Bruijn graph-based assemblers, such as MEGAHIT and metaSPAdes, the extraction of *k*-mers as building blocks presents a challenge. These assemblers often start with *k* = 21 or higher [58, 59]. With already short fragment lengths, the number of *k*-mers that can be extracted is limited. For instance, in the case of a 40 bp read where the first and last 5 bases have relatively high substitution rates, only the central portion of the read contributes *k*-mers that accurately represent the original sequence. *k*-mers spanning damaged regions are more likely to contain mismatches, making it difficult to find matching *k*-mers from other reads, and thereby impeding graph extension.

Overlap-based assemblers, such as CarpeDeam and PenguiN, face a different set of challenges. Reads are clustered together based on shared *k*-mers, but the full overlap must meet a minimum sequence identity threshold (90% for CarpeDeam, 99% for PenguiN). For ultra-short reads with high substitution rates in the first few positions, the sequence identity can quickly fall below these thresholds. While lowering the threshold could mitigate this issue, it would significantly increase computational resource usage (due to larger clusters) and the risk of misassemblies.

These findings are critical for future studies utilizing *de novo* assembly for aDNA and highlight the importance of developing methods specifically optimized for highly fragmented and damaged DNA.

### Impact of Coverage Depth on the Assembly Process and the Issue of Misassemblies

The role of coverage depth in the assembly process has been risen in other studies as well. For instance, Klapper *et al.* [17] assembled ancient metagenomic datasets to identify biosynthetic gene clusters (BGCs), leading to the experimental validation of previously unknown molecules. Their findings showed that while modern metagenomic samples produced more contiguous assemblies than ancient samples, deeply sequenced ancient metagenomes achieved similarly high-quality assemblies [17].

Our simulations support this observation. Higher average coverage depths consistently resulted in better assemblies across multiple metrics. However, as detailed in Supplementary Material, Section S2, current assemblers still face challenges when dealing with the short fragment lengths and high levels of damage. For instance, while MEGAHIT often achieved genome fractions comparable to CarpeDeam, it underperformed in terms of largest alignment and NA50. This highlights the limitations of assemblers not specifically designed for aDNA, even under high-coverage conditions.

Another study by Jackson *et al.* [102] demonstrated as well that deeply sequenced samples could be successfully assembled with MEGAHIT. However, deep sequencing is often impractical due to sample limitations or cost constraints [6].

The improved performance of de Bruijn graph-based assemblers with high coverage is straight-forward to explain: higher coverage ensures that genome positions are more likely to be covered by *k*-mers free of deaminated bases, enabling more contiguous paths through the assembly graph. Theoretically, the same principle applies to overlap-based assemblers like CarpeDeam and PenguiN. While higher coverage datasets in our simulations led to improved assembly metrics overall, a significant misassembly rate could be observed. In unsafe mode, CarpeDeam exhibited particularly high misassembly rates, especially in high-coverage datasets.

This issue led to the introduction of the safe mode for CarpeDeam. In early assembly stages, short reads may overlap with 100% sequence identity, particularly in cases involving mobile genetic elements (MGEs) shared between species [103]. Under these conditions, the assembler cannot reliably distinguish sequences from different species. Graph-based assemblers handle this by marking such regions as diverging paths, effectively “cutting” contigs at these points, resulting in shorter but accurate assemblies. The safe mode of CarpeDeam addresses this issue by utilizing a consensus calling mechanism, thereby reducing misassembly rates severalfold, albeit at the cost of shorter contigs. Users can alternatively choose the unsafe mode, which disables the consensus calling mechanism, potentially increasing sensitivity but at the risk of producing more misassembled or chimeric contigs.

It must be noted that it might seem counterintuitive why CarpeDeam’s unsafe mode exhibited exceptionally high misassembly rates in high-coverage datasets, where higher coverage should theoretically improve assembly. We believe that this is triggered by sequences of highly abundant species. In some cases, clusters may contain reads that overlap perfectly with the center sequence at 100% identity. While the correct extension may originate from a less abundant species, the assembler is more likely to select a perfectly overlapping but incorrect sequence from the most abundant species, leading to misassemblies. A detailed investigation of this issue is provided in Supplementary Material, Section S4, where we are showing that identical sequences can occur within a single genome or between genomes of different species, with considerable variation across prokaryotic taxa [104, 103] (Supplementary Material Fig. S16 and Fig. S17).

Our observations highlight the trade-off between sensitivity and accuracy in metagenomic assembly, especially in the context of aDNA and diverse prokaryotic communities. The availability of safe and unsafe modes allows users to adjust the assembly approach based on their specific research goals and the nature of their datasets.

A potential future solution could involve incorporating a graph-like structure within the clusters sharing a *k*-mer, combining the overlap-based approach with the flexibility of graph assemblers. However, this approach requires further research. Downstream analyses—such as protein prediction or contig alignments against reference databases—can help identify misassembly breakpoints, enabling researchers to extract valuable insights from the contigs for further investigation.

### Protein-Centric Approaches in aDNA Research

We assembled 20 empirical datasets from 5 sample sites, two of which were further analysed for taxonomic classification. The results underscore the potential of protein prediction in aDNA research to uncover novel insights. Protein prediction becomes feasible only with sequences that are sufficiently long for ORF prediction, which is challenging when dealing with inherently short aDNA reads. As highlighted by Borry *et al.* [93], predicting ORFs directly from ancient DNA reads is extremely difficult, if not impossible, with current tools. By contrast, *de novo* assembly generates sequences long enough for ORF prediction.

Proteins offer unique advantages in aDNA research as they are more conserved than nucleotide sequences. Modern tools like MMseqs2 [76], and DIAMOND [105] enable rapid and sensitive screening of translated sequences against large protein databases. These advantages highlight the potential of protein-centric approaches for the discovery of functional elements and taxonomic profiling in aDNA datasets.

Gene- and protein-centric approaches have already demonstrated success in related fields. Klapper *et al.* [17] identified and experimentally validated bioactive molecules from biosynthetic gene clusters in ancient metagenome-assembled genomes. Wan *et al.* [50] applied deep learning to analyse proteomes of extinct organisms, identifying potential antimicrobial peptides effective against highly pathogenic bacteria. Such studies could benefit from advanced *de novo* assembly methods with higher sensitivity, such as those provided by CarpeDeam.

Our results confirm this potential but also reveal substantial variability in assembler performance across datasets. While CarpeDeam and MEGAHIT generally performed best, the ECO datasets presented an exception, with MEGAHIT outperforming CarpeDeam. This may be explained by the high variability in damage patterns within the ECO datasets. The damage patterns per sample are shown in the Supplementary Material, Fig. S19. The ECO datasets deviate most from the average damage profile used as input, whereas the EMN and GDN datasets show relatively small deviations. Incorporating species-specific damage patterns could significantly enhance assembly performance, but implementing this approach would be challenging as it requires substantial modifications to the tool’s core code base. While our method yields superior results in many cases, these findings highlight that no single assembler performs best across all datasets, emphasizing the importance of employing diverse assembly strategies. We recommend that assembling large environmental samples should include careful preprocessing to remove non-damaged contaminants and minimize extreme variance in ancient sequence damage profiles.

Although protein-based taxonomy profiling is not yet widely used in aDNA research, it remains a promising alternative to nucleotide-based approaches, which have their own challenges, as mapping methods face significant limitations due to the low mapability of short reads in metagenomic datasets [41, 106]. By leveraging the conserved nature of amino acid sequences, protein-based taxonomy offers the potential for more accurate assignments. Our taxonomic classification results showed that both CarpeDeam and MEGAHIT identified a substantial number of unique taxa not detected by other assemblers. This result demonstrates that further development of assembly strategies holds significant potential for uncovering additional biological insights from existing datasets.

### Further Applications: 16S rRNA-Based Taxonomic Assignment and Exploration of Biosynthetic Gene Clusters

Sequencing of 16S rRNA genes is widely used for phylogenetic analyses to discriminate taxa at various levels [107, 108]. Sequencing the full-length ( 1500 bp) gene provides the most accurate classifications [94]. Targeting specific hypervariable regions of the 16S rRNA gene has also been explored for phylogenetic analysis [109]. Previous efforts in ancient microbiome reconstructions [83, 84, 85, 86, 87, 88] have encountered challenges due to ancient DNA fragmentation—often with lengths rarely exceeding 200 bp—which hampers the sequencing of long hypervariable regions such as the V4 region [89, 110]. In contrast, shotgun metagenomics, widely employed for profiling complex microbiomes, can yield increased false positives and negatives because of the high species diversity in environmental samples. Although Eisenhofer *et al.* [110] contend that shotgun metagenomics is best suited to capture the full spectrum of microbial diversity, we propose that *de novo* assembly can be leveraged for refining taxonomic specificity. However, reconstructing 16S rRNA genes is inherently challenging because their highly conserved regions often extend beyond the typical *k*-mer length used by de Bruijn graph assemblers, resulting in highly branched graphs and fragmented assemblies [62, 111, 112]. Despite these challenges, our approach successfully assembled contigs that align with curated 16S rRNA gene databases with high sequence similarity.

We compared the taxonomic classification from the study by Fellows Yates *et al.*, which used read mapping on shotgun-sequenced samples, with our 16S rRNA *de novo* assembly approach applied to the same sample. Although we did not recover all the taxa reported by Fellows Yates *et al.*, our method successfully identified three full-length 16S rRNA genes at species-level that were absent in their analysis. Thereby, we highlighted the complementary potential of *de novo* assembly to mapping approaches for phylogenetic analysis.

Our analysis of the assemblers’ ability to reconstruct genes within BGCs indicated that CarpeDeam increases sensitivity by identifying both additional BGC types not detected by other assemblers and a higher overall number of BGC candidates. This enhanced performance was further supported by longer alignment lengths of BGC genes recovered by CarpeDeam. However, as shown by Klapper *et al.*, deeply sequenced samples are preferable for BGC analyses, with detection being particularly effective in datasets that yield contigs of at least 10 kb.

### Limitations of CarpeDeam

CarpeDeam uses a greedy iterative approach, which may not be as time-efficient as the de Bruijn graph methods employed by assemblers such as MEGAHIT and metaSPAdes (see runtime and memory usage in Supplementary Material, Section S7). However, CarpeDeam represents a valuable alternative to the long-established dominance of de Bruijn graph algorithms, demonstrating its potential to outperform these methods in specific benchmark metrics. While de Bruijn graphs have been the foundation of *de novo* assembly for decades [113], our results highlight the promise of alternative strategies. The greedy overlap-based approach, initially introduced by the protein-level assembler PLASS, adapted by the protein-guided nucleotide assembler PenguiN, and further refined in CarpeDeam, offers increased sensitivity, uncovering sequences that remain undetectable by other assemblers.

A unique feature of CarpeDeam is its use of a damage matrix, introducing a novel concept in aDNA assembly. The assembler currently applies a global damage model, which does not account for differences in damage across sequences within a sample. Future updates could include species-specific damage profiles to better reflect the diverse damage patterns seen in metagenomic datasets. Despite this limitation, CarpeDeam has successfully assembled empirical datasets with damage variation within a limited range of ±10%, demonstrating its applicability to ancient metagenomic samples.

CarpeDeam’s primary limitation is its tendency to assemble a higher fraction of misassemblies. This issue, along with potential solutions, was discussed earlier, including the introduction of safe and unsafe modes to mitigate the problem. While the safe mode reduces misassemblies, it raises the question of which mode is best suited for a given dataset, as results can vary. For instance, our investigation of Non-Misassembled Contigs (see Section 3 and Supplementary Material Fig. S18) revealed that in 8 out of 9 datasets, the unsafe mode produced more non-misassembled contigs than the safe mode. This is due to a key feature of both PenguiN and CarpeDeam, which allows single reads to be used multiple times during assembly, potentially creating redundant contigs. Redundancy is reduced during the final clustering step, where sequences are grouped at 97% sequence identity, but this process also allows any sequence to become a cluster’s center sequence if it is the longest and shares a *k*-mer with at least one other sequence. Consequently, the unsafe mode generates more non-misassembled contigs, but also more misassemblies, leading to a higher “misassemblies per contig” ratio. In contrast, the safe mode aborts extensions when potential misassembly origins are detected, resulting in shorter contigs and fewer non-misassembled contigs larger than 2000 bp. Ultimately, the choice between modes depends on the downstream analysis. For instance, researchers prioritizing sensitivity in aDNA assembly might prefer the unsafe mode, albeit with the need for more careful downstream filtering (e.g., alignment-based methods), while others may favour the safe mode for its higher robustness across assembled contigs.

The analysis of protein sequences derived from the assembled contigs revealed that CarpeDeam could recover more unique proteins compared to other assemblers, as illustrated in Figure 5, panel **(A)**. Additionally, CarpeDeam demonstrated a higher aligned fraction of amino acid residues belonging to high-confidence aligned proteins, as shown in **panel (B)**. However, the differences between MEGAHIT and CarpeDeam in protein recovery were not as pronounced as the mapped fraction of nucleotide base pairs suggested (Figure 4 panel **(A)**). For instance, CarpeDeam’s assembly in unsafe mode mapped more than twice as many nucleotides of non-duplicated base pairs against the reference compared to MEGAHIT. This discrepancy between nucleotide mapping and protein recovery indicates that a notable proportion of potential protein-coding sequences are not accurately predicted, likely due to frame-shifts, indels, or erroneous base corrections. Such errors in the assembled sequences can disrupt reading frames or introduce premature stop codons, leading to incomplete or missed protein predictions. This observation highlights that while CarpeDeam’s damage correction algorithm demonstrates significant improvements over existing methods, there remains potential for further refinement.

## 5 Conclusion

CarpeDeam represents a significant advance in aDNA assembly as the first sample-specific, damage-aware assembler. Our experiments on both simulated and empirical data demonstrate CarpeDeam’s ability to improve the assembly of long, continuous sequences from degraded aDNA. By incorporating a maximum-likelihood framework that utilizes sample-specific damage pattern matrices, CarpeDeam outperforms existing assemblers in recovering genes and RNA sequences, from metagenomic samples. The tool’s dual operational modes - safe and unsafe - offer users flexibility in balancing sensitivity and accuracy based on their research goals. While challenges remain in reconstructing full-length genomes from highly degraded samples, CarpeDeam’s improved recovery of RNA content highlights its potential for taxonomic classification and functional analysis, particularly in ancient microbiomes. As the first damage-aware metagenome assembler, CarpeDeam lays the groundwork for future innovations in this field. Looking ahead, combining CarpeDeam’s novel approach with established de Bruijn graph methods could potentially lead to even more robust and accurate aDNA assembly techniques, further advancing our ability to reconstruct and analyse ancient microbial communities.

## 6 Methods

### Test Data

To assess the assembly performance of ancient metagenomic samples, we simulated three distinct metagenomic environments. Each environment varied in complexity, including different abundance profiles of species. The taxonomic composition was modeled on previously published studies from Granehall *et al.* [35], Wibowo *et al.* [44], and Der Sarkissian *et al.* [69]. We simulated data with three levels of average coverage (3X, 5X, and 10X), three fragment length distributions (medium, short, and ultra-short) and three levels of damage (moderate, high, and ultra-high). In total, 81 datasets were generated. Table 5 provides an overview of representative datasets with short fragment length and moderate damage.

**Table 5:** Characteristics of simulated ancient metagenomic datasets representing three distinct environments: Bone, Dental Calculus, and Gut. The table summarizes the number of taxa, average coverage depth, number of sequences, total sequence length, and GC content (%) for representative datasets simulated with short fragments and moderate damage.

| Sample | # Taxa | Coverage | # Reads<br>(merged) | Total<br>Length | GC<br>Content (%) |
| --- | --- | --- | --- | --- | --- |
| Bone | 34 | 3X | 13,481,538 | 689,950,613 | 57.62 |
| Bone | 34 | 5X | 22,469,438 | 1,149,702,991 | 57.62 |
| Bone | 34 | 10X | 44,938,811 | 2,299,239,612 | 57.62 |
| Dental<br>Calculus | 59 | 3X | 8,169,233 | 417,738,204 | 43.77 |
| Dental<br>Calculus | 59 | 5X | 13,615,501 | 696,358,011 | 43.76 |
| Dental<br>Calculus | 59 | 10X | 27,231,047 | 1,392,706,548 | 43.77 |
| Gut | 116 | 3X | 25,583,926 | 1,308,958,512 | 42.45 |
| Gut | 116 | 5X | 42,640,124 | 2,181,743,052 | 42.45 |
| Gut | 116 | 10X | 85,279,973 | 4,363,114,755 | 42.45 |

The fragment length distributions used in the simulations were labeled medium, short, and ultra-short, with the medium distribution derived from the EMN001.A0101 sample (Fellows Yates *et al.* [36]), the short distribution from the Vi33.19 sample (Gansauge *et al.* [114]), and the ultra-short distribution from the OAK001.A0101 sample (Fellows Yates *et al.* [36]). The distributions as well as the damage level categories are shown in the Supplementary Material, Fig. S1. We used gargammel [115] to simulate paired-end reads for the bacterial communities using the aforementioned damage patterns and fragment length distributions. Adapters were trimmed, and paired-end reads were merged with leeHom [116] using the--ancientdna option. The simulated reads were aligned to the reference genomes with Bowtie2 [71], and the resulting BAM files were processed with DamageProfiler [9]. The estimated damage patterns were then used as input for CarpeDeam, which incorporates these patterns into its maximum likelihood framework. This workflow avoided any bias in favour of CarpeDeam, as the simulation closely reflects how damage patterns are realistically derived.

The empirical datasets we benchmarked the assemblers against were obtained from Fellows Yates *et al.* [36]. Fellows Yates *et al.* provided extensive analysis of the species found in these datasets, which made them highly suitable for our analysis, as we could utilize the presented damage patterns for the input of CarpeDeam. The damage patterns observed in these datasets varied across species. This variability allowed us to infer CarpeDeam’s robustness against varying damage levels, as it uses a single damage profile for all input data. An overview of the C→T substitution frequencies for identified species can be found in the Supplementary Material, Fig. S19.

### Benchmark Workflow

#### Evaluation of Simulated Metagenomic Datasets

The assembly performance was evaluated using metrics reported by metaQUAST [70]. For all 81 simulated datasets, we ran metaQUAST and mapped the reads against the assemblies, providing the reference genomes used for the simulations. The minimum sequence identity for alignments was set to 90%, as suggested in the metaQUAST documentation. While the full metaQUAST results for all datasets can be found in the Supplementary Material, the results in the main manuscript focus on a subset of the simulations: three levels of coverage (3X, 5X, and 10X) for the three different environments (Gut, Dental Calculus, and Bone) simulated with moderate damage and short fragment length distribution. Informative metrics for this subset are reported in Fig. 3 and Table 1, with additional tables for dental calculus and bone simulations available in the Supplementary Material (Tables S2 and S3). For the alignment evaluation, metaQUAST uses the pairwise sequence aligner minimap2 [58].

To complement the genome fraction metric and provide a more comprehensive assessment of assembly quality, we evaluated the ability of each assembler to reconstruct longer, correct contigs. Specifically, we quantified the number of correct contigs of length 2000 bp or longer for each assembler across a subset of the datasets. This analysis addresses a limitation of the genome fraction metric, which does not account for the length distribution of contigs mapping to the reference genome (Supplementary Material Fig. S18).

We further assessed the assemblers’ performance in reconstructing protein content. This analysis involved annotating the produced contigs and searching for similar proteins using Prokka [75], which employs Prodigal [117] for open reading frame prediction and searches against three core databases.

We utilized the MMseqs2 map [76] module in--cov-mode 1 with the--min-seq-id parameter set to 0.9 to find very similar sequence matches with at least 90% sequence identity and covers the target by at least 90%. We then reported the number of unique matches in the reference (Fig. 5, panel **(A)**).

In panel **(B)** of Fig. 5, we analysed the fraction of amino acid residues in the assembled proteins that aligned to the predicted reference proteins. Again, we used Prokka [75] to predict proteins from the assembled contigs. We then searched these predicted proteins against the reference proteins using the MMseqs2 map module. We set the minimum coverage parameter -c to 0 and the --min-seq-id parameter to 0.9, to report any alignments with at least 90% sequence identity. From these high-confidence alignments, we counted the number of amino acid residues for each assembled protein. We then calculated the fraction of mapped amino acid residues compared to the total number of residues in the assembled proteins. This fraction of aligned amino acid residues is shown in panel **(B)** of the figure.

### Taxonomy Assignment of Contigs and Protein Similarity Search of Empirical Datasets

For two empirical datasets, EMN001.A0101 and GDN001.A0101, we conducted a taxonomic assignment analysis using the assembled contigs. Since we did not have the ground truth for the empirical datasets (only selected identified species from Fellows Yates *et al.* [36]) we employed different evaluation methods. First, we used the MMseqs2 taxonomy module [34] for taxonomic classification. This module translates contigs in all six reading frames and searches the amino acid sequences against a target database. For our target, we used proteomes that are representatives for species according to the Genome Taxonomy Database (GTDB) [80]. These proteomes are the cluster representatives for each species, and this database was downloaded using the databases module of MMseqs2 [76]. The GTDB is a phylogenetically consistent, genome-based taxonomy that uses a large number of quality-controlled genomes to infer evolutionary relationships among bacteria and archaea.

We analysed the number of unique species assigned to the translated contigs, considering all species where at least one translated contig covered a single-copy protein by at least 90%. We then examined the number of shared taxa and unique taxa per assembler (see Fig. 6), panel **(B)**.

Additionally, we analysed the capability of the assemblers to assemble protein fractions that can be used for identifying evolutionary related proteins. We predicted open reading frames (ORFs) from the contigs generated by all assemblers using Prodigal [117] with the “meta” setting, considering only full-length ORFs for further analysis. We then used MMseqs2’s easysearch module [76] to search the amino acid sequences against the UniProt100 database [118], filtering the results by a minimum sequence identity of 0.35, an E-value smaller than 1e-5, and a minimum alignment length of 100 residues. The number of unique protein hits is reported in Fig. 6, panel **A**.

### rRNA Detection and BGC Screening

We analysed the ability of the assemblers to reconstruct 16S rRNA genes, which are key markers for taxonomic classification. Assembled contigs were annotated for RNA sequences using Prokka [75], and the annotated RNA sequences were clustered at 99% sequence similarity and a minimum coverage of 99% to reduce redundancy with MMseqs2 linclust [68]. The clustered RNA sequences were then searched against the SILVA database [119], which includes 510,495 high-quality, full-length 16S and 18S rRNA sequences. To focus specifically on bacterial origin, we filtered the database to retain only bacterial rRNA genes, resulting in a subset of 432,033 sequences.

To ensure specificity, the results were filtered to retain only sequences that covered at least 80% of a database entry. The number of unique hits in the SILVA database, spanning sequence similarity thresholds from 90% to 100%, is shown in Fig. 7, panel **A**.

For BGC screening, we used antiSMASH 7.1 [95] to identify biosynthetic gene clusters by searching against the antiSMASH database. Subsequently, MMseqs2 [68] was employed to search the BGC sequences against the nr database, allowing us to evaluate sequence similarity and alignment length.

### Deamination Correction in Assembly Process

The assembly workflow incorporates a correction stage for deaminated bases at the start of each iteration. First, sequences are clustered based on shared *k*-mers, following PenguiN. The choice of *k*-mer size impacts clustering accuracy, with larger *k* values generally producing clusters that more reliably group sequences of common origin. Given the typically short length of aDNA fragments, a default *k*-mer size of 20 is set. The clusters are characterized by:

- A center sequence, which is either a contig (an extended sequence not found in the raw input data) or an aDNA fragment. It is the longest sequence in the cluster.
- All remaining members of the cluster are aDNA fragments (non-extended fragments).
- All members of a cluster fulfill threshold criteria such as sequence identity and E-value (as in PenguiN) and the newly introduced RYmer sequence identity, which exclusively counts mismatches between purines and pyrimidines, thereby tolerating mismatches due to deamination, i.e. C→T or G→A substitutions.

The goal is to correct the center sequence of each cluster, using alignments from member sequences and user-provided damage patterns that estimate base substitutions in the input data.

Consider a cluster containing a center sequence *C* and at a given position in this sequence, we seek to find the most likely base *b_C_* given the data. The data consists of a set of aDNA fragments R aligning at that position. The fragments in R share at least one *k*-mer with *C*. More prosaically, we seek *P* [*b_C_* | R] which can be obtained by finding *b_C_* which maximizes the expression *P* [R | *b_C_*] assuming a uniform prior for all 4 DNA nucleotides. The independence assumption allows us to say:

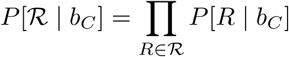

For a base *b_R_*in *R* at that position, each factor is defined as:

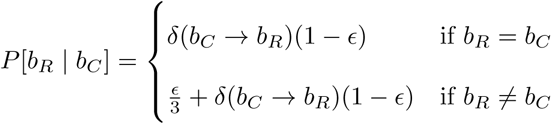

Here *ɛ* represents the sequencing error rate and is set at 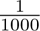 for all bases as PenguiN does not store the quality scores at this stage. The term *δ*(*b_C_* → *b_R_*) represents the rate of substitution from C to R due to aDNA damage as provided by the user for this particular position in the fragment. To illustrate, if the rate of C to T at the 5’ end is 30%, then *δ*(^′^*C*^′^ →^′^ *C*^′^) = 0.7 and *δ*(^′^*C*^′^ →^′^ *T* ^′^) = 0.3. In the case of the bases not matching, we consider the possibility of either sequencing error or damage. If the bases match, then no damage has occurred and the base was called correctly. We neglect the possibility of a deamination followed by a sequencing error which would result in the same base as it is dwarfed by the other term.

### Assembly with aDNA fragments

#### Selecting the extension candidate

Following the initial correction of the center sequence in each cluster, the workflow proceeds with a damage-aware extension process. This involves extending the center sequences of each cluster using sequences that are part of the cluster that share at least one *k*-mer with the center sequence and meet additional criteria such as minimum sequence identity, E-value, and RYmer sequence identity. In this phase, only member sequences that are fragments rather than assembled contigs are considered, as the provided damage patterns are only applicable to sequences not yet merged as contigs. The selection of the most suitable sequence for extension is based on the overlap length and the number of matches within this overlap. However, as deamination events occur in aDNA, potential mismatches between the center sequence and the extension candidates are introduced. Therefore, the identification of the optimal sequence for extension is challenging.

Assuming that all bases in the center sequence *C* are correctly identified, we can calculate the likelihood of observing a specific overlap between *C* and an aDNA fragment *R_i_*, conditional on the damage pattern *δ*. This approach accounts for potential deamination events, such as observing a base ‘G’ in the center sequence and ‘A’ in the overlapping fragment, without classifying them as highly improbable, depending on the damage pattern.

The question is which *R_i_* is the best candidate for extension. More specifically, we seek *R_i_* which maximizes *P* [*R_i_* | *C*]. Let us define *b_C_* as the base in the center sequence and *b_Ri_* in the fragment. We have a nuisance parameter *b_δ_* which is the base post damage.

The likelihood for a single nucleotide *b_Ri_* of fragment *R_i_* is given by:

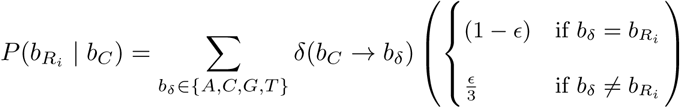

again, *ɛ* is 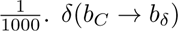 is the rate of nucleotide substitution due to aDNA damage.

However, not every aDNA fragment overlaps with the same amount of bases with *C*. Only computing the overlapping portions would put longer overlaps at a disadvantage. Let us say that the longest overlap among the candidates for extension is *l_max_* bp long. If *l_Ri_ < l_max_* counting the *l_max_* − *l_Ri_* missing bases as mismatches would be unfair to shorter overlaps as even random sequences will be 25% similar. We therefore count the probability of the missing bases as *K*^(^*^lmax^*^−^*^lR^i* ^)^. *K* is a constant between 0 and 1, representing the probability of a random match between two bases. The value of *K* is set by default at 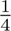.

### Stopping criterion

After identifying the extension candidate with the highest likelihood, this candidate is merged with the center sequence. This process continues iteratively, incorporating aDNA fragments that meet the sequence identity criteria, until no further suitable candidates remain. The extension process is terminated if no candidates satisfy the sequence identity threshold, or if the likelihood is significantly lower than the likelihood for a random match, given that it satisfies the minimum sequence identity threshold in the overlap (e.g. 90%). The likelihood of an alignment given the damage pattern is calculated, as well as the likelihood of observing a random match. The ratio of these two likelihoods determines the termination of the extension and can be set by the user. The minimum ratio used as threshold for accepting an extension was established through empirical testing to optimize the balance between sensitivity and precision. In Fig. S20 in the Supplementary Material the influence of different damage levels on the odds ratio threshold is visually modeled. The likelihood of observing the alignment by chance is calculated by multiplying the probability of a random match. Since every member sequence aligns to the center sequence with a user-defined sequence identity threshold, we use these values for a random match. For instance, if the threshold is set at 90% and the longest overlap is 20 bases, then the random model will have a likelihood of 0.9^20^.

This likelihood ratio approach allows for a relatively low sequence identity threshold (e.g., 90%) in the first phase which includes clustering sequences based on shared *k*-mers and calculating the overlap alignment, facilitating the inclusion of highly deaminated sequences. At the same time, it ensures an early termination of the extension process to guard against erroneous extensions.

### Damage-aware Rule for Merging Contigs

The extension process is divided into two distinct phases: an initial extension phase that exclusively considers non-extended aDNA fragments (damage patterns are only valid for unmodified sequences), followed by a second extension phase dedicated to merging contigs in the same cluster.

This second phase of extension focuses on merging contigs, allowing for a higher sequence identity threshold (98-99%) due to the correction of contigs in the previous phase. This threshold aligns with the default values in PenguiN. However, the potential presence of residual uncorrected bases, i.e. due to very low coverage, requires a modification of the extension strategy. However, we could not use the same strategy as described in Section 6, because firstly, the damage matrix becomes inapplicable to previously extended contigs due to the loss of positional information. Secondly, the significant variation in overlap length when merging contigs demands different penalizing strategies for shorter overlaps. For example, the contig merging step may encounter extension candidates with overlap lengths ranging from a few hundred to several thousand base pairs.

We modified PenguiN’s original extension rule (see Supplementary Note 1 in Jochheim *et al.* [62]) to score residual uncorrected bases. The original Bayesian approach in PenguiN’s algorithm models the extension process using a beta-binomial distribution, considering alignment length and mismatch count within overlaps. The extension of contigs stops when there are no longer any contigs matching above the identity threshold.

In the original rule, a match is assigned a value of 1 and a mismatch a value of 0. However, we have modified the value returned for mismatches between either C and T or G and A. The idea being that an extension candidate might have a mismatch due to an uncorrected base. Therefore, the sequence similarity of the overlap will be lower than it would be if the mismatch had been correctly corrected.

Given an observed mismatch of either C and T or G and A, we established a scoring function *f* (*s, ϕ*(*L*)*, δ*), that is:

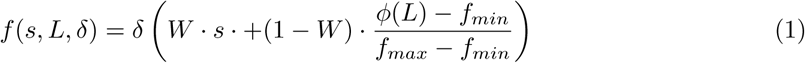

where:

- *L* is the length of the alignment
- 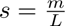 is the sequence similarity, with *m* being the number of matches in the alignment
- *ϕ*(*L*) is a scaling factor that accounts for the overlap length, which is based on a power law described by Sheinman *et al.* [104].
- *δ* is an approximation for the substitution rate. As the positional context is lost, we set it as the average damage rate for a particular base pair found in the original damage matrix.

Here, *W* is a weighting constant that balances the influence of the sequence similarity and the length-dependent factor. The value of *W* is set to 0.5. The function *ϕ*(*L*) is motivated by the work of Sheinman *et al.* [104], describing the relative frequency of identical sequences found among bacterial genomes depending on their length. We utilize the power law to evaluate an overlap being genuine by scoring the overlap length:

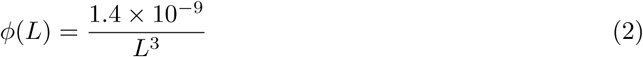

For practical implementation, we establish boundary conditions for the scaling factor *ϕ*(*L*):

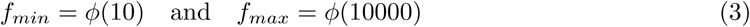

These constraints indicate that an overlap of 10 base pairs provides minimal confidence that two sequences belong to the same genome, while an overlap of 10000 base pairs or more provides maximum confidence.

Thereby, we aim to account for residual damaged bases in the contig merging step, ensuring that extension candidates with short overlaps but very high sequence identity, and candidates with long overlaps but lower sequence identity due to residual damages, are not disproportionately favoured.

### Misassembly Prevention by Consensus Calling: Introducing two modes of CarpeDeam

CarpeDeam’s algorithm relies on clustering sequences based on shared *k*-mers, utilizing the algorithm from the MMseqs2 [76] *k*-mers matcher module. As described by Jochheim *et al.* [62], this approach enables the use of whole sequence overlaps, providing an advantage over de Bruijn graph assemblers that use fixed-size *k*-mers, losing the context of the aDNA fragment information.

For modern samples with low sequencing error rates ( ≤1% [120]), PenguiN can apply a high default sequence similarity threshold of 99% between the center sequence and any member sequence, ensuring highly specific clusters. However, aDNA samples require a more lenient approach due to deaminated bases appearing as mismatches during clustering. To mitigate this issue while still excluding unsuitable candidates for extension, we implemented an additional threshold: the RYmer sequence identity. This method examines overlaps in Purine-Pyrimidine space, allowing mismatches between C and T as well as G and A, but restricting other mismatches. We also considered modifying the clustering algorithm to cluster sequences based on RYmers instead of *k*mers. This would enable clustering impervious to damage. However, the extremely short nature of aDNA fragments prevented us from increasing the RYmer size sufficiently to achieve the specificity required for effective clustering and extension. We demonstrated this by plotting the fraction of unique *k*-mers and RYmers of different sizes found in 1000 different bacterial genomes. The results can be found in the Supplementary Material, Fig. S21.

In practice, we applied an RYmer sequence identity threshold of 99.9% and a sequence identity threshold of 90%, permitting a certain level of Purine-Purine or Pyrimidine-Pyrimidine mismatches while minimizing other mismatches. Despite these measures, we observed a higher rate of misassemblies, particularly emerging in early extension phases where sequences are still very short (≤200bp), as these sequences lack sufficient specificity for clustering while allowing for deamination-induced mismatches. Investigation of the higher rate of misassemblies observed with CarpeDeam can be found in the Supplementary Material, Section S4.

To address these challenges, we developed a consensus-based approach that independently evaluates the left and right extending regions of all candidate sequences. These extending regions are the areas not covered by the center sequence but present in overlapping extension candidates. For each side (left and right), we construct a consensus sequence by considering all extension candidates. The consensus is built base-by-base, requiring at least five extension candidates to cover each position. This method results in two consensus sequences—one for the left extension and one for the right extension. The new center sequence is then formed by concatenating the left consensus, the original center sequence, and the right consensus. A schematic representation of this approach is shown in in the Supplementary Material Fig. S22. This approach is referred to as “safe mode” in the manuscript and is used as the default mode in CarpeDeam. It can be changed to “unsafe mode” by setting the parameter flag--unsafe to 1.

By aligning each member sequence to the consensus sequence and applying the updated overlap information to both the fragment extension and contig merging steps, we successfully reduced the level of misassemblies. However, this improvement came at the cost of a slight decrease in sensitivity. This trade-off resulted in the development of two distinct modes for CarpeDeam, offering users flexibility in balancing assembly accuracy and completeness based on their specific research requirements.

## 7 Declarations

### Data Availability

The simulated reads for the analysed datasets in this publication, along with additional files, are accessible on Zenodo under the DOI: 10.5281/zenodo.13208898.

### Code Availability

The software is publicly available at https://github.com/LouisPwr/CarpeDeam. All scripts and benchmark data for assembly and analysis presented in this manuscript are available at https://github.com/LouisPwr/CarpeDeamAnalysis.

### Software Versions

PenguiN 4-687d7 (GitHub clone from Jan. 2024), MEGAHIT v1.2.9, metaSPAdes v3.15.5, metaQUAST v5.0.2, Bowtie2 2.3.5.1-6, SAMtools 1.19, seqkit 2.6.1, prodigal V2.6.3, MMseqs2 Release 15-6f452, Prokka 1.14.6, skani 0.2.2, miniprot 0.13-r248

## Acknowledgements

We would like to acknowledge that this research and the PhD scholarships of Louis Kraft were funded by the Novo Nordisk Data Science Investigator grant number NNF20OC0062491. We thank the Technical University of Denmark’s Department of Health Technology for additional funding. Additionally, we would like to thank Joshua Rubin and Nicola Vogel for their thoughtful comments on our manuscript. The authors thankfully acknowledge the computer resources and the technical support provided by the BMBF-funded de.NBI Cloud within the German Network for Bioinformatics Infrastructure (de.NBI) (031A537B, 031A533A, 031A538A, 031A533B, 031A535A, 031A537C, 031A534A, 031A532B). AF-G is supported by a grant from the Danish National Research Foundation (DNRF174). L.K. acknowledges the use of OpenAI/ChatGPT and Anthropic/Claude to proofread the manuscript.

## Author Contributions

G.R. conceived the project. L.K. and G.R. designed and developed the method. L.K. implemented the method and tested it. L.K. conducted all tests. L.K. and A.F.G simulated datasets. A.F.G. provided expertise, computational resources and benchmark workflows and conducted additional analyses. J.S. supported the method development. M.S. and A.J. contributed software expertise and guidance in addressing data-centric issues. P.W.S provided IT infrastructure expertise. L.K. and G.R. wrote the manuscript. All authors approved the final manuscript.

