## Supplementary Material for "CarpeDeam: A *De Novo* Metagenome Assembler for Heavily Damaged Ancient Datasets"

[6]International Max-Planck Research School for Genome Sciences (IMPRS-GS)

[7]Campus Institute Data Science (CIDAS), University of Göttingen, Germany

[8]School of Biological Sciences, Seoul National University, Seoul, South Korea

[9]Artificial Intelligence Institute, Seoul National University, Seoul, South Korea

[10]Institute of Molecular Biology and Genetics, Seoul National University, Seoul, South Korea

[†]These authors contributed equally to this work.

### Contents

**S1 Simulated Datasets: Additional Information** 3

**S2 Benchmark Workflow** 3

**S3 Redundancy in Assembly and its Effect on Protein Analysis** 13

**S4 Non-uniform Simulated Datasets Assembly and Investigation of Misassemblies** 13

**S5 Number of Non-Misassembled Contigs per Assembler** 20

**S6 Damage Patterns in Empirical Datasets** 20

**S7 Algorithm Details** 21

**S8 Memory and Runtime** 24

### S1 Simulated Datasets: Additional Information

We simulated metagenomic datasets of different complexity to test CarpeDeam's performance against commonly used metagenome assemblers.

The taxonomies and their abundances were derived from previous studies and were used to simulate paired-end Illumina sequencing short reads with three different levels of DNA damage, three fragment length distributions, and three depths of coverage. The simulated damage levels were named moderate, high, and ultra-high, while the fragment length distributions were labeled medium, short, and ultra-short. The fragment length distributions are shown in Fig. S1 **(A)**: the medium distribution was derived from the EMN001.A0101 sample (Fellows Yates *et al.* [1]), the short distribution from the Vi33.19 sample (Gansauge *et al.* [2]), and the ultra-short distribution from the OAK001.A0101 sample (Fellows Yates *et al.* [1]). The damage patterns used in the simulations are detailed in Fig. S1 **(B)**.

For each combination of parameters, we generated reads with average depths of coverage of 3X, 5X, and 10X. We performed these simulations using gargammel [3]. We then used leeHom [4] to remove adapters and merge the sequences. As metaSPAdes requires both merged and unpaired reads as input, we trimmed the adapters from the raw, unpaired reads using adapterremoval [5]. Upon that, we estimated the damage from the simulated reads with DamageProfiler [6], which served as input for CarpeDeam.

The first metagenomic dataset was derived from Der Sarkissian *et al.* [7] including 34 taxa. The samples stem from bones and teeths and were pooled together. The second dataset mirrors the taxonomic composition of sample 2095 from Granehaell *et al.* [8]. It includes 59 taxa from ancient teeth. The third and most complex dataset is derived from Wibowo *et al.* [9] and contains 116 different species representing an ancient sample from the human gut.

All taxonomic identifiers and their abundances are listed in Supplementary File 1. All species belonged to either the kingdom of bacteria or archaea.

### S2 Benchmark Workflow

All scripts and benchmark data for assembly and analysis presented in this manuscript are available at https://github.com/LouisPwr/CarpeDeamAnalysis

Additional results for evaluating the assembly performance of the assemblers CarpeDeam, PenguiN [10], MEGAHIT [11] and metaSPAdes [12] as reported by metaQUAST [13]. For details on the NA50 and LA50 statistics, see the metaQUAST manual.

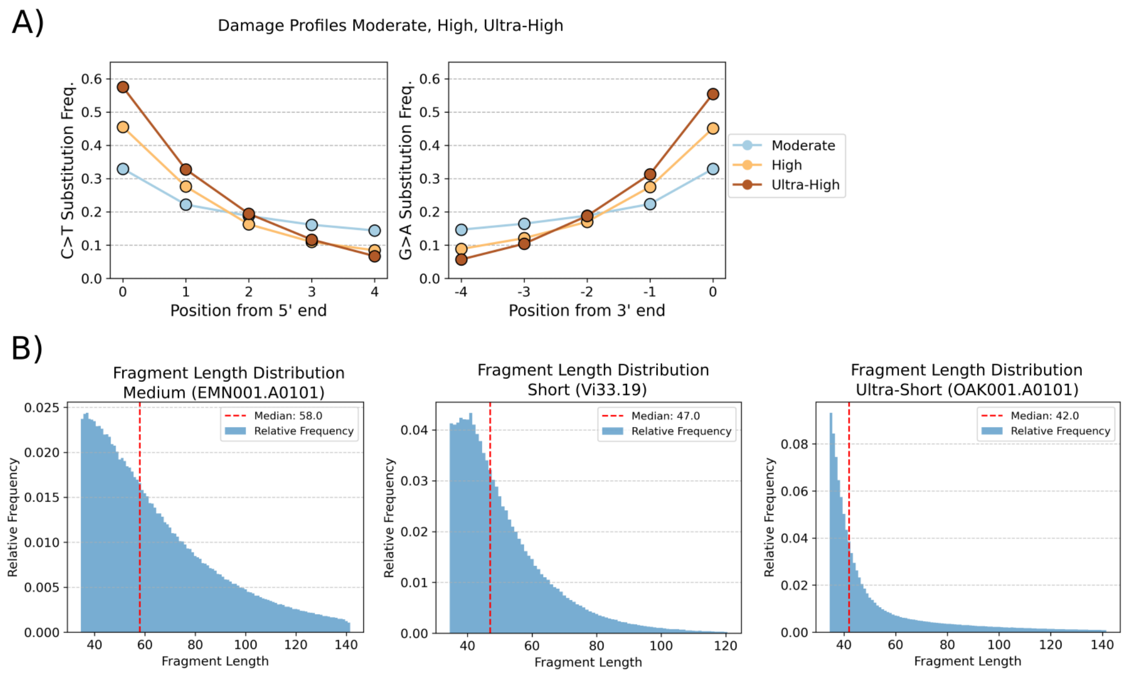

**Figure S1**: **(A)** Three damage profiles for simulated ancient DNA datasets: `moderate`, `high`, `ultra-high` Shown are the damage patterns for the first and last five positions. **(B)** Three fragment length distributions used for simulations: `medium`, `short` and `ultra-short`

The simulation of reads with three damage levels, three fragment length distributions, three coverages, and three environments resulted in a total of 81 assemblies. Their evaluations are presented as heatmaps. For each environment-gut, dental calculus, and bone-we include one plot for each of four key metrics reported by metaQUAST [13], resulting in 12 plots in total. Each plot visualizes the 27 parameter combinations for all assemblers.

#### Genome Fraction

The first three heatmaps (Fig. **S2**, Fig. **S3** and Fig. **S4**) show the genome fraction recovered by each assembler. We observe that higher average coverage depths consistently improve assembly quality, as indicated by an increased recovered genome fraction across all assemblers. Notably, in many cases, the recovered genome fraction in the 10X datasets is more than double that of the 3X datasets.

We also observe that fragment length distributions have a significant impact on the recovered genome fraction. Assemblies derived from the `medium` fragment length distribution (median 58 bp) generally outperform those from the `short` (median 47 bp) and `ultra-short` (median 42 bp) distributions. Interestingly, datasets with `ultra-short` fragment lengths often yield a slightly higher genome fraction than those with `short` fragment lengths. This trend is particularly pronounced for the assemblers `CarpeDeam`, `PenguiN` and `metaSPAdes`, whereas it is less evident for `MEGAHIT`.

A possible explanation for the observed differences between the `short` and `ultra-short` distributions lies in their fragment length characteristics. The `ultra-short` distribution includes a small proportion of longer fragments, with a maximum fragment length of 140 bp, while the `short` distribution is capped at 120 bp. Although these long fragments are rare, they may have a disproportionate effect on assembly performance by providing large overlaps.

Looking at the heatmaps, we observe that the highest genome fraction is achieved by both `CarpeDeam` modes under the parameter combination of `ultra-high` damage and `medium` fragment length. This is a counterintuitive observation, as one would expect better results with less damage, such as in the `high` or `moderate` damage scenarios. Interestingly, the `moderate` damage datasets exhibit the lowest genome fraction values.

A closer examination of the damage profiles in Fig. S1 reveals that the slopes of the substitution rates differ between damage levels. While `moderate` damage has the lowest substitution rate at position 1 (for both C→T and G→A transitions), it still maintains a relatively high substitution rate at position 5. This suggests that the slope and the rate at which substitution rates decrease across positions play a significant role in influencing assembly performance.

Overall, `CarpeDeam` consistently achieves the highest recovered genome fraction across the majority of datasets. `MEGAHIT` [11] also performs well, particularly demonstrating strong performance in datasets with high coverage.

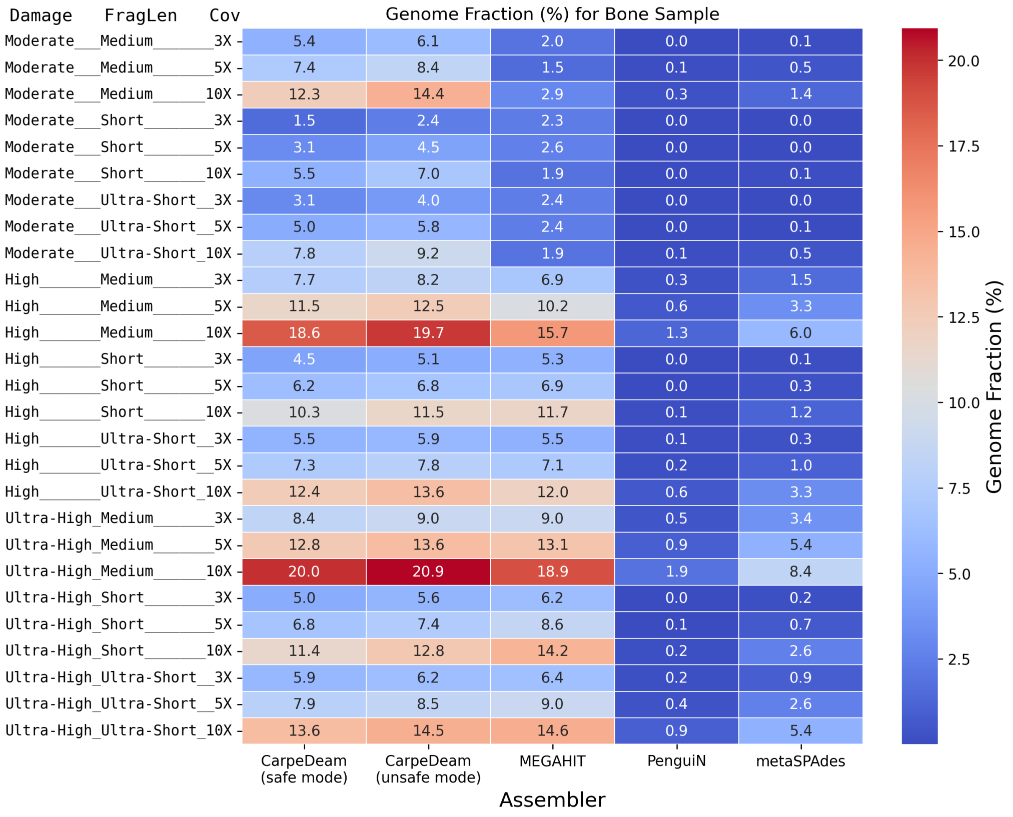

**Figure S2**: Genome fraction for the bone sample across different assemblers and parameter combinations. The plot shows the fraction of the genome recovered, as reported by metaQUAST.

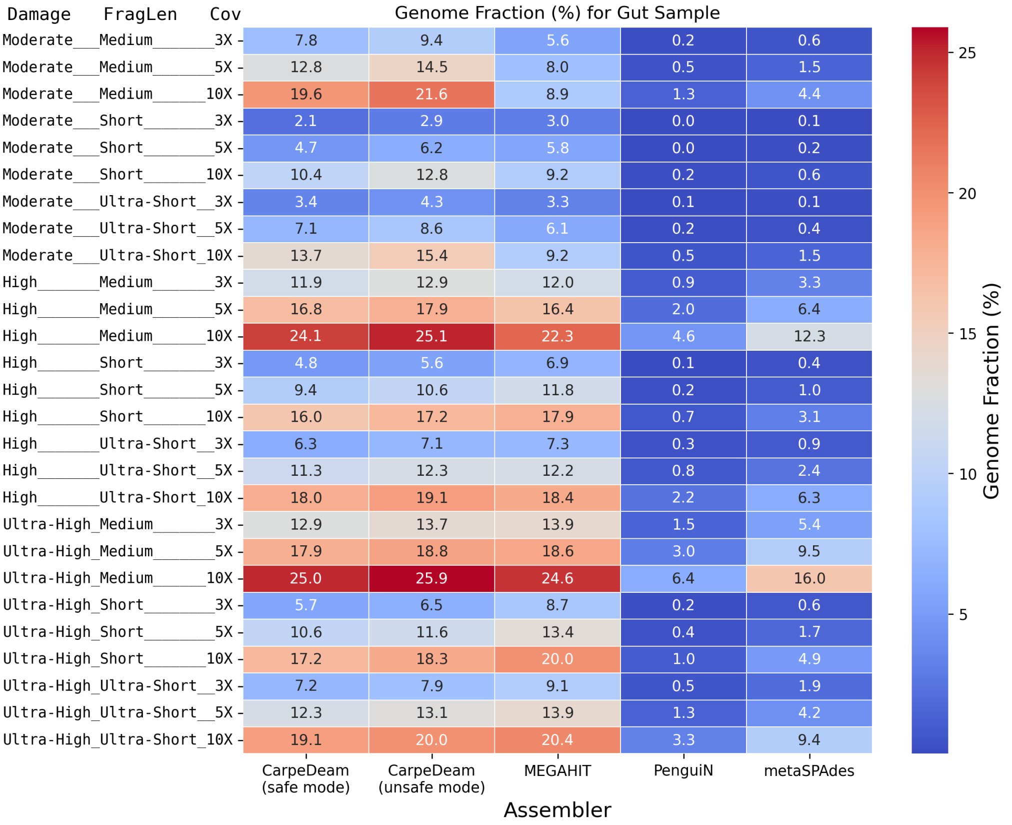

**Figure S3**: Genome fraction for the calculus sample across different assemblers and parameter combinations. The plot shows the fraction of the genome recovered, as reported by metaQUAST.

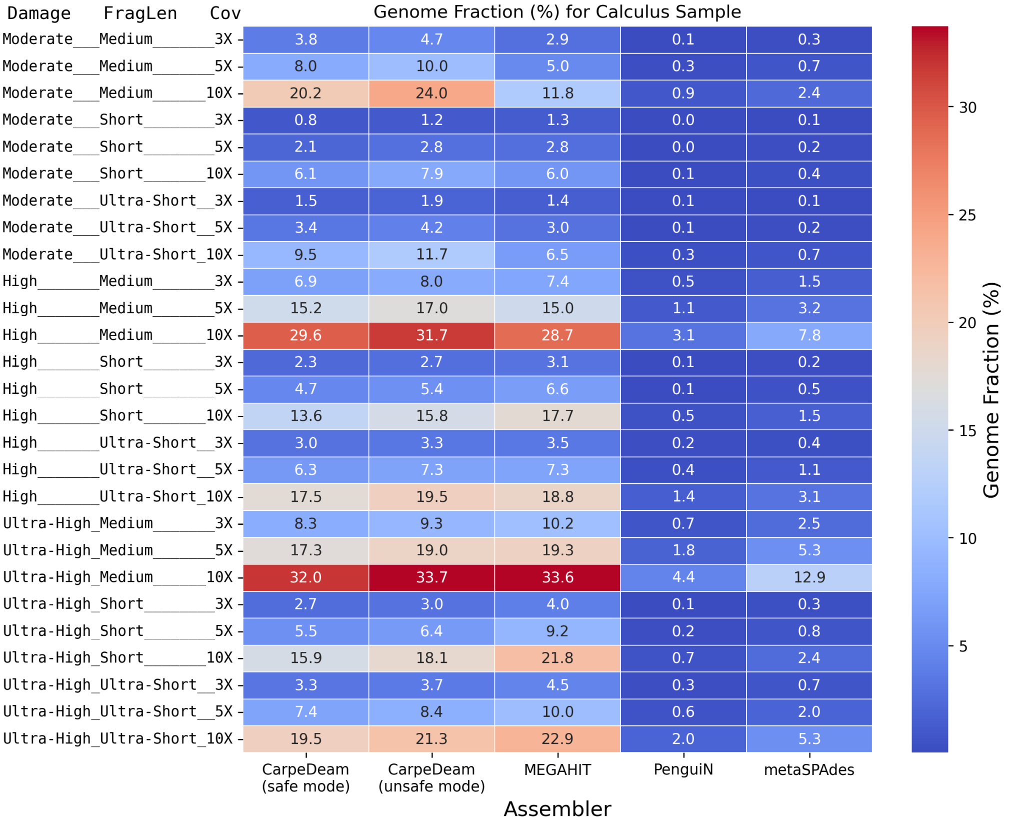

**Figure S4**: Genome fraction for the gut sample across different assemblers and parameter combinations. The plot shows the fraction of the genome recovered, as reported by metaQUAST.

# NA50

The next three heatmaps display the NA50 values for each sample and parameter combination (Fig. S5, Fig. S6, and Fig. S7). Among the assemblers, `CarpeDeam` in unsafe mode stands out, achieving notably higher NA50 values compared to the other assemblers. While `CarpeDeam` in safe mode also outperforms the other assemblers, its NA50 values are significantly lower than those observed in unsafe mode.

Interestingly, similar idiosyncrasies to those observed in the previous heatmaps showing the genome fraction metric can be observed. The NA50 values tend to be higher for datasets with challenging parameter combinations, such as those with high damage levels and shorter fragment lengths, compared to datasets with supposedly easier conditions.

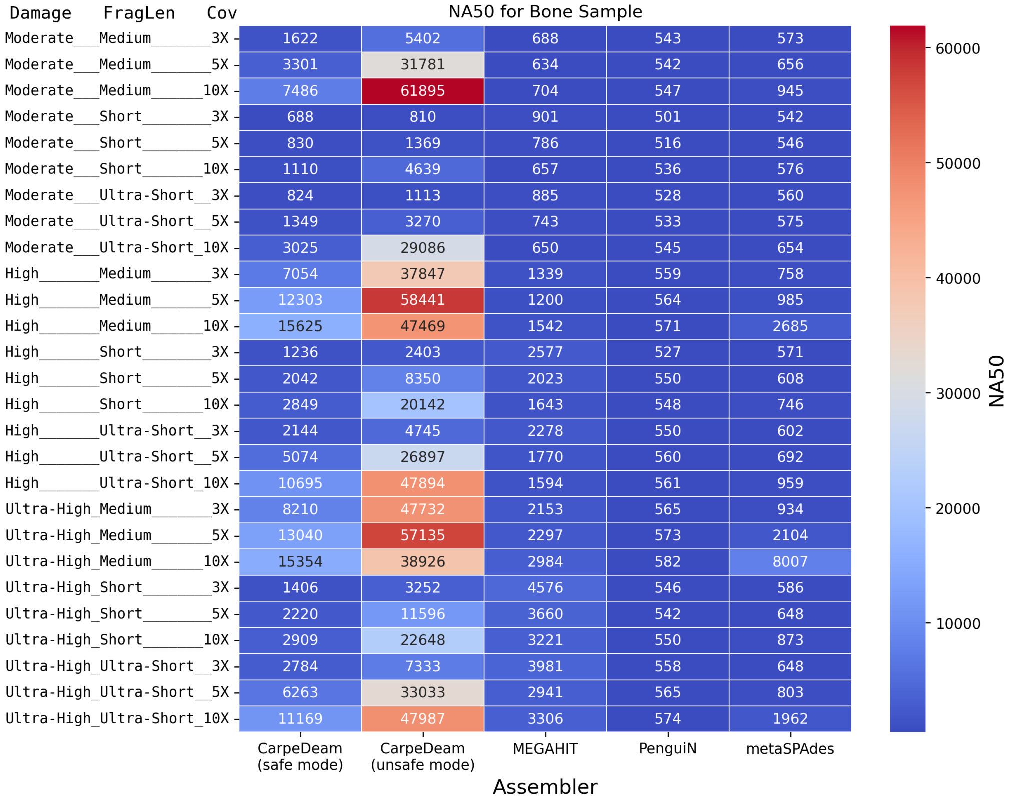

**Figure S5**: NA50 values for the bone sample across different assemblers and parameter combinations. The plot displays the NA50 values as reported by metaQUAST.

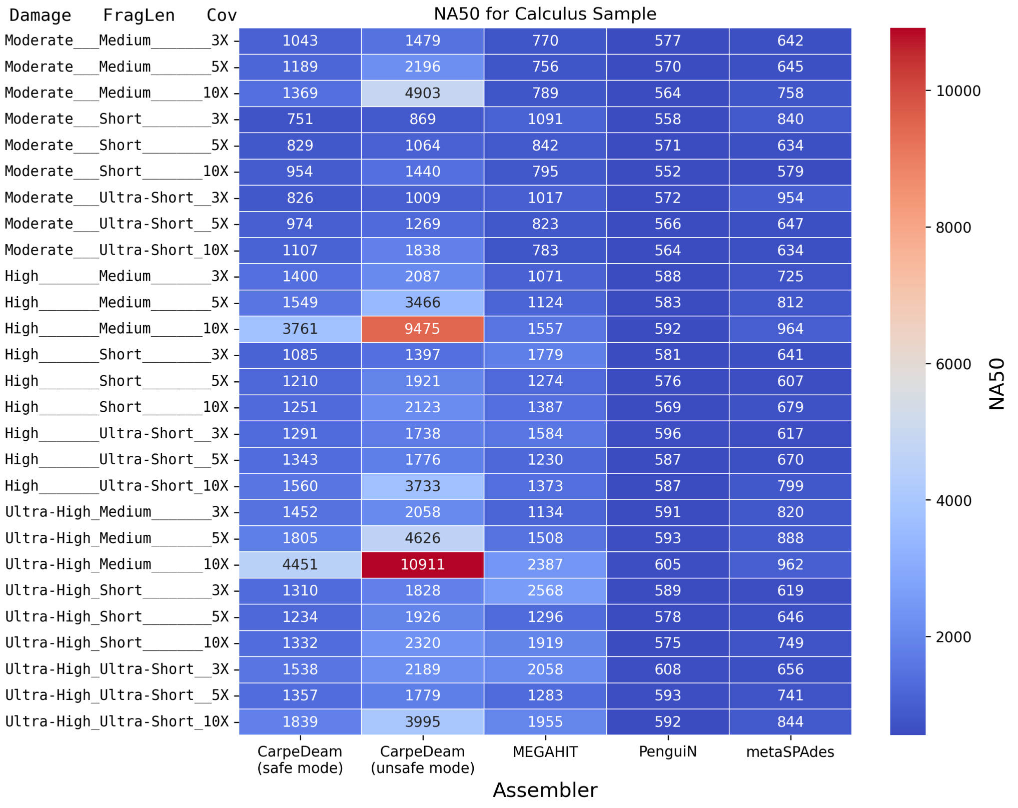

**Figure S6**: NA50 values for the calculus sample across different assemblers and parameter combinations as reported by metaQUAST.

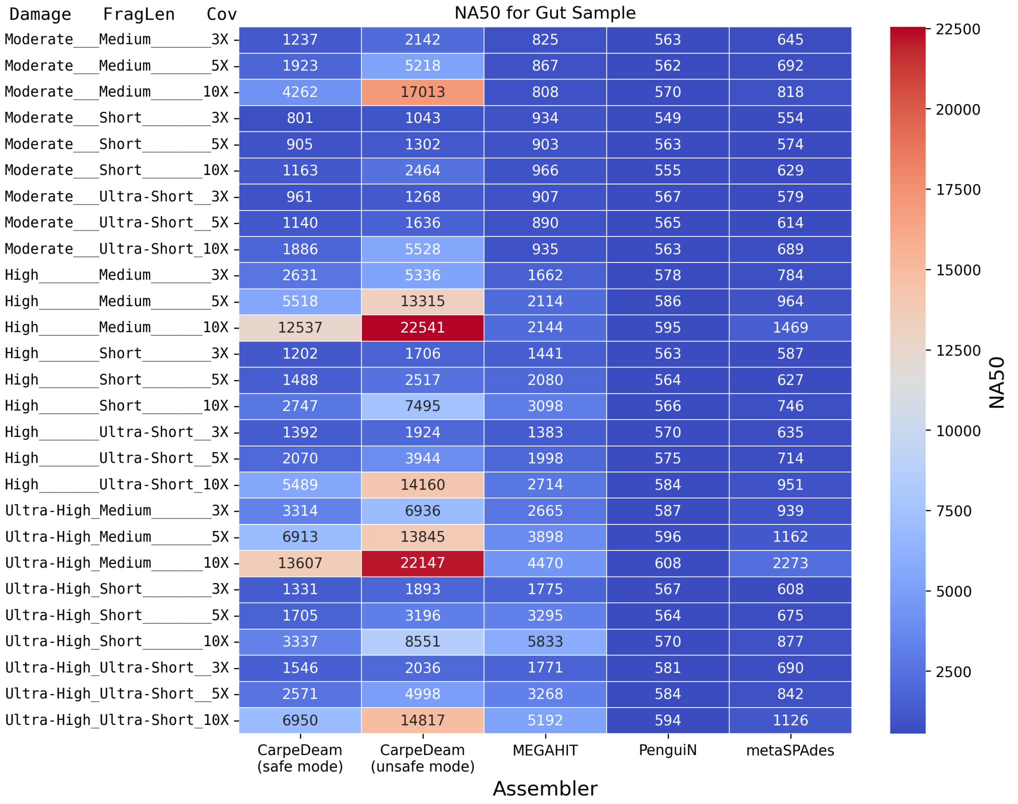

**Figure S7**: NA50 values for the gut sample across different assemblers and parameter combinations as reported by metaQUAST.

#### Largest Alignment

The next three heatmaps illustrate the largest alignment lengths for each parameter combination (Fig. S8, Fig. S9, and Fig. S10). Overall, in most cases `CarpeDeam` in unsafe mode achieved the longest alignments, followed by the safe mode and `MEGAHIT`. In many cases, `CarpeDeam` in unsafe mode converged to a specific maximum alignment length. For instance, in the bone dataset (Fig. S8), `CarpeDeam` frequently reached alignment lengths just under 200,000 bp across several parameter combinations. This observation is consistent with the fact that `CarpeDeam` currently sets a default maximum contig length of 200,000 bp, meaning the largest contigs align fully in these cases. However, this parameter can be adjusted to allow for longer contigs if required. In contrast, the safe mode of `CarpeDeam` yielded notably shorter alignments. Among the other assemblers, `MEGAHIT` consistently performed well, frequently achieving the second-longest alignment lengths. Notably, `metaSPAdes` assembled exceptionally long contigs that aligned to the reference in datasets with `medium` fragment lengths and higher coverage (5X and 10X). In three instances across all datasets, `metaSPAdes` even produced the longest aligning contigs.

Interestingly, as observed with the NA50 and genome fraction metrics, the parameter combinations that might be considered challenging, such as `ultra-high` damage and `ultra-short` fragment lengths, tended to result in longer alignments compared to the supposedly easier datasets.

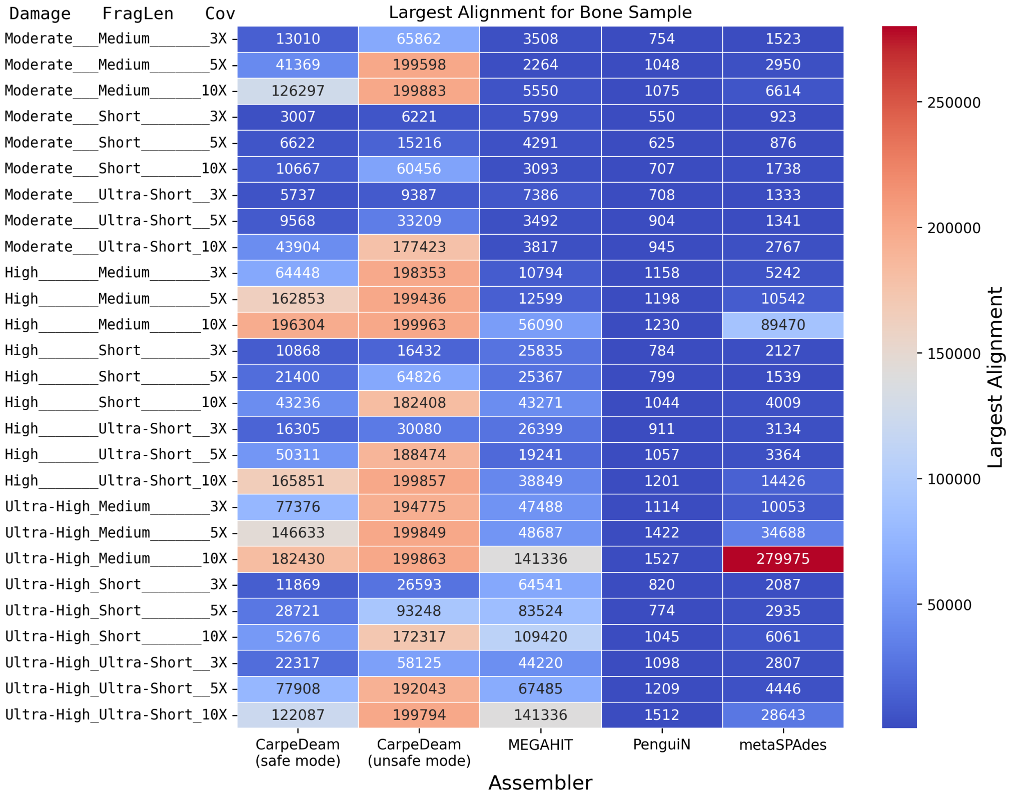

**Figure S8**: Largest alignment for the bone sample across different assemblers and parameter combinations as reported by metaQUAST.

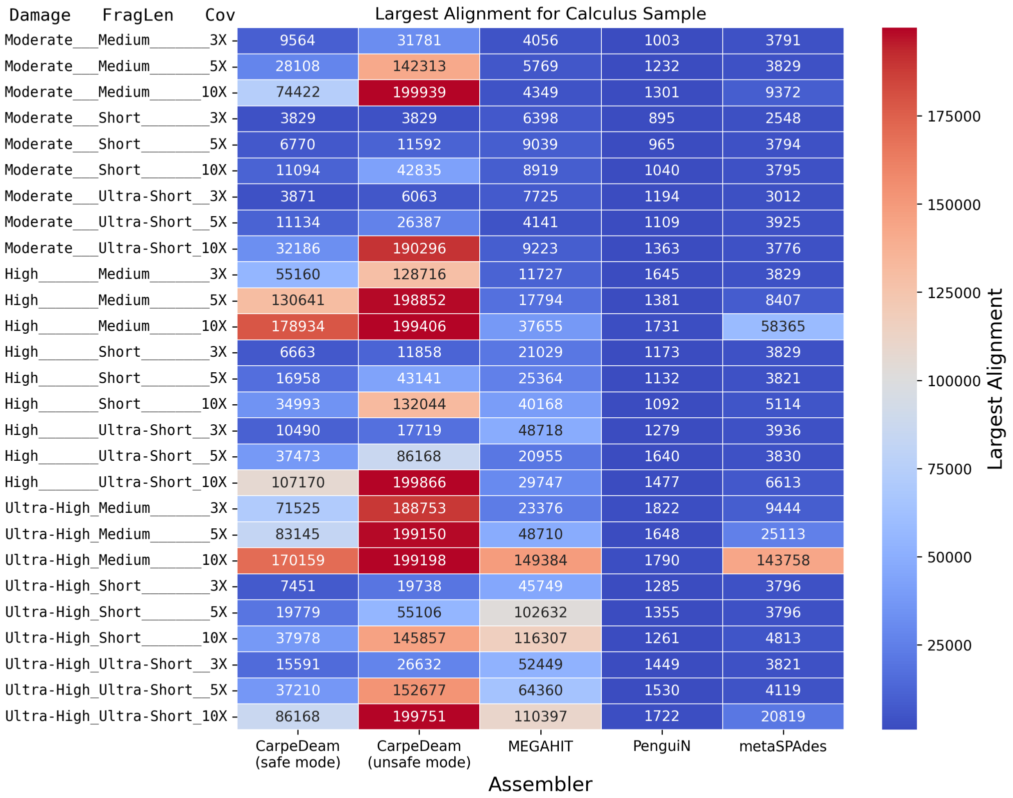

**Figure S9**: Largest alignment for the calculus sample across different assemblers and parameter combinations as reported by metaQUAST.

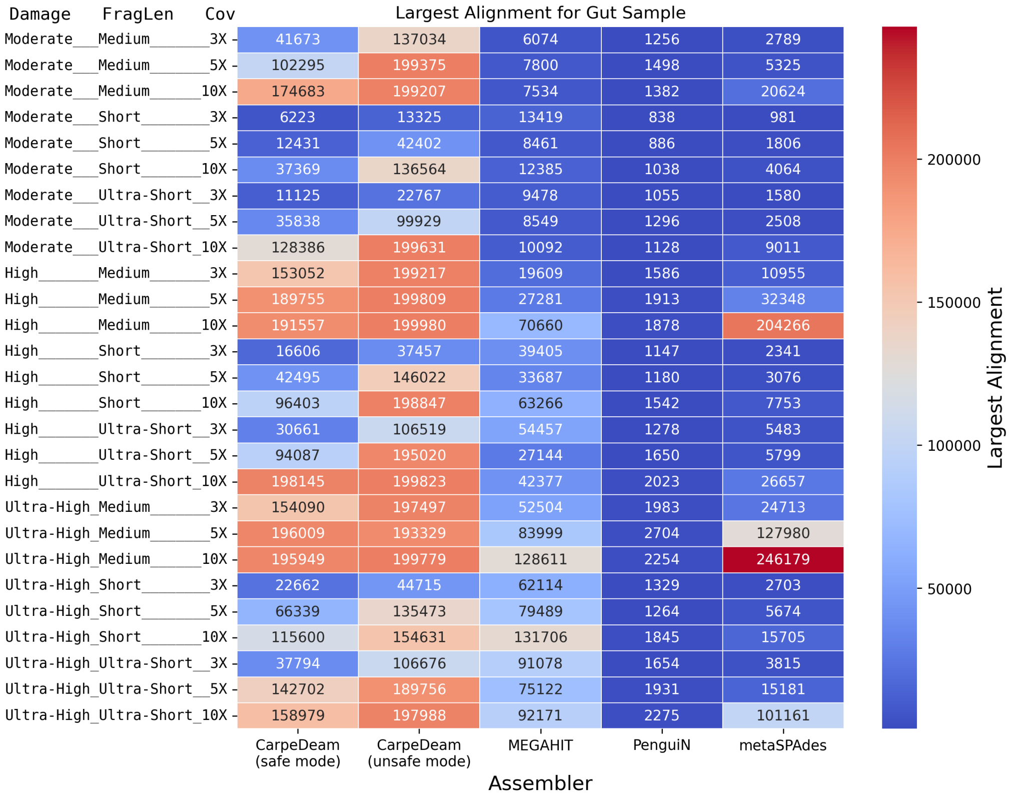

**Figure S10**: Largest alignment for the gut sample across different assemblers and parameter combinations as reported by metaQUAST.

#### Misassemblies per Contig

The heatmaps for misassemblies per contig are shown in Fig. S11, Fig. S12, and Fig. S13. Among the assemblers, `CarpeDeam` in unsafe mode performed the worst in this category. In contrast, the de Bruijn graph-based assemblers, `MEGAHIT` and `metaSPAdes`, demonstrated consistently low misassembly rates, with `MEGAHIT` remaining below 2%. `metaSPAdes` also maintained misassembly rates under 2%, except for three datasets where the rate did not exceed 5%. Similarly, `PenguiN` showed low misassembly rates, likely due to its production of fewer contigs overall.

Interestingly, `CarpeDeam` in safe mode also achieved low misassembly rates in most cases, with only a few instances exceeding 5%. However, `CarpeDeam` in unsafe mode showed particularly poor performance in high-coverage datasets, with rates ranging from 5% to 10% in many cases. Notably, extreme values were observed in the 10X coverage datasets, where the misassembly rate reached up to 38%.

Of course, these such high misassembly rates are concerning. Therefore, downstream analysis is essential to examine contigs for potential misassemblies. The reasons behind these high misassembly rates and potential approaches for conducting downstream analyses are elaborated in the discussion.

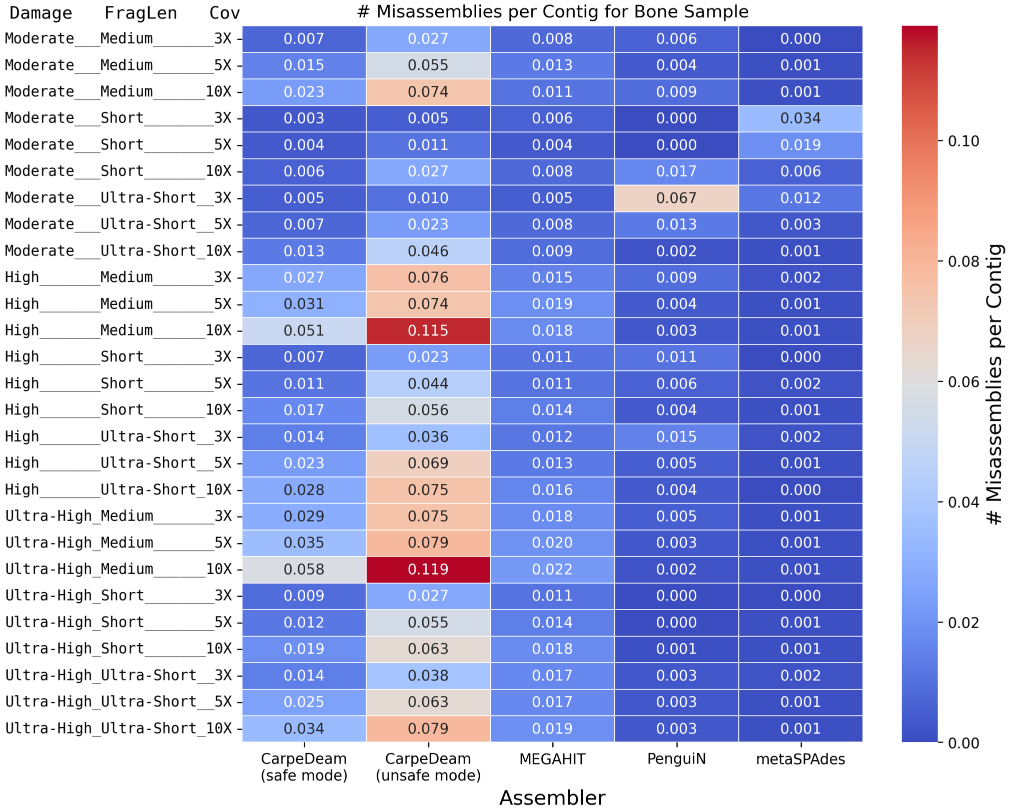

**Figure S11**: Misassemblies per contig for the bone sample across different assemblers and parameter combinations.

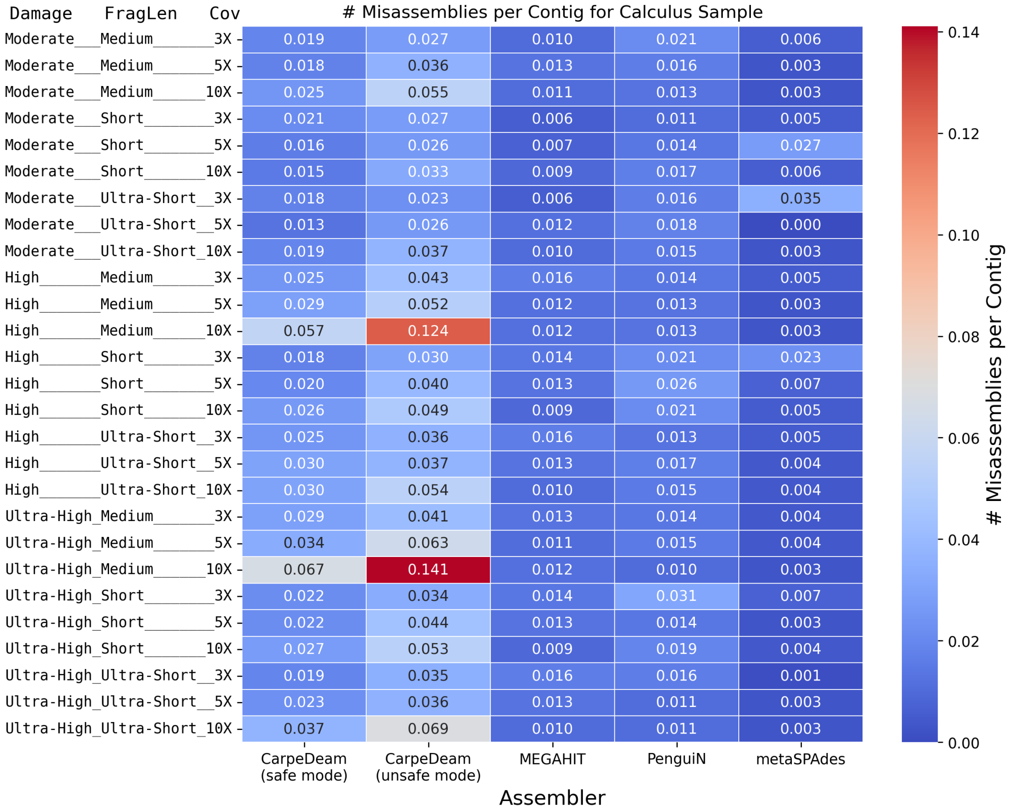

**Figure S12**: Misassemblies per contig for the calculus sample across different assemblers and parameter combinations.

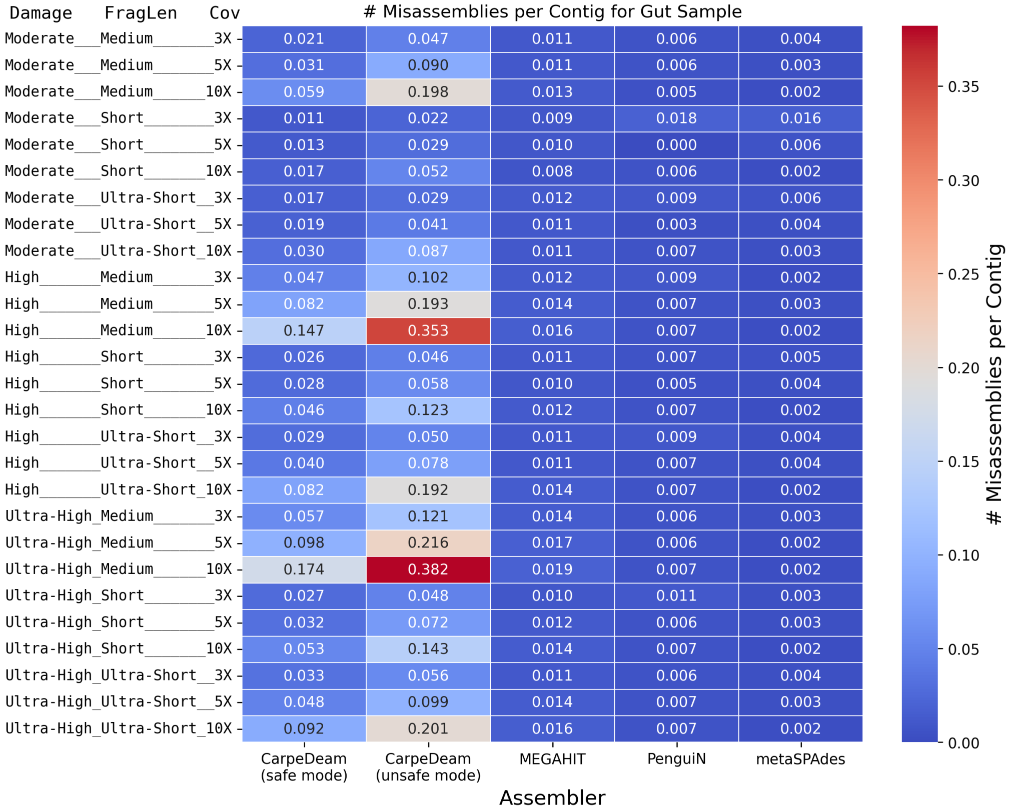

**Figure S13**: Misassemblies per contig for the gut sample across different assemblers and parameter combinations.

In Supplementary File 1, we provide the complete `metaQUAST` results for all 81 datasets as tables, omitted here for clarity and readability.

### S3 Redundancy in Assembly and its Effect on Protein Analysis

Fig. **S14** demonstrates our strategy to address potential inflation of unique protein counts due to assembly redundancy. We analysed the simulated datasets with `short` fragment length and `moderate`. We implemented a three-step approach: (1) clustering predicted proteins from contigs at 100% sequence identity and 80% coverage, followed by searching against reference proteins with the `MMseqs2 map` [14] module; (2) clustering reference proteins under identical parameters, then mapping against contig-derived proteins; and (3) employing `miniprot` to search reference proteins against nucleotide sequences. This methodology, illustrated in Fig. **S14**, enables a comprehensive assessment of protein diversity while accounting for duplications in both assembled contigs and reference genomes, thereby providing a more accurate representation of the true protein landscape across different assemblers and datasets.

### S4 Non-uniform Simulated Datasets Assembly and Investigation of Misassemblies

To further evaluate and validate the assemblies of empirical datasets, we created non-uniform datasets that mirror the difficulties of empirical data. Specifically, these datasets have the additional difficulty of having uneven fragmentation patterns and damage rates across bacterial species. We utilized aMGSIM [15] to simulate datasets that imitate the empirical samples by incorporating different damage profiles for various species, including modern non-damaged species, and simulating species-specific fragment length distributions. These parameters were estimated by mapping the reads against a reference database and analyzing the resulting alignments. Detailed information about this process can be found at `https://github.com/genomewalker/aMGSIM`.

While this approach provides a more accurate representation of a metagenomic environment, it is important to note that the empirical data may contain species and/or strains not present in any existing database. This non-uniform dataset approach allows for a controlled evaluation of assembler performance under conditions that closely resemble real ancient metagenomic samples, while providing a known ground truth for assessment.

The `metaQUAST` [13] metrics for the non-uniform simulated datasets revealed results similar to those

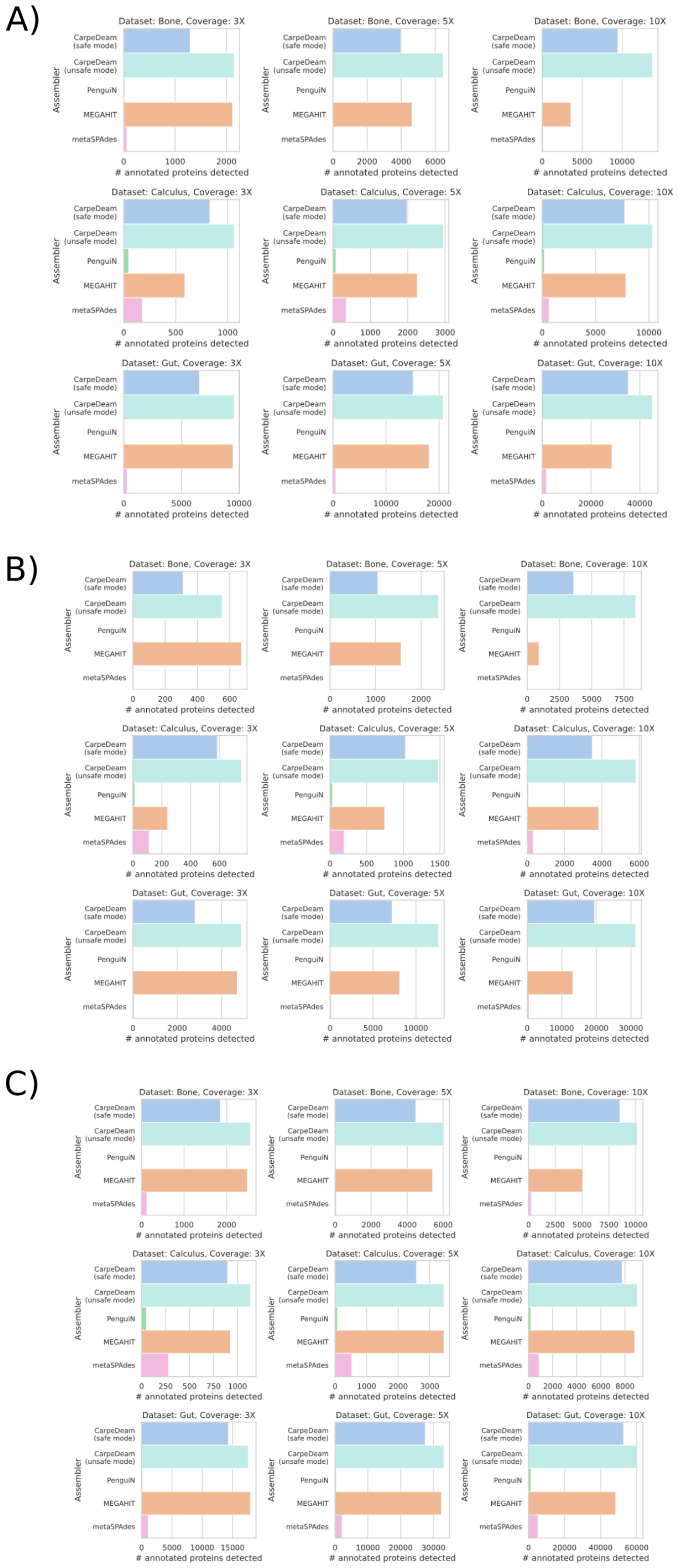

**Figure S14**: Protein redundancy analysis across different assemblers. (A) Contig-derived proteins clustered and mapped to reference proteins. (B) Reference proteins clustered and mapped to contig-derived proteins. (C) Reference proteins directly mapped to contig DNA sequences using `miniprot`. Each panel shows the number of unique proteins identified after accounting for redundancy, demonstrating the performance of different assemblers across various datasets.

observed in our main simulations (Fig. **S15**). `CarpeDeam`'s assemblies demonstrated the highest NA50 values. However, in contrast to the empirical datasets, `MEGAHIT` achieved the largest alignment in these non-uniform assemblies. The recovered genome fractions were relatively consistent across all datasets and assemblers. `metaSPAdes` was not included in this analysis as single-end reads were generated for this analysis, which are incompatible with `metaSPAdes`' requirement for paired-end data. We observed a notable increase in misassembly rates in the non-uniform GDN001 dataset assembly for `CarpeDeam` compared to other assemblers. In its unsafe mode, `CarpeDeam` produced slightly over 0.15 misassemblies per contig, a rate significantly higher than observed in other datasets. While the safe mode successfully reduced this rate to just above 0.1 misassemblies per contig, it still represents a noteworthy increase compared to other assemblers. Therefore we conducted further analysis to investigate why such a high misassembly rate arose when assembling the GDN001 dataset.

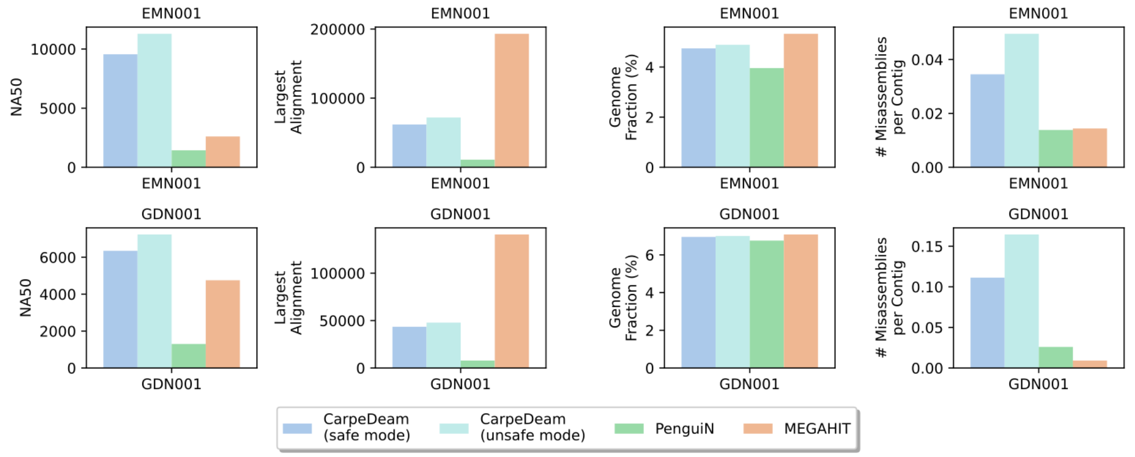

**Figure S15**: Performance evaluation of assemblers `CarpeDeam` (safe and unsafe modes), `MEGAHIT`, and `PenguiN` across non-uniform simulations EMN001 and GDN001 datasets. The metrics shown are NA50 (column 1), Largest alignment (column 2), Genome fraction (column 3), and Misassemblies per contig (column 4).

The analysis of misassembly types categorized by QUAST revealed that interspecies and intraspecies misassemblies were the predominant classes. Interspecies misassemblies result in chimeric contigs containing sequences from different species, while intraspecies misassemblies are chimeric contigs with sequences from different genomic regions within the same species.

We investigated interspecies misassemblies, by generating a heatmap depicting the number of misassemblies per pair of reference genomes in the non-uniform simulated GDN001 dataset (Fig. **S16**, panel **(A)**). While most genome pairs had zero misassemblies, a single pair stood out as the primary source of misassemblies: *Actinomyces dentalis* and *Actinomyces sp. 900323545* (Table S1). Several studies have identified *Actinomycetes* as particularly challenging to assemble due to their high GC content and repetitive regions, suggesting long-read sequencing technologies [16, 17, 18] for assembly of *Actinomycetes* species. However, the inherently ultra-short nature of ancient DNA fragments prevent the use of long-read sequencing in this context. The studies' findings coincide with what we observed when backtracking the assembly of individual contigs. Many chimeric contigs emerged, when identical sequences of considerable length (over 50bp) were present in reads from two different species. In such cases `CarpeDeam` could not distinguish between the two species and was left with two perfect extension candidates.

To validate both the reasons proposed in the literature and our own findings, we employed a $k$-mer approach to quantify exact sequence matches between species. We compared the set of the three most abundant species with the three most misassembly-prone species, resulting in a set of five species: *Actinomyces_dentalis*, *Actinomyces_sp900323545*, *Desulfobulbus_oralis*, *Flexilinea_sp001717545*, and *Pauljensenia_cardiffensis*. Fig. **S16**, panel **(B)** clearly demonstrates a significantly higher rate of shared $k$-mers (sizes 50 to 100) between *Actinomyces dentalis* and *Actinomyces sp. 900323545*. We retrieved all $k$-mers occurring at least once from each reference genome using `CHTKC` [19], reporting the fraction of shared $k$-mers among the set of unique $k$-mers.

Examination of intraspecies misassemblies revealed that *Actinomyces dentalis* was also the primary source of chimeras from different locations within a genome (Fig. **S17**, panel **(A)**). To evaluate whether this was due to highly similar sequences within the genome, we utilized the `easy-search` module of `MMseqs2` [14] in `--search-mode-3` and high sensitivity `-s 7.5`, searching the reference genomes against themselves. We reported all alignments with ≥99% sequence identity at different genomic locations and plotted the density of occurrences per alignment length (Fig. **S17**, panel **(B)**). While three out of five species indicated a uniform distribution of conserved sequences from different regions, two outliers emerged: *Desulfobulbus oralis* showed a significantly high amount of ultra-conserved sequences up to 250bp within its genome, and *Actinomyces dentalis*, the most abundant species in the non-uniform simulated GDN001 dataset and the reference with the most intraspecies misassemblies, showed a significant large fraction of ultra-conserved sequences with at least 99% sequence identity up to 500bp.

Our analysis identified specific scenarios where `CarpeDeam`'s approach faces limitations in assembling exclusively correct contigs. The non-uniform simulated GDN001 dataset exemplifies a combination of factors that can contribute to an increased misassembly rate: Firstly, *Actinomyces dentalis* and *Actinomyces sp. 900323545* are the highest abundant species in the simulated GDN001 dataset (Table S2) and share a considerable number of $k$-mers. This abundance increases the likelihood of encountering situations with two perfect overlaps during extensions, where `CarpeDeam` struggles to determine the optimal choice. While the safe mode demonstrates improvement in this regard, it does not entirely resolve the issue. Secondly, the *Actinomyces dentalis* genome contains a comparatively large fraction of highly conserved sequences (Fig. **S17**, panel **(B)**), further complicating the assembly process due to the aforementioned reasons. It is worth noting that Mobile Genetic Elements (MGEs) are known to present a challenge in assembly, and tools for their identification have been published [20]. Sheinman *et al.* [21] report that the frequency of horizontal gene transfer events varies among different bacterial taxa. They have carefully modeled the frequency of 100% identical sequences found in bacteria that are likely due to horizontal gene transfer events.

Despite these challenges, the simulated EMN001 dataset and the main simulations demonstrate that such problems are not consistently encountered with `CarpeDeam`. However, it is crucial to acknowledge this limitation of `CarpeDeam`'s approach. The next step in advancing ancient metagenome assembly could involve combining De Bruijn graphs with our overlap-centric approach, potentially mitigating these challenges and improving the overall assembly quality.

| Species1 | Species2 | # Misassemblies |
|---|---|---|
| *Actinomyces dentalis* `GCF_000429225.1` | *Actinomyces sp.* 900323545 `3300008059_1` | 521 |
| *Actinomyces dentalis* `GCF_000429225.1` | *Pauljensenia cardiffensis* `GCA_905373095.1` | 39 |
| *Actinomyces sp.* 900323545 `3300008059_1` | *Pauljensenia cardiffensis* `GCA_905373095.1` | 25 |
| *Actinomyces dentalis* `GCF_000429225.1` | *Brooklawnia sp.* `GCF_000413315.1` | 12 |
| *Flexilinea sp.* `GCA_001717545.1` | *Hornefia minuta* `GCA_003433295.1` | 5 |
| *Desulfobulbus oralis* `GCA_905373745.1` | *Desulfomicrobium orale* `GCF_001553625.1` | 4 |
| *Actinomyces sp.* 900323545 `3300008059_1` | *Lautropia mirabilis* `3300009381_5` | 4 |
| *Actinomyces dentalis* `GCF_000429225.1` | *Hornefia minuta* `GCA_003433295.1` | 3 |
| *Actinomyces dentalis* `GCF_000429225.1` | *Arachnia sp.* `3300008089_8` | 2 |
| *Actinomyces dentalis* `GCF_000429225.1` | *Flexilinea sp.* `GCA_001717545.1` | 2 |

Table S1: Interspecies Miassemblies: Assembly of the non-uniform simulated GDN001 dataset with `CarpeDeam` (unsafe mode).

| Species | Coverage |
|---|---|
| *Actinomyces dentalis* `GCF_000429225.1` | 11.4 |
| *Actinomyces sp.* 900323545 `3300008059_1` | 4.6 |
| *Desulfobulbus oralis* `GCA_905373745.1` | 2.9 |
| *Hornefia minuta* `GCA_003433295.1` | 1.5 |
| *Flexilinea sp.* `GCA_001717545.1` | 2.4 |

Table S2: Coverage of five most abundant species in the non-uniform simulated GDN001 dataset.

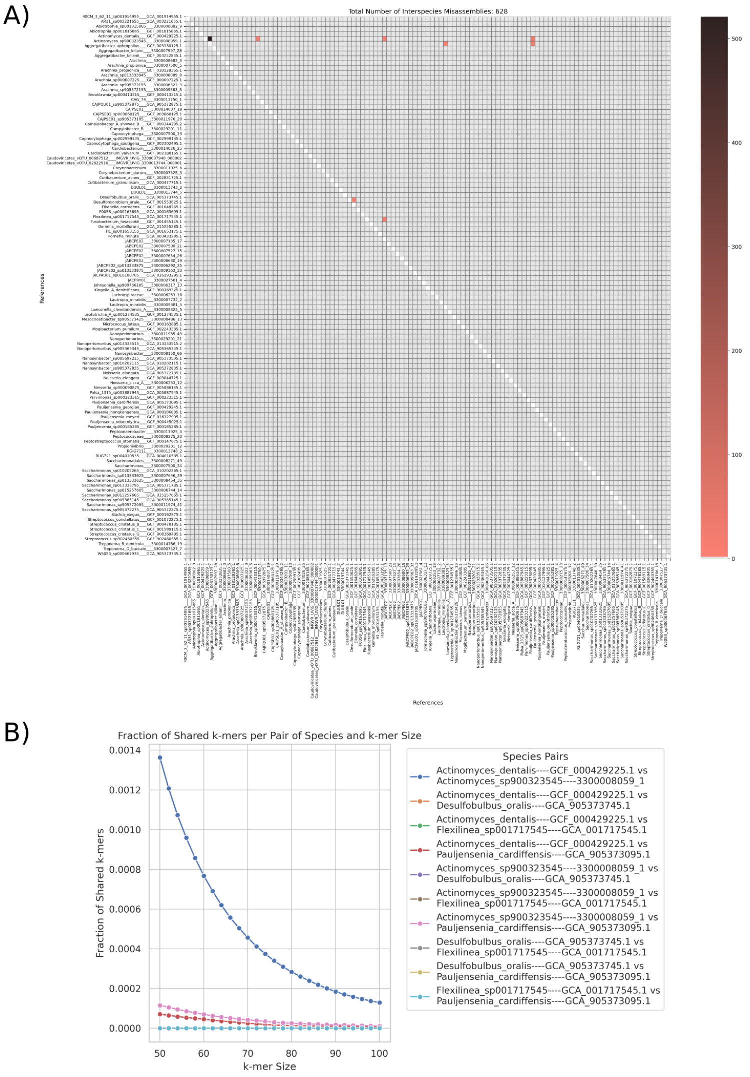

**Figure S16**: Analysis of interspecies misassemblies in the non-uniform simulated GDN001 dataset assembled by `CarpeDeam` (unsafe mode). **(A)** Heatmap showing the number of misassemblies between pairs of reference genomes. Each cell represents a pair of species, with the color intensity indicating the number of misassemblies between them. **(B)** Comparison of shared $k$-mers between selected species. The x-axis shows $k$-mer sizes ranging from 50 to 100 bp, while the y-axis represents the fraction of shared $k$-mers among the set of unique $k$-mers. Each line corresponds to a pair of species from the set of five selected species: *Actinomyces dentalis*, *Actinomyces sp900323545*, *Desulfobulbus oralis*, *Flexilinea sp001717545*, and *Pauljensenia cardiffensis*. The species combinations that are shown in the legend, but cannot be found in the plot itself, indicate that no $k$-mers of any size are shared between those pairs. The analysis was performed using `CHTKC` [19] to retrieve and compare $k$-mers occurring at least once in each reference genome.

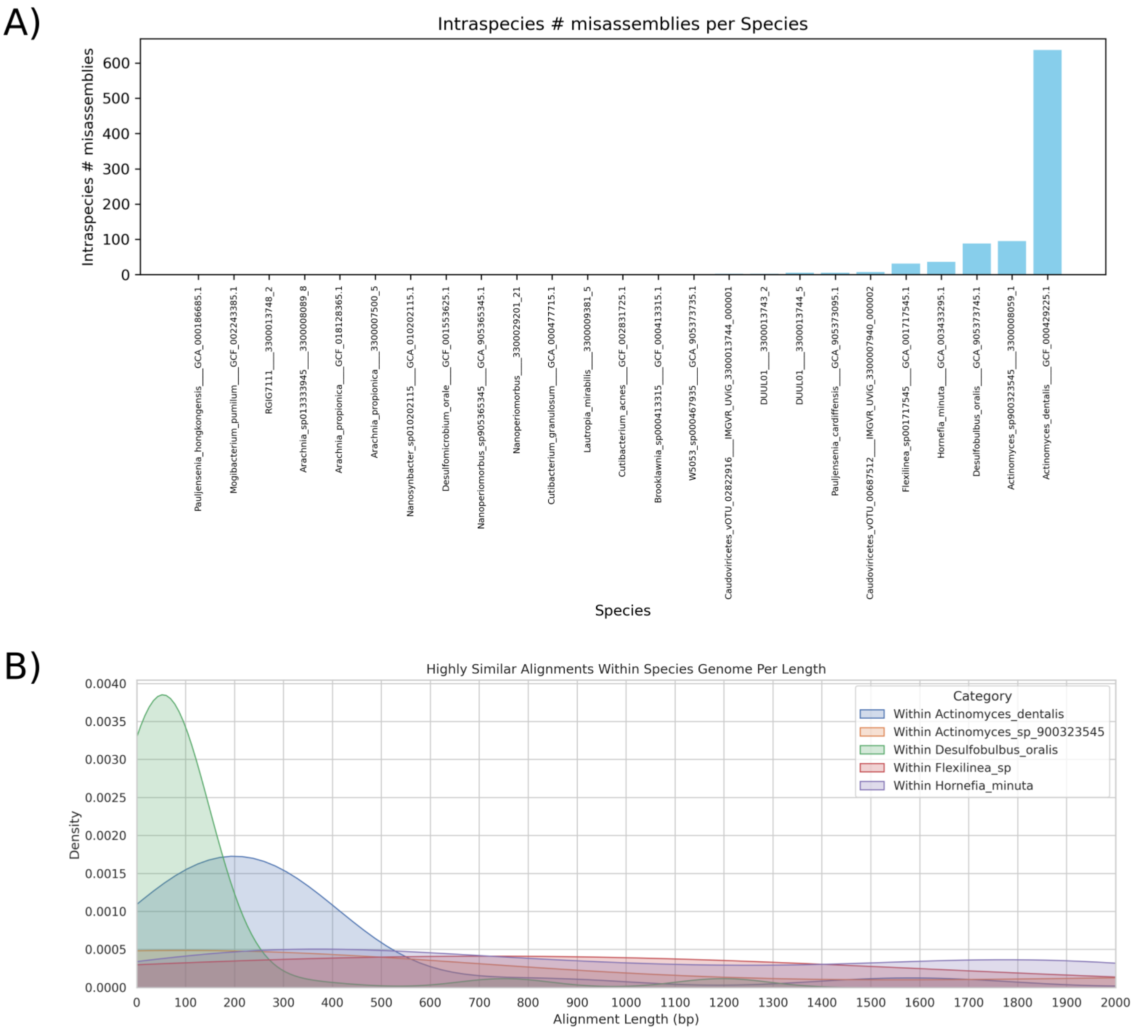

**Figure S17**: Analysis of intraspecies misassemblies in the non-uniform simulated GDN001 dataset assembled by `CarpeDeam` (unsafe mode). **(A)** Bar plot showing the number of intraspecies misassemblies for each reference genome. The x-axis lists the species names, while the y-axis represents the count of misassemblies. **(B)** Density plot of highly conserved sequences within genomes. The x-axis shows the alignment length in base pairs, while the y-axis represents the density of occurrences. Each line corresponds to a different species from the set of five selected species. The analysis was performed using the `easy-search` module of `MMseqs2` [14] in `--search-mode-3`, reporting alignments with ≥99% sequence identity at different genomic locations within each genome.

### S5 Number of Non-Misassembled Contigs per Assembler

To further support our evaluation of assembly quality, we examined the number of long, non-misassembled contigs produced by each assembler. Since genome fraction alone, as reported by `metaQUAST`, [13], does not capture contiguity, we focused on contigs exceeding 2000 bp across datasets with short fragment length and moderate damage. The results, presented in Fig. S18, highlight differences in assembler performance, particularly showcasing `CarpeDeam`'s ability to reconstruct longer, accurate contigs compared to other methods.

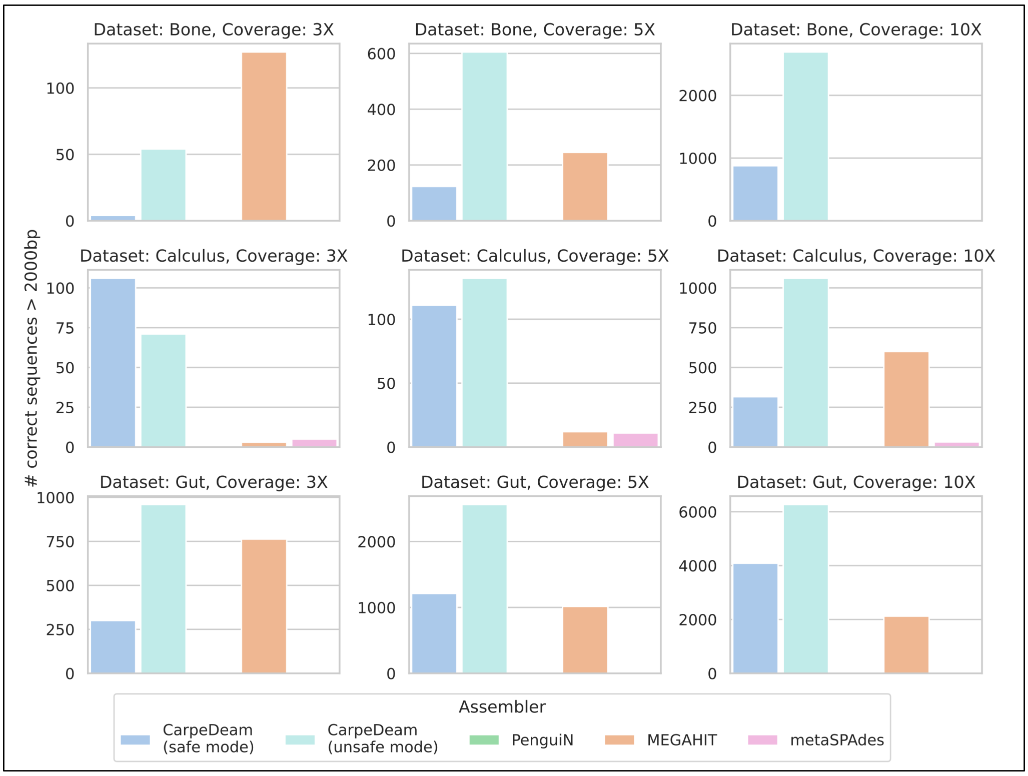

**Figure S18**: Comparison of non-misassembled contigs over 2000 bp produced by different assemblers across simulated metagenomic datasets. The bar plots show the number of non-misassembled contigs exceeding 2000 bp for `CarpeDeam` (safe and unsafe modes), `MEGAHIT`, `metaSPAdes`, and `PenguiN`. Results are presented for datasets with `moderate` damage and `short` fragment length distribution across three simulated environments (bone, dental calculus, and gut) with varying coverage levels (3X, 5X, and 10X). Each row represents an environment (from top to bottom: bone, calculus, gut), while columns represent coverage levels (from left to right: 3X, 5X, 10X). Within each subplot, bars indicate the performance of individual assemblers for the specified environment and coverage combination.

### S6 Damage Patterns in Empirical Datasets

Fig. S19 illustrates the C→T substitution frequencies across 20 empirical datasets from five distinct sample groups (TAF, EMN, ECO, OAK, and GDN). Each dot represents the substitution frequency of a single species, with overlapping dots resulting in different colour shades; for improved readability, only the C→T substitution frequencies are shown. The black line indicates the average damage frequency per position, which is used as input for `CarpeDeam` assemblies. Additionally, the figure highlights variability in damage patterns—showing more homogeneous profiles in the GDN and EMN datasets and increased variability in the ECO group. Accession IDs and further details about the empirical datasets are provided in the data repository (see Section 7, Code Availability).

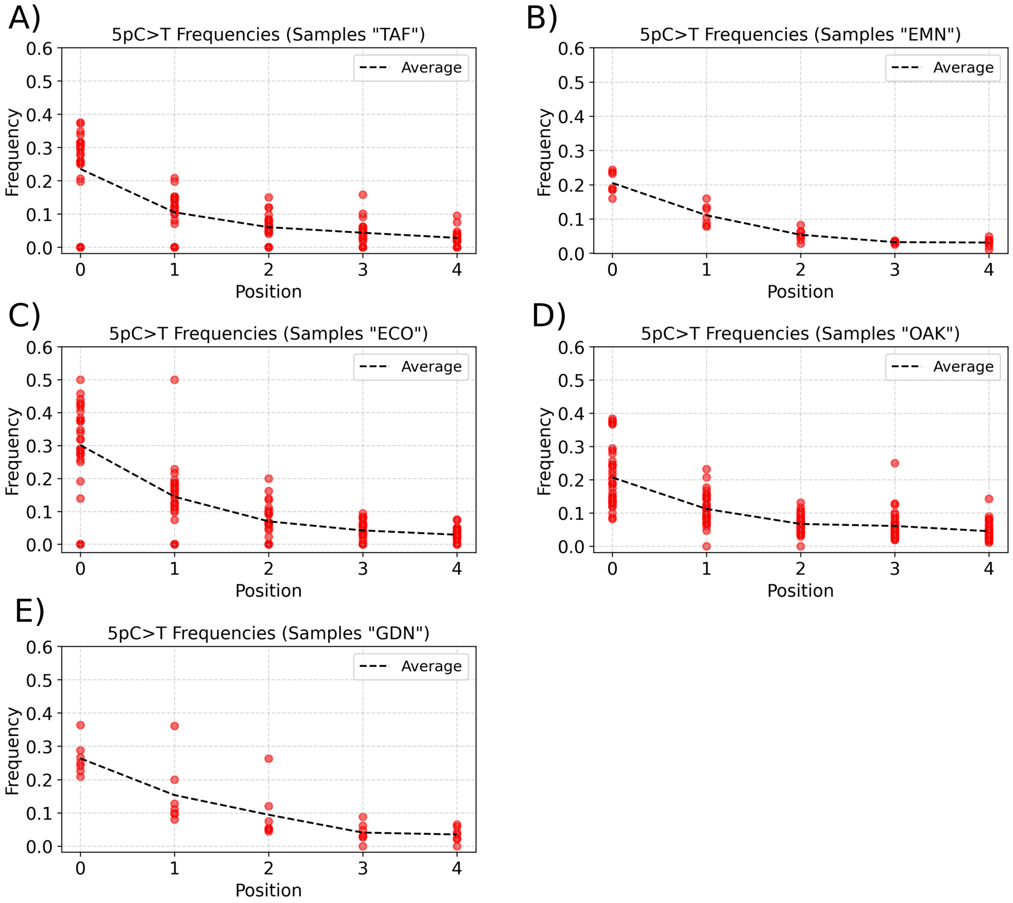

**Figure S19**: Overview of damage patterns in 20 empirical datasets, grouped by their sample sites **(A)** TAF, **(B)** EMN, **(C)** ECO, **(D)** OAK, and **(E)** GDN. The damage patterns belong to the species *Fusobacterium nucleatum*, *Porphyromonas gingivalis*, *Pseudopropionibacterium propionicum*, *Streptococcus gordonii*, *Tannerella forsythia*, *Treponema denticola*, and *Treponema socranskii* as reported in the data repository of the study by Fellows-Yates *et al.* [1]. Each dot represents the C→T substitution frequency of a single species. The black line indicates the average substitution rate at each position. There are more than seven values per position, since each sample group consists of several samples. The damage matrix containing the average values was used as input for `CarpeDeam` assemblies of these datasets. We observe that the damage patterns vary across datasets. While the GDN and EMN datasets show more homogeneous damage spread, the ECO dataset exhibits the highest variability.

### S7 Algorithm Details

#### S7.1 Likelihood model for Read Extension

The likelihood model that is used stop the extension during the second phase, where only unmodified aDNA fragments are considered and the damage patterns remain valid, can be illustrated as in Fig. S20.

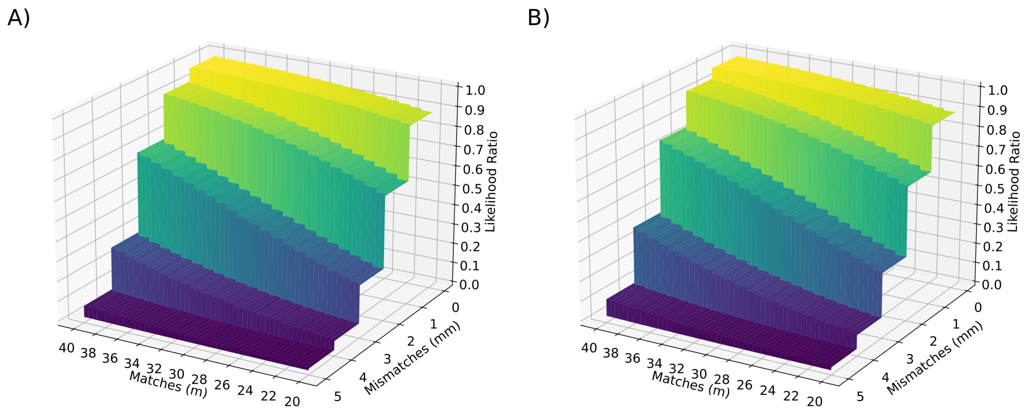

**Figure S20**: Illustration of `CarpeDeam`'s likelihood model to determine the end of the extension during the initial extension phase of the algorithm where reads are selected to extend the center sequence of the cluster. **(A)**) The likelihood ratio (Z axis) with mild damage with a certain number of matches (X axis) and mismatches (Y axis). This likelihood ratio determines if the extension is stopped or not. If the ratio falls below 0.5, the extension is stopped. For instance, with 2 mismatches and 20 matches, the algorithm would conclude that these are not convincingly more likely than a random match given that the read has some similarity to the center sequence. This similarity is by design as these were selected and aligned to the center sequence. More matches and fewer mismatches trigger a continuation of the extension process. **(B)**) same figure but with higher damage. This subfigure shows that the algorithm is more lenient to mismatches as damage rates are higher.

#### S7.2 RYmer Space and Sequence Identity

RYmer space offers a simplified representation of sequences by translating them into a binary system of purines (R) and pyrimidines (Y). This binary encoding effectively reduces sequences to a two-character alphabet, where purines ($A$ and $G$) are represented as one binary state, and pyrimidines ($C$ and $T$) as another.

We applied this concept in the clustering process prior to assembly. A cluster contains sequences that share at least one $k$-mer. Members are filtered by sequence identity of the overlap between center sequence and member sequence. This filtering is crucial to avoid unrelated sequences being taken into account for extension. The sequence identity in native `PenguiN` is as high as 99%, reflective for the low error rates in modern metagenomic data. Ancient DNA fragments on the other hand are characterized by shorter lengths and deaminated bases. Therefore, a clustering threshold set at 90% sequence identity can incorporate both deaminated, yet authentic sequences and numerous unrelated sequences. To mitigate this, we convert overlapping sequences to RYmer space and applying a stringent threshold for RYmer sequence identity. This strategy allows for the inclusion of mismatches due to deamination, thus improving the assembly of ancient DNA sequences.

The concept of RYmer space translates sequences into a format where the effects of deamination, do not impede the identification of genuinely related sequences. It should be noted that while RYmer sequence identity can help refine clusters after they have been clustered based on shared $k$-mers, RYmers are not particularly suitable for use in the clustering process itself. Our investigations revealed limitations in specificity for clustering. RYmers, due to their reduced information content, are less specific as $k$-mers. As ancient DNA fragments are inherently short, doubling the size of RYmers to achieve a similar specificity as $k$-mers is not viable in our application (see Fig. **S21**). This diminished specificity led to the conclusion that while RYmers enhance the assembly process by allowing for deamination-induced mismatches, they do not offer sufficient specificity for effective clustering, compared to $k$-mers.

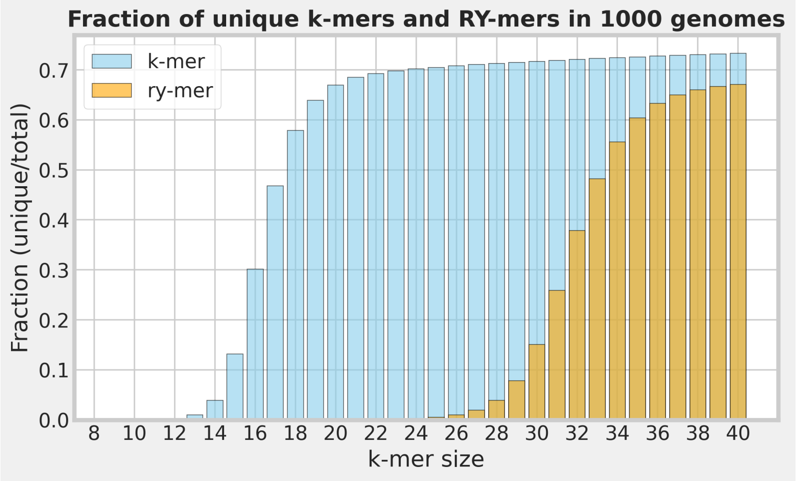

**Figure S21**: Fraction of unique $k$-mers and RYmers in a sample of 1000 genomes. In native PenguiN a default $k$-length of 22 is used in the clustering process. The same level of specificity is not achieved by RYmers smaller than 40bp. As ancient DNA fragments can be ultra short (e.g. 35bp), RYmers of length greater 30bp are not applicable for clustering.

#### S7.3 Implementation of the Safe Mode

`CarpeDeam` clusters sequences using shared k-mers, enabling whole-sequence overlap detection that preserves aDNA fragment context. To account for deamination-induced mismatches in aDNA, it applies a relaxed 90% sequence identity threshold along with an RYmer identity filter that tolerates purine–pyrimidine mismatches. Due to the short length and reduced specificity of aDNA fragments—particularly during early extension phases where deamination can lead to misassemblies we implemented a "safe mode". In this mode, the left and right extending regions of the center sequence are independently evaluated, and consensus sequences (one per end) are generated from overlapping extension candidates (requiring a minimum coverage of five sequences per position). The approach is illustrated in Fig. **S22**. This approach minimizes chimeric assemblies by ensuring that only well-supported extension regions are integrated.

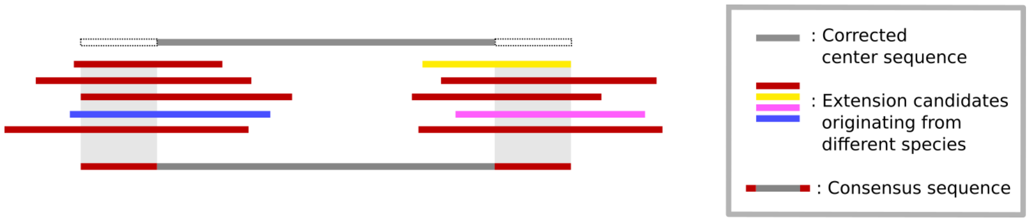

**Figure S22**: Consensus calling in the extending regions in a sequence cluster. `CarpeDeam` derives a consensus sequence from regions covered by at least five member sequences. Users can disable this "safe mode" to increase sensitivity, potentially at the cost of higher chimeric content ("unsafe mode"). The top sequence in grey is the center sequence: The longest sequence in the cluster that is to be extended.

### S8 Memory and Runtime

We tested all assemblers on a single node with a 32 core AMD EPYC-Rome 1996.250 MHz processor with 472GB available memory. The reported number refer to the assembly of the datasets with `moderate` damage and `short` fragment length distribution (see Section S1 in the Supplmentary Material).

Table S3: Gut

| assembler | sample | damage | coverage | runtime (h:m:s) | RAM (Gb) |
|---|---|---|---|---|---|
| `CarpeDeam` (safe mode) | Gut | High | 3X | 0:49:00 | 15.64 |
| `CarpeDeam` (unsafe mode) | Gut | High | 3X | 0:53:07 | 15.93 |
| `PenguiN` | Gut | High | 3X | 0:26:39 | 13.87 |
| `MEGAHIT` | Gut | High | 3X | 0:08:43 | 1.22 |
| `metaSPAdes` | Gut | High | 3X | 0:42:14 | 18.99 |
| `CarpeDeam` (safe mode) | Gut | High | 5X | 1:35:26 | 24.54 |
| `CarpeDeam` (unsafe mode) | Gut | High | 5X | 1:27:23 | 26.71 |
| `PenguiN` | Gut | High | 5X | 0:40:24 | 23.66 |
| `MEGAHIT` | Gut | High | 5X | 0:16:53 | 2.25 |
| `metaSPAdes` | Gut | High | 5X | 1:14:38 | 18.78 |
| `CarpeDeam` (safe mode) | Gut | High | 10X | 3:19:31 | 51.27 |
| `CarpeDeam` (unsafe mode) | Gut | High | 10X | 3:12:03 | 67.47 |
| `PenguiN` | Gut | High | 10X | 1:26:52 | 49.03 |
| `MEGAHIT` | Gut | High | 10X | 0:42:07 | 4.11 |
| `metaSPAdes` | Gut | High | 10X | 3:02:43 | 25.97 |

Table S4: Dental Calculus

| assembler | sample | damage | coverage | runtime (h:m:s) | RAM (Gb) |
|---|---|---|---|---|---|
| `CarpeDeam` (safe mode) | Calculus | High | 3X | 0:16:57 | 5.01 |
| `CarpeDeam` (unsafe mode) | Calculus | High | 3X | 0:17:13 | 5.38 |
| `PenguiN` | Calculus | High | 3X | 0:07:40 | 4.98 |
| `MEGAHIT` | Calculus | High | 3X | 0:02:16 | 0.46 |
| `metaSPAdes` | Calculus | High | 3X | 0:13:22 | 2.77 |
| `CarpeDeam` (safe mode) | Calculus | High | 5X | 0:28:37 | 8.49 |
| `CarpeDeam` (unsafe mode) | Calculus | High | 5X | 0:26:33 | 10.62 |
| `PenguiN` | Calculus | High | 5X | 0:13:33 | 7.08 |
| `MEGAHIT` | Calculus | High | 5X | 0:04:04 | 0.67 |
| `metaSPAdes` | Calculus | High | 5X | 0:21:58 | 9.17 |
| `CarpeDeam` (safe mode) | Calculus | High | 10X | 0:58:07 | 16.74 |
| `CarpeDeam` (unsafe mode) | Calculus | High | 10X | 0:59:38 | 19.10 |
| `PenguiN` | Calculus | High | 10X | 0:26:23 | 15.89 |
| `MEGAHIT` | Calculus | High | 10X | 0:10:05 | 1.30 |
| `metaSPAdes` | Calculus | High | 10X | 0:47:00 | 18.79 |

Table S5: Bone

| assembler | sample | damage | coverage | runtime (h:m:s) | RAM (Gb) |
|---|---|---|---|---|---|
| `CarpeDeam` (safe mode) | Bone | High | 3X | 0:27:20 | 8.00 |
| `CarpeDeam` (unsafe mode) | Bone | High | 3X | 0:26:46 | 8.44 |
| `PenguiN` | Bone | High | 3X | 0:13:18 | 7.55 |
| `MEGAHIT` | Bone | High | 3X | 0:03:06 | 0.72 |
| `metaSPAdes` | Bone | High | 3X | 0:22:53 | 6.13 |
| `CarpeDeam` (safe mode) | Bone | High | 5X | 0:45:45 | 13.44 |
| `CarpeDeam` (unsafe mode) | Bone | High | 5X | 0:48:25 | 13.88 |
| `PenguiN` | Bone | High | 5X | 0:21:15 | 12.53 |
| `MEGAHIT` | Bone | High | 5X | 0:06:50 | 1.17 |
| `metaSPAdes` | Bone | High | 5X | 0:39:30 | 18.90 |
| `CarpeDeam` (safe mode) | Bone | High | 10X | 1:38:27 | 26.90 |
| `CarpeDeam` (unsafe mode) | Bone | High | 10X | 1:34:37 | 27.17 |
| `PenguiN` | Bone | High | 10X | 0:44:58 | 25.43 |
| `MEGAHIT` | Bone | High | 10X | 0:16:59 | 2.07 |
| `metaSPAdes` | Bone | High | 10X | 1:38:39 | 19.12 |
